# Diversity at single nucleotide to pangenome scales among sulfur cycling bacteria in salt marshes

**DOI:** 10.1101/2023.06.08.544202

**Authors:** Sherlynette Pérez Castro, Elena L. Peredo, Olivia U. Mason, Joseph Vineis, Jennifer Bowen, Behzad Mortazavi, Anakha Ganesh, S. Emil Ruff, Blair G. Paul, Anne E. Giblin, Zoe G. Cardon

## Abstract

Sulfur-oxidizing and sulfate-reducing bacteria in salt marsh sediments are major controllers of ecosystem-scale carbon cycling. Cross-site comparisons of S-cycling communities are difficult given the rampant uncultured microbial diversity in sediment, yet comparisons are essential for revealing biogeographic, phylogenetic and functionally significant variation. Here, we use deep shotgun metagenomic sequencing data to construct and compare metagenome-assembled genomes (MAGs) of sulfur-cycling bacteria from Massachusetts and Alabama salt marshes that contrast in seasonality and sediment organic matter content. Samples were collected from sediments under *Sporobolus alterniflorus* and *Sporobolus pumilus* in separate MA vegetation zones, and under *Sporobolus alterniflorus* and *Juncus roemerianus* co-rooted in AL marsh. We grouped metagenomic data by plant species and site and identified 38 MAGs that included pathways for dissimilatory sulfate reduction or sulfide oxidation. Phylogenetic analyses indicated that 30 of the 38 were affiliated with uncultivated lineages. Read-mapping to MAGs showed significant differentiation of AL and MA samples, differentiation of samples taken in *S. alterniflorus* and *S. pumilus* vegetation zones in MA, but no differentiation of samples taken under *S. alterniflorus* and *J. roemerianus* that were rooted together in AL marsh. Pangenomic analyses of eight ubiquitous MAGs also detected site- and vegetation-specific genomic features, including varied sulfur-cycling operons, carbon fixation pathways, fixed single nucleotide variants, and active diversity-generating retroelements. This genetic diversity, detected at multiple scales even within uncultured groups, suggests evolutionary relationships affected by distance and local environment, and demonstrates differential microbial capacities for sulfur and carbon cycling in salt marsh sediments.

**Importance:** Salt marshes are known for their significant carbon storage capacity, and sulfur cycling is closely linked with the ecosystem-scale carbon cycling in these ecosystems. Sulfate reducers are the major decomposers in salt marsh systems, and sulfur-oxidizing bacteria remove sulfide, a toxic byproduct of sulfate reduction, supporting the productivity of marsh plants. To date, the complexity of coastal environments, heterogeneity of the rhizosphere, high microbial diversity and uncultured majority hindered our understanding of the genomic diversity of sulfur-cycling microbes in salt marshes. Here we use comparative genomics to overcome these challenges and provide an in-depth characterization of microbial diversity in salt marshes. We characterize sulfur-cycling communities across distinct sites and plant species and uncover extensive genomic diversity at the taxon level and specific genomic features present in MAGs affiliated with sulfur-cycling uncultivated lineages. Our work provides insights into the partnerships in salt marshes and a roadmap for multiscale analyses of diversity in complex biological systems.

## Introduction

Sulfur-cycling bacteria are central to the function of salt marsh ecosystems worldwide (1). While sulfate reducers catalyze the dominant decomposition pathway in salt marshes, sulfide is produced as a metabolic byproduct that can be toxic to plant roots (2–4). Microorganisms that oxidize sulfide (5), can detoxify rhizosphere sediments for plants. Thus, the metabolism of S-cycling microbes is a major contributor to high rates of marsh plant productivity (6–9). The tremendous diversity of sulfur-cycling bacteria has made cross-site comparisons of their broad genomic content difficult (10). Studying the genomic diversity of S-cycling microbes requires deeply sequenced metagenomic data to reveal gene function and phylogenetic relationships (11).

Salt marshes distributed along the eastern coast of the United States are characterized by dominant vegetation regulated by climatic, tidal and edaphic factors. Salt marsh plants are usually distributed along an elevation gradient depending on their adaptability and tolerance to reduced sediments and saline conditions. Typical species located from low to high marsh include *Sporobolus alterniflorus* (formerly *Spartina alterniflora*), *Sporobolus pumilus* (formerly *Spartina patens*), and *Juncus roemerianus*. *S. alterniflorus* is most tolerant of the frequent flooding and high salinity characterizing the intertidal zone and low elevations (12, 13). In New England, *S. pumilus* is dominant in the less frequently flooded high marsh zone (14). In Alabama, *J. roemerianus*, though often in high marsh, can mix with *S. alterniflorus* (15).

Over the past several decades, examination of microbial communities by sequencing 16S rRNA genes has identified a wide range of sulfur-cycling bacterial lineages within marsh sediments. Sulfate reducers belong to bacterial phyla such as *Desulfobacterota* (formerly *Deltaproteobacteria*), *Acidobacteriota*, and *Bacteroidota* (16, 17), while sulfur oxidizers are often found in the phyla *Proteobacteria*, *Bacteroidota*, and *Campylobacterota* (5, 18, 19), including orders *Chlorobiales*, *Chromatiales*, *Rhizobiales,* and *Rhodobacterales* (5). In a biogeographic survey of sulfate reducers in multiple East Coast salt marsh sediments, (20) found that the distribution of 16S rRNA genes varied with the environment and geographic distance, but the sulfate reductase gene did not. New metagenomic approaches are now emerging as a powerful tool to detect heterogeneity and compare sequences in multiple genes, even in highly diverse microbial populations (21, 22).

Here, we focus on the sulfur-cycling microbial communities inhabiting two contrasting salt marshes differing in type of sediment (rich in organic matter/fine), latitude (North/South), and vegetation (*S. alterniflorus* and *S. pumilus*/ *S. alterniflorus*, *J. roemerianus*). We explored the occurence of diverse metagenome-assembled genomes (MAGs) encoding dissimilatory sulfate reduction or sulfide oxidation genes (1) across sites and under different plant taxa and characterized their genomic diversity from single nucleotide to pangenome scales (Fig. 1). We investigated shared features and site- and vegetation-specific genetic diversity, providing insights into biogeographic patterns and functional diversity.

**Figure 1.**
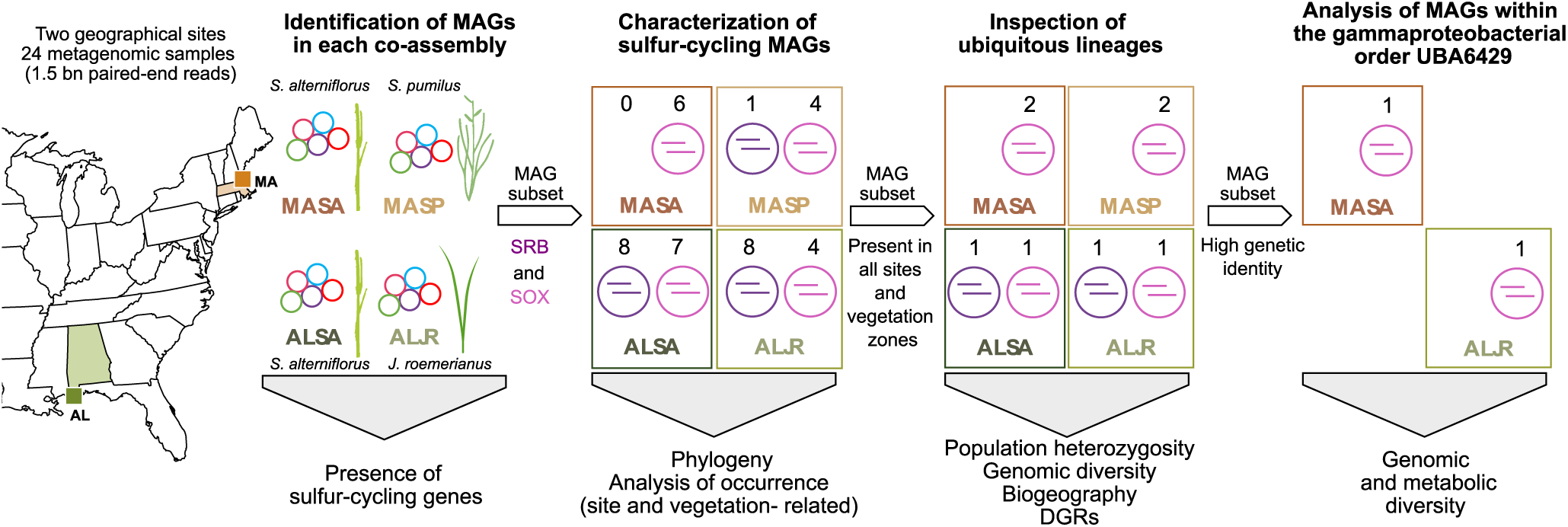
Overview of study sites and metagenomic workflow. Rhizosphere sediment samples were collected in Alabama under *J. roemerianus* (ALJR) and *S. alterniflorus* (ALSA) and in Massachusetts under *S. alterniflorus* (MASA) and *S. pumilus* (MASP). Twenty-four metagenomes yielded 38 MAGs for S-cycling bacteria (17 SRB and 21 SOX). Ubiquitous lineages were used to investigate genomic diversity.

## Results

### Dataset comparison and identification of MAGs

In our study, deep metagenomic datasets allowed us to characterize sulfur cycling microbial diversity across distinct environments and plant species. For the reconstruction of “primary MAGs” (see Methods Section), metagenomic samples were grouped according to geographical site, Alabama (AL) and Massachusetts (MA) and dominant vegetation on the collection site, *S. alterniflorus* (SA), *S. pumilus* (SP), and *J. roemerianus* (JR) (Table S1). The four co-assemblies yielded 251,034 (ALJR), 249,343 (ALSA), 70,380 (MASA), and 100,526 (MASP) contigs, with similar N50 values (7 Kb) (Table S2) and taxonomic profiles (Fig. S1). Binning resulted in 118 medium-quality MAGs (>90% completion, <5% contamination), 38 were identified by their gene content as sulfur cycling bacteria (Fig. 2, Supplemental Data 1-2), and they belong to bacterial groups traditionally associated with salt marshes including *Acidobacteriota, Bacteroidota, Desulfobacterota*, *Gemmatimonadota*, and *Proteobacteria* (Fig. 3, Table S3). Seventeen of the 38 S-cycling MAGs were sulfate reducers (including genes *sat*, *apr*AB, and *dsr*AB) belonging to uncultured lineages (labeled by alphanumeric identifiers), with five identifiable only at the family level (Fig. 3, Table S3). All but one of these SRB MAGs were assembled from AL samples. The sulfide detoxifying enzyme *sqr* (sulfide:quinone oxidoreductase) was found in 6 of the SRB MAGs (Fig 2). Twenty-one of the 38 MAGs were sulfur oxidizers with genes encoding the truncated thiosulfate oxidation system (Fig. 2); approximately equal numbers of MAGs were assembled from AL and MA samples. All were in the phylum *Proteobacteria* (Table S3). Two MAGs were affiliated with the uncultivated lineage UBA6429, which was the only lineage assembled in both Massachusetts and Alabama samples. All but three SOX MAGs harbored *sat*, *apr*AB, and *dsr*AB, likely operating in reverse, oxidizing sulfide to sulfate (Fig. 2). The majority of MAGs encoded *sqr*, but three lacked *sqr* and encoded *soxCD* genes (Fig. 2). Genes for sulfite (*soeABC*) and sulfide (*fccAB*) dehydrogenase were variably represented (Fig. 2).

**Figure 2.**
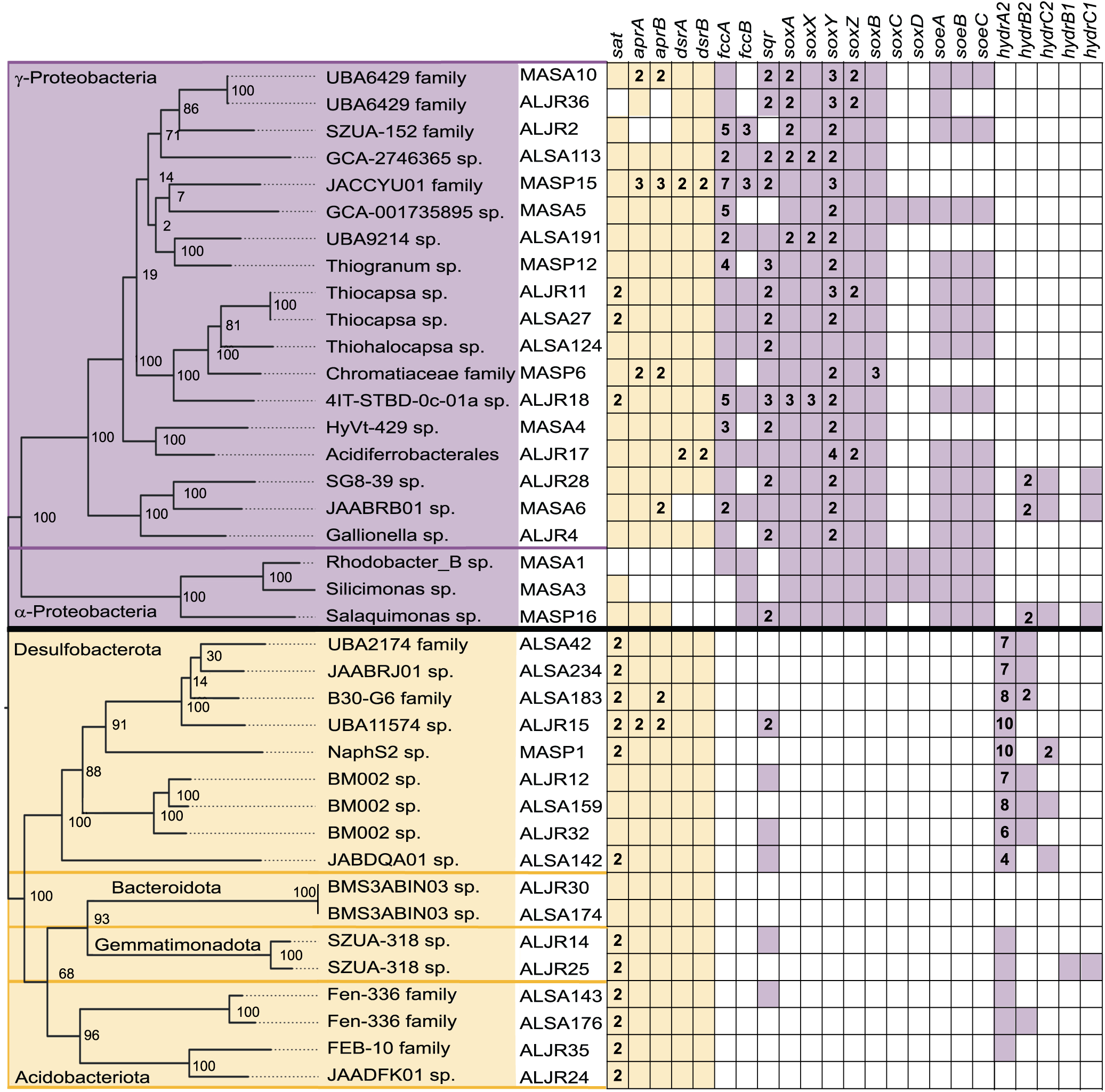
Core sulfur cycling genes detected in MAGs from Alabama and Massachusetts salt marsh samples. Key genes encoded dissimilatory sulfate reduction (purple) and sulfur oxidation (yellow). MAGs are organized by their GTDB-tk taxonomy, and named by site (AL or MA), vegetation, and bin identification number. Multiple copy genes are given as numbers.

**Figure 3.**
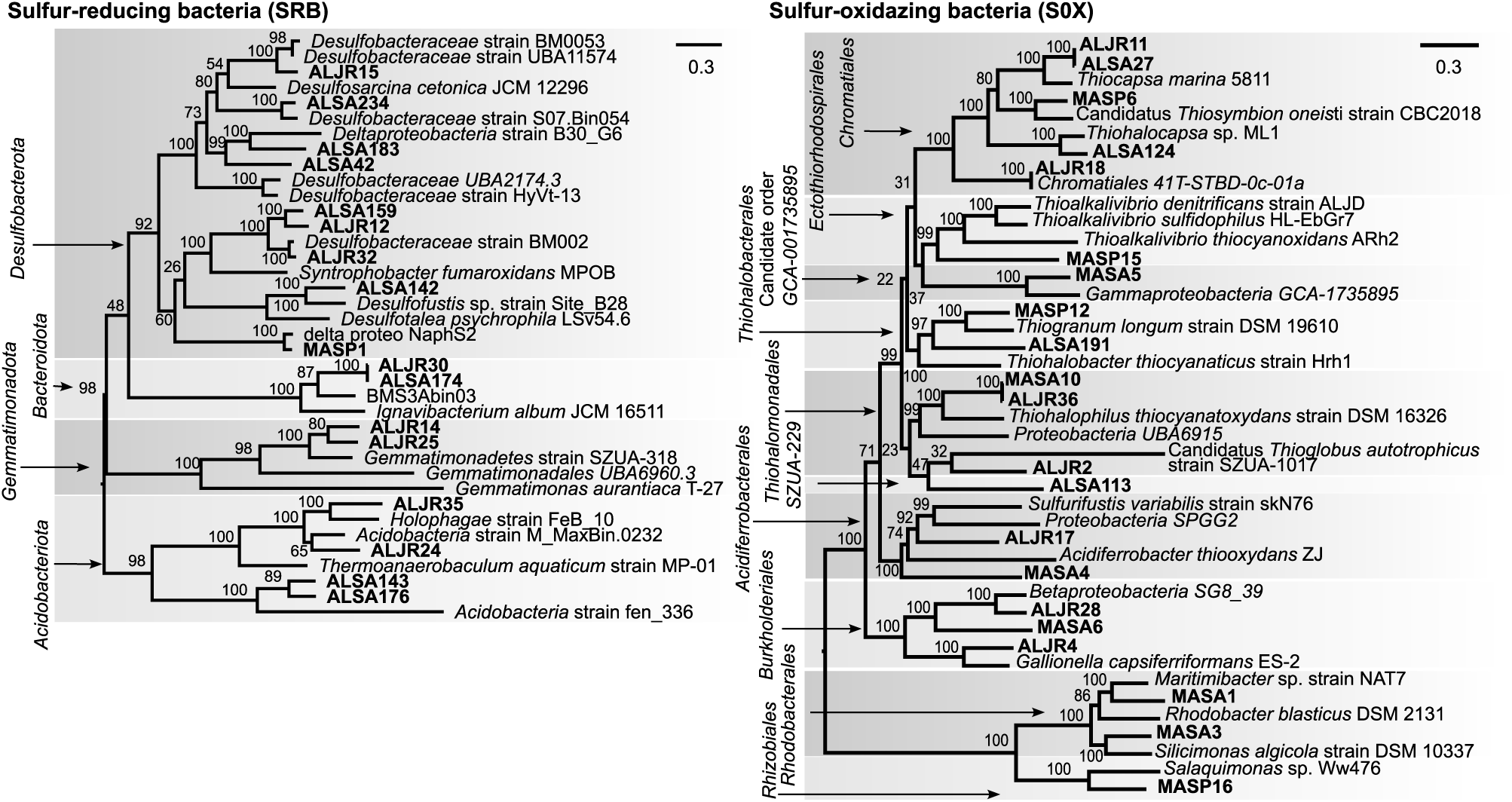
Maximum likelihood phylogenetic trees of sulfate-reducing (left) and sulfur-oxidizing (right) MAGs from Alabama and Massachusetts salt marshes, and their closest relatives, based on analysis of single-copy genes. Phyla are shown on the left of each tree.

### Distribution of sequences from the 38 MAGs across samples

To quantify and compare MAG sequence distribution across samples, we used read-mapping (Fig. 4A). On average, more reads mapped to contigs from sulfur-oxidizing “primary” MAGs (see Methods section) across all samples (average CPM 11 for SOX, vs. average CPM 3 for SRB). Also, we observed a geographical pattern related to the percentage of the MAG successfully recruiting reads. When AL metagenomic samples were mapped to AL-derived MAGs, up to 99% of contigs had CPM values different from 0. For samples collected in MA, this was reduced to 92% of contigs in samples from sediments dominated by *S. alterniflorus* and further reduced to ~75% samples collected by *S. patens*. We did not observe such a pronounced pattern in MA-derived MAGs as values ranged from 99% to 92-95% (Supplemental Data 3). For each primary MAG, patterns of read recruitment to individual contigs can be found in the supplementary results (Fig. S3-S5).

**Figure 4.**
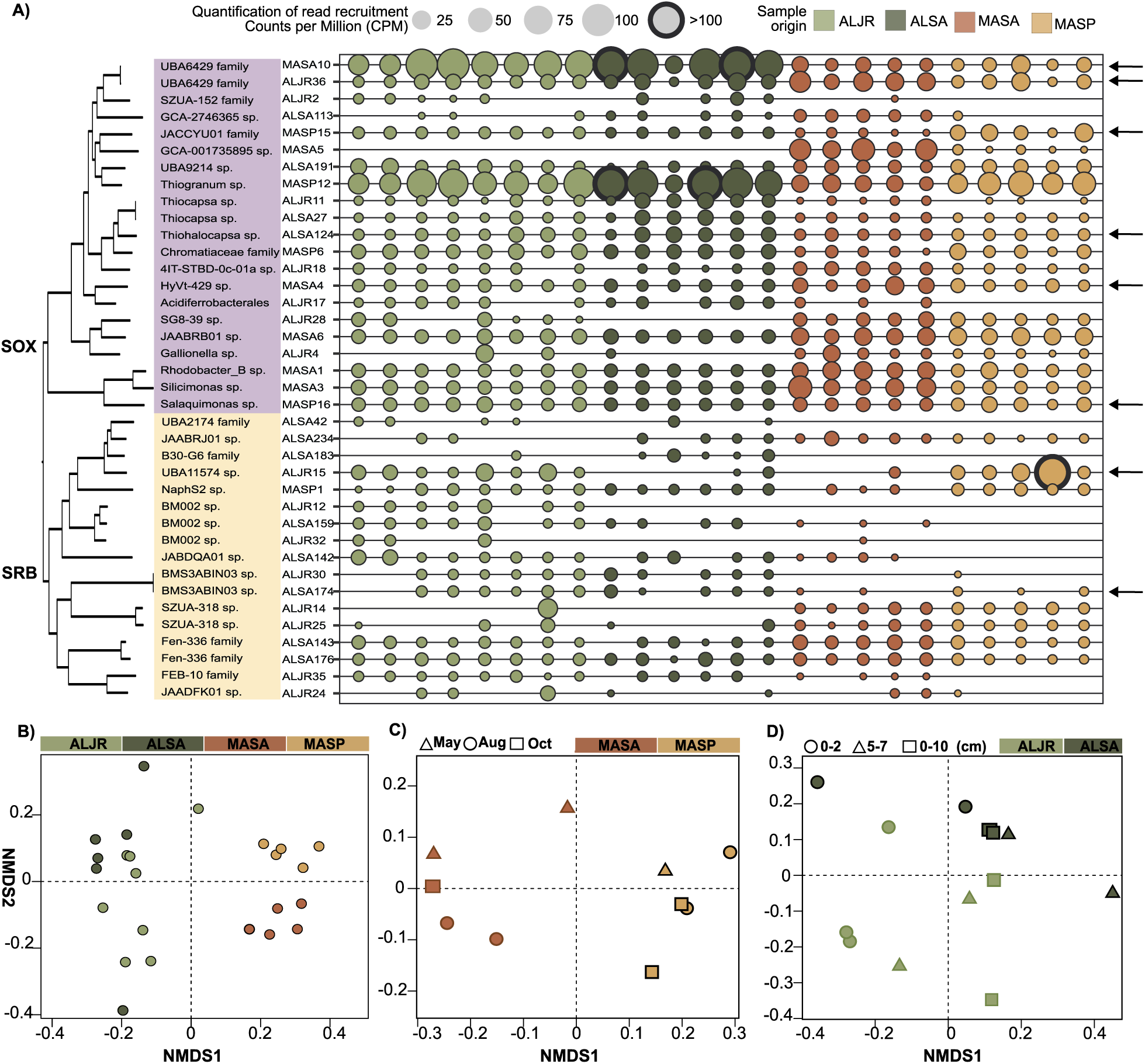
Distribution of sulfur cycling primary MAG sequences across sites and vegetation zones. **A)** Quantification of read-mapping from AL and MA samples to primary MAG contigs, with coverage standardized by library size and contig length and expressed as counts per million reads (CPM). Circle size reflects averaged CPM across all contigs for each MAG. **B-D)**Non-metric multidimensional scaling (NMDS) analyses comparing MAGs distribution as expressed as CPM in: **B)** Alabama and Massachusetts salt marshes, **C)** Massachusetts salt marshes by month, and **D)** Alabama salt marshes by depth.

Various site- and vegetation-specific patterns emerged. For example, the presence of MASA5 (GCA-1735895 sp.) was only detected in MA samples and was enriched in sediments inhabited by *S. alterniflorus* (Fig. 4A). Among sulfate reducers, ALSA176 (*Acidobacteriota* Fen-336 family) was the only MAG which recruited reads in all samples (average 6 CPM, Fig. 4A), and closely-related ALSA143 (also in the Fen-336 family) recruited reads from all but one sample (Fig. 4A). Sequences from BM002 MAGs (related to *Syntrophobacteria*) were poorly represented in MA samples vegetated with *S. pumilus* (Fig. 4A).

Such differences in MAG abundance were revealed in patterns by NMDS analysis based on the read recruitment values (CPMs). This analysis uncovered separation by site (MA or AL; Fig. 4A) and vegetation within site (Fig. 4B), by the month that the sample was collected (MA only, Fig. 4C) and by the depth of sediment (AL-only, Fig. 4D). PerMANOVA post-hoc comparisons detected significant differences between MASP and MASA samples when compared to each other and to ALJR (p = 0.01) and ALSA (p = 0.01, Fig. 4A), but no significant differences were found between ALJR and ALSA samples (p = 0.15, Fig. 4A). No significant patterns were detected by sampling month in MA (Fig. 4B). In AL (Fig. 4D), 0-2 cm depth samples were significantly different from 5-7 cm and 0-10 cm samples (p = 0.045).

The same read recruitment approach was applied to more sparsely sampled metagenomic data from unvegetated creek bed sediment samples, revealing for several primary MAGs clear variation in read recruitment as a function of salinity in two tidal creeks at PIE-LTER (Fig. S6, Supplemental Data 4).

### Contig-based analysis of genomic variability

Genomic coverage and genetic diversity in eight primary MAGs (identified by arrows, Fig. 4A) that occurred in all sampling locations (Fig. S7) were analyzed further using the Anvi’o profiling pipeline (Supplemental Data 5). For all MAGs (Fig. 5A), we observed a relative uniformity in GC content (inner ring) and mean read coverage (colored rings) across the contigs forming each bin. Overall, read recruitment to contigs was influenced by, but was not solely governed by sample sequencing depth (see grey histograms Fig. 5A, Fig. S8-S15). While sequence quality, contig length, assembly method, genetic diversity, and experimental design can influence read recruitment, here we observe an effect on the geographical origin of the sample. Each MAG was better represented in the pool of samples originally used for its assembly and was similarly abundant in the samples of a given site (Fig. 5A). This was not the case for the sediments inhabited by *S. pumilus,* where a MAG could be highly abundant just in one sample. This seems to suggest that in this area microorganisms might have a greater spatial heterogeneity than those areas inhabited by *S. alterniflorus* and *J. roemerianus* (Table S4).

**Figure 5.**
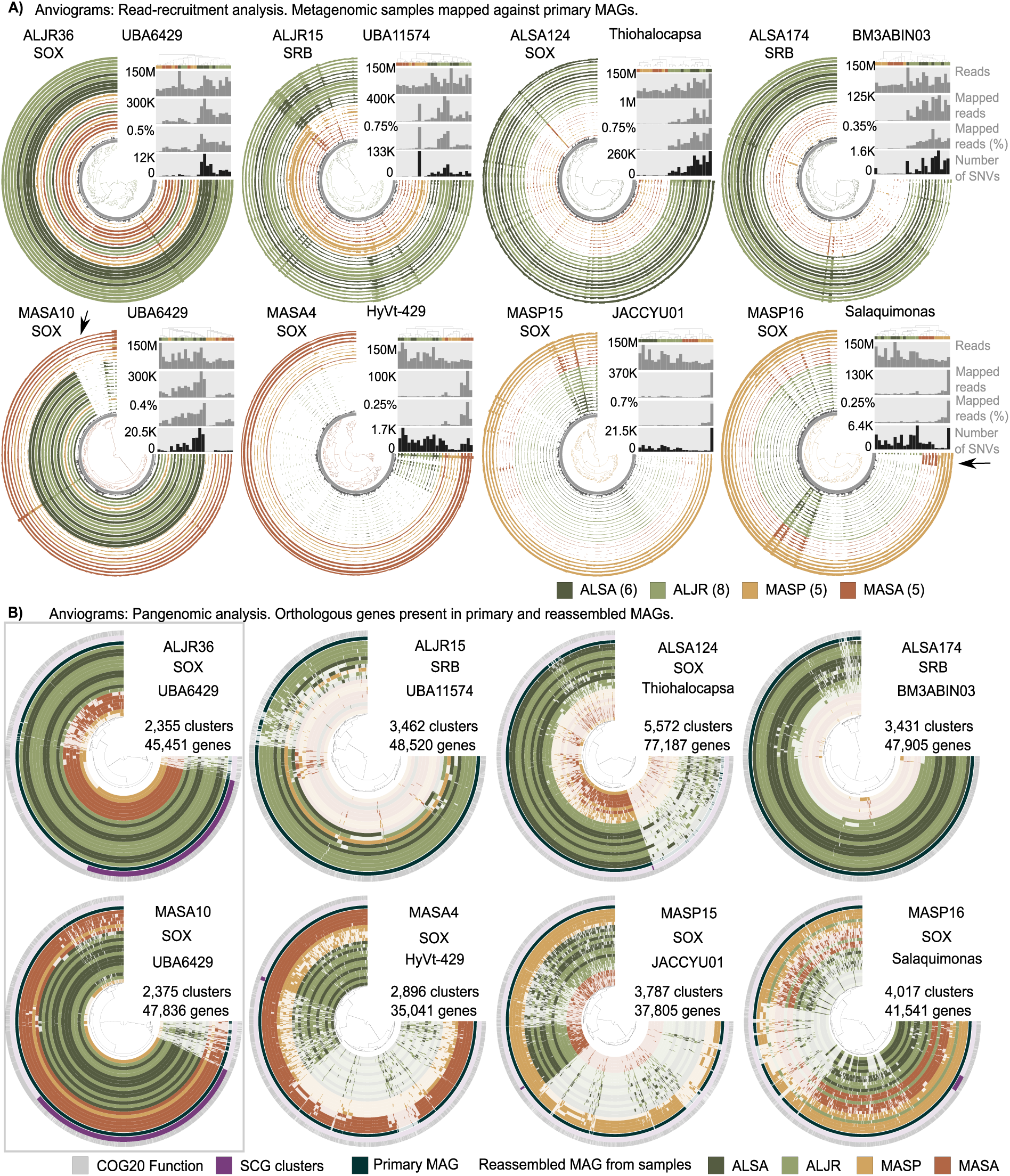
**A).** Anvi’o coverage profiles of eight selected sulfur-cycling bacteria MAGs. Outer cycles represent the contig coverage (as mean coverage) of each of the 24 salt-marsh metagenomic samples included in this analysis. Inner rings represent GC content (dark gray) and contig length (gray). Contigs are clustered (inner tree) based on the sequence composition and differential coverage using Euclidean distance and Ward hierarchical clustering method. Sample order (rings) was determined using a clustering method based on the mean coverage and each ring is color-coded according to site and vegetation type. Bar plots represent (top to bottom) the total number of reads in each library, the total number of reads mapped to each respective MAG, the percentage of mapped reads, and the total number of Single Nucleotide Variants (SNV). **B)** Anvi’o pangenomic analysis of eight sulfur-cycling bacteria MAGs. In each ring, each vertical line represents a gene. Gene order was determined using Euclidean distance and Ward clustering method based on presence/absence of a gene cluster across MAGs. Annotation based on COG 2020 is indicated in gray and single-copy core genes in purple. Primary MAG is shown in dark teal blue. Each of the remaining circles represents a reference-guided reassembled MAG from each of the salt-mash metagenomic samples included in this analysis. Samples are color-coded according to geographical site and vegetation.

Patterns of recruitment varied across primary MAGs. For example, UBA6429 MASA10 contained an 80.4 Kbp region, encoding 86 genes, that was only detected in samples collected in Massachusetts under *S. alterniflorus* (Fig. 5A, arrow). This MASA-exclusive region encodes duplicated copies of genes involved in the biosynthesis of molybdenum cofactor (Moco), and it was rich in transport genes (Supplemental Data 6). Moco is essential for the activity of molybdoenzymes involved in the metabolism of sulfur compounds such as the conversion of toxic sulfite to sulfate. *Salaquimonas* sp. MASP16 included an 18.5 Kb region that was only present in the samples collected in Massachusetts, in either *S. alterniflorus* or *S. pumilus* vegetation zones (Fig. 5A, arrow). This region encoded 22 genes, including a second set of genes of the phospholipid transporter system MlaFEDB and the genes *arcC* and *arcA* involved in arginine biosynthesis (Supplemental Data 6).

We investigated single nucleotide variants (SNVs) by quantifying the divergence of recruited reads from the primary MAGs (Fig. 5A, Supplemental Data 5). For each of the four AL-derived MAGs, and for UBA6429 MASA10, the number of SNVs (black histograms) was correlated with the number of mapped reads (grey histograms) across all 24 samples (Fig. 5A, Fig. S7, and Supplemental Data 5). But for primary MAG MASA4, AL samples had far more SNVs per mapped read than did MA samples (Fig. 5A). Similar global variation was observed in samples MASP15 and MASP16, all AL samples, and MA samples under *S. alterniflorus*, wherein all had far more SNVs per mapped read than the MA *S. pumilus* samples (Supplemental Data 5).

### Gene-based analysis of sample-specific reassembled genomes

Using the same 8 primary MAGs as references, we reassembled sample-specific MAGs where possible (Table S5). Completeness varied widely and was highest for samples that had originally been coalesced to generate the primary MAG co-assembly. We used Anvi’o pangenomic analysis (Fig. 5B, Fig. S8-S15) to annotate the genomes and identify clusters of homologous genes. Our results indicate that the geographical origin of the primary MAG (AL or MA) determined the reassembly success. In ALJR15, ALSA124, and ALSA174, initially identified in co-assemblies of AL samples, we observed a high degree of completeness among Alabama-reassembled MAGs (55-92%) while we observed limited reassembly success among the MA samples. An extreme case was MAG ALSA174 where Massachusetts-derived MAGs completeness was calculated at 1-3%. For all the MA MAGs and a single AL one (ALJR36), we successfully reassembled sample-specific MAGs, whether derived from MA (55-65%) or AL (75%) samples (Supplemental Data 7). For UBA6429 MASA10 and ALJR36, sample-specific reassemblies from all sites shared a very large proportion of the gene clusters found in the original primary MAG. Single-copy core genes were readily identified as 42% and 33%, respectively, of the total number of genes (Fig. 5B, Fig. S8 and S12). For the other 6 MAGs, SCGs were sparse or absent (Fig. 5B, Fig. S9-S15).

### Hot spots of variability in sample-specific reassembled MAGs

The close phylogenetic relationship of UBA6429 ALJR36 and MASA10 (Fig. 3, ANI: 99.43%) and coexistence across sites (Fig. 4, 5B) provided a unique opportunity for the study of genetic diversity and microevolution that might be linked to metabolic capabilities. Previous analyses (described above) generated SNV counts for each of the 24 metagenomic samples (Fig. 5A, grey histograms). To determine the average number of SNVs per kilobase, we coupled the analysis of SNVs in the sample contigs with gene annotation data (black bars, Fig. 6A). We show that SNVs remained stable across the MAGs, with the exception of a handful of ORFs with outlier values (Fig. 6A, arrows). Notably high variability was observed within prophages and transposons (COG5361), rRNA, tetratricopeptide repeats responsible for protein-protein interactions or assembly of multiprotein complexes (COG0457), and the large subunit of a Rubisco-like protein (COG1850). As observed in the previous SNV analysis (Fig. 5A, Supplemental Data 5), SNVs were more abundant across the full genomes of Alabama-specific MAGs that had more mapped reads (Fig. 6A, dense black bars across full genomes.)

**Figure 6.**
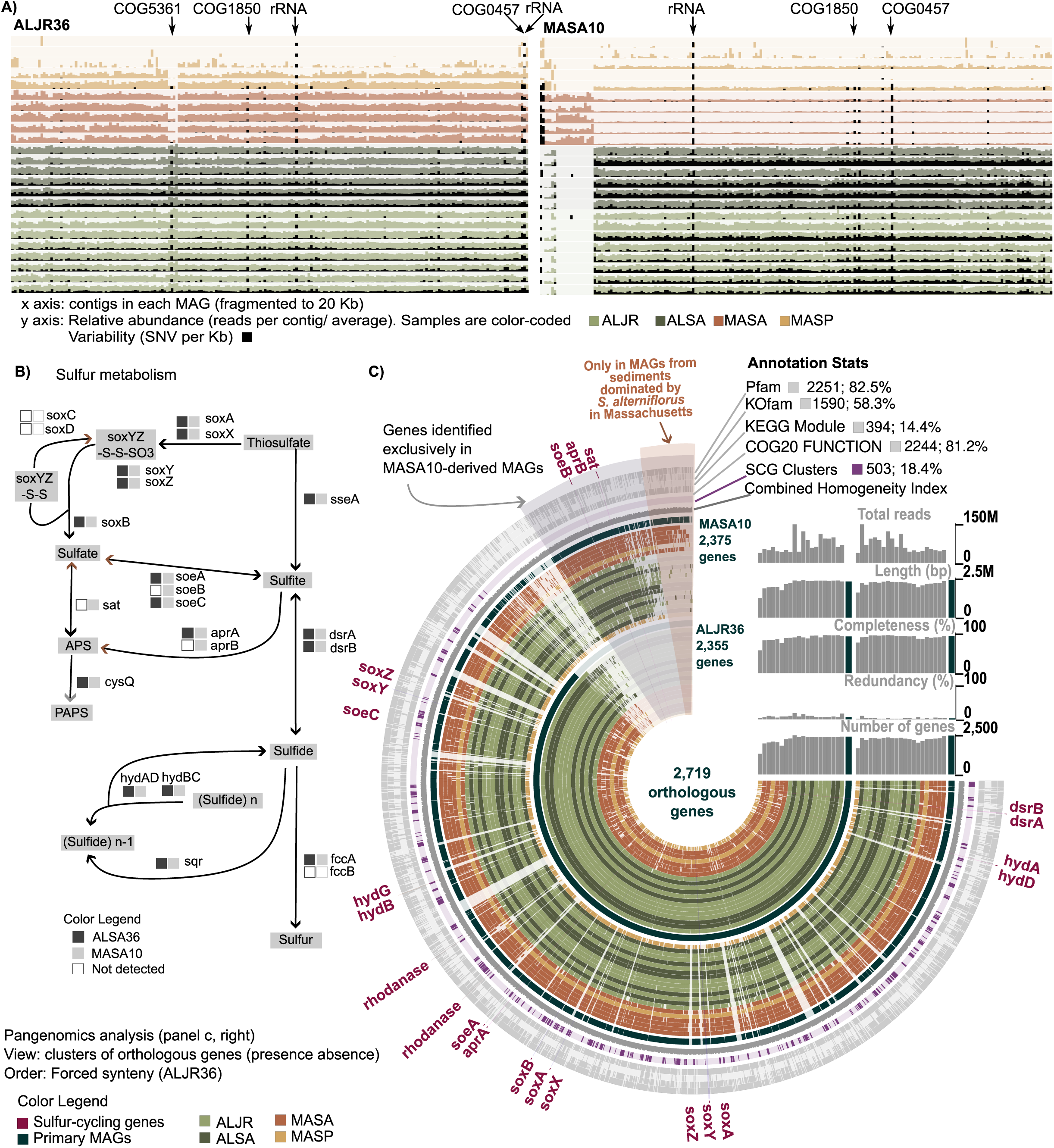
Comparative analysis of genomic and metabolic diversity of two closely-related sulfur-oxidizing MAGs (*Thiohalomonadales*) **A)** Distribution of genetic variability represented as SNV per kilobase (black bars) and calculated for each metagenomic sample. Hotspots of variability include COG5361(Mobilome), COG1850 (Rubisco-like protein), COG0457 (Tetratricopeptide (TPR) repeat), and rRNA. **B)** Diagram of sulfur metabolism pathways highlighting the differences in gene content between sample-specific MAGs from the ALJ36 and MASA10 primary MAG groups (adapted from Kegg map00920) **C)** Pangenomic analysis of metagenomes from group ALJR36 and MASA10. Primary MAGs are indicated by the dark ring and sample-specific re-assembled MAGs are color-coded by site and vegetation. Genes clusters are ordered with forced synteny to ALJR36 to highlight the genes exclusively found in the MASA10 group. Annotations, single-copy core genes, and the combined homogeneity index are shown in outer rings.

### Pangenomic analysis

Sample-specific genomes reassembled using as a guide the closely related UBA6429 MASA10 and ALRJ36 primary MAGs were grouped by site (AL or MA) for pangenomic analysis within Anvi’o. Three MA *S. pumilus* samples were excluded because the completeness of the reassembled genomes was <50% (Supplemental Data 7). All samples were compared in terms of presence/absence or orthologous genes, revealing a shared subset of the genomes (Fig. 6C). Of the 2,719 genes identified in the pangenomic analysis, 2,375 were present in the MASA10 group and 2,355 in the ALJR36 group (Fig. 6C). Functional characterization was relatively complete with 81.5% of the genes identified using COG20, and 82.5% with Pfam. Moreover, 58.3% of the genes were assigned Kegg numbers (grey outer rings, Fig. 6C). A repertoire of 503 SCG were identified across the two groups of metagenomes (purple ring, Fig. 6C). The order of ORFs around the rings in Fig. 6C (which is not related to the order in genomes) visually emphasizes genes found only in re-assembled MAGs from MA samples taken under *S. alterniflorus* (labeled MASA), and genes found only in the family of sample-specific MAGs reassembled with primary MAG MASA10 as a guide (labeled MASA10-specific genes).

Despite the large similarity among both groups of genomes, 284 genes were present only in ALJR36 reassembled metagenomes and 292 genes in the MASA10 group. Within the MASA10 group, 80 genes were exclusively identified in samples collected in Massachusetts in intertidal sediments inhabited by *S. alterniflorus* (Fig. 6C, sector at top of anviogram). Functional enrichment analysis indicated that 11 COG categories were statistically enriched in MAGs from the group ALJR36 and 8 in MAGs from MASA10 (Table S6, Supplemental Data 8). These COG categories include inorganic ion, carbohydrate, and amino acid transport. KEGG enrichment analysis indicated different metabolic capabilities between the MASA10 and ALJR36 groups of metagenomes. Notably, phosphate acetyltransferase-acetate kinase and the reductive citrate cycle (Arnon-Buchanan cycle) pathways were enriched in the ALJR36 group (Table S7, Fig. S17). This was caused by the presence of the genes encoding the enzymes acetate kinase (*AckA*) and aconitate hydratase (*AcnB*) in the ALJR36 group, and the lack of the same genes in the MASA10 group. The MASA10 group was significantly enriched for sulfur-related KEGG modules. Particularly distinguishing are that the ALJR36 group MAGs are missing complete genes encoding subunit B of the sulfohydrogenase *soe*ABC complex, and several key genes (*sat, aprB*) in sulfur cycling pathways (Fig. 2, Fig. 6C). Although DRAM did not find *soeC* in ALJR36 (Fig. 2), we were able to identify an incomplete copy using Anvi’o (Fig. 6C).

### Phylogenetic analysis indicates structure in genomic variability

In the analyses described above, nucleotide variation and entropy analyses captured the heterogeneity of the samples, while pangenomic analysis suggested metabolic differences between closely-related MAGs. Using reassembled MAGs, we conducted phylogenetic analyses of single-copy genes and protein-coding marker genes (Supplemental Data 9), capitalizing on fixed changes at the sequence level (Fig. 5, Fig. 7A) indicative of divergent populations.

**Figure 7.**
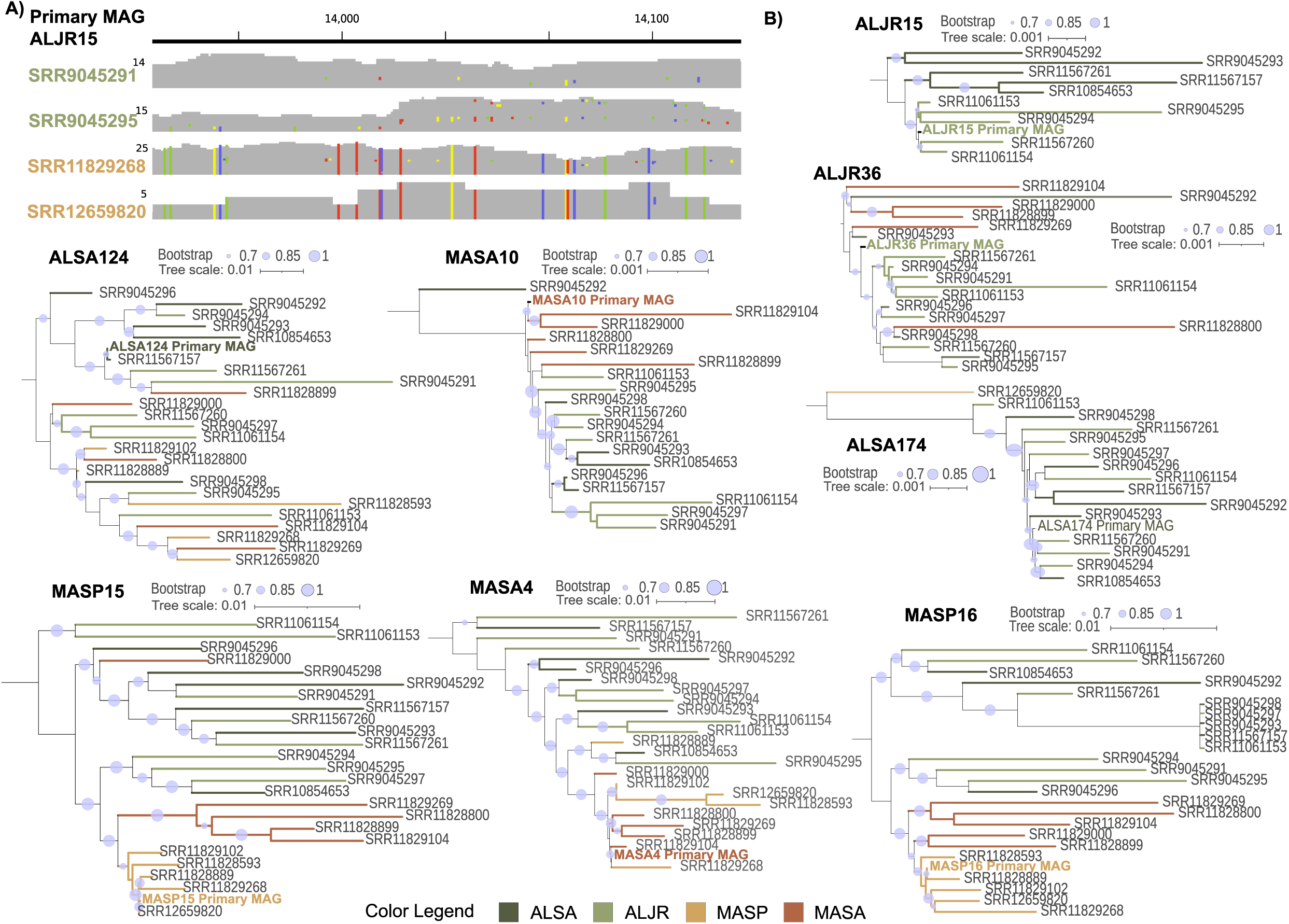
Biogeographic and vegetation-specific distributions of genomic variability. **A)** Example aggregated view of metagenomic reads from different samples mapped to the MAG ALJR15. Samples collected in Alabama matched the reference while those collected in Massachusetts in sediments dominated by *S. pumilus* displayed numerous fixed mismatches. Mismatches are indicated with colors: green, T; yellow, G; blue, C; red, A. **B)** Trees depicting phylogenetic relationships among reassembled sample-specific metagenomes. Trees were built using single-copy genes, housekeeping genes, and sulfur-cycling genes. Tree branches are color-coded according to site and vegetation.

For the four groups of sample-specific MAGs whose reassembly had been guided by MA primary MAGs (MASA10, MAS15, MASP16, MASA4), sample-specific MAGs largely fell in well-supported Massachusetts (red-brown) and Alabama (green-olive) clades (Fig. 7B). MAGs from samples from the two MA vegetation zones (*S. alterniflorus* and *S. pumilus*) also tended to fall in separate clades, possibly either because the vegetation was different, the inundation intensity was different, or both.

MAGs from Alabama samples in group ALJR15 clearly segregated by plant type (Fig. 7B), though the plants occurred together under the same inundation regime. Sample-specific MAGs in the other three AL primary MAG groups did not segregate by vegetation or site (Fig. 7B).

### Diversity-Generating Retroelements (DGRs) are present and active

Finally, we investigated the possibility that DGRs contributed to the diversification of the sample-specific MAGs. DGRs use an error-prone reverse transcriptase to generate variability in specific target genes (Fig. 8A). Two sulfur-oxidizing MAGs (ALSA124 and MASP15, Fig. 2), both in the *Gammaproteobacteria*, encoded full DGRs. (Beyond the 38 MAGs otherwise targeted in this paper, DGRs were also found in two other *Gammaproteobacteria* in the original 118 MAGs – ALJR7 and MASP22.) ALSA124 and ALJR7 (*Chromatiales*) contain a single DGR and a single target gene. MASP15 and MASP22 (UBA4575) encoded two and three DGRs, respectively, one (in MASP22) with multiple target genes (Fig. 8B).

**Figure 8.**
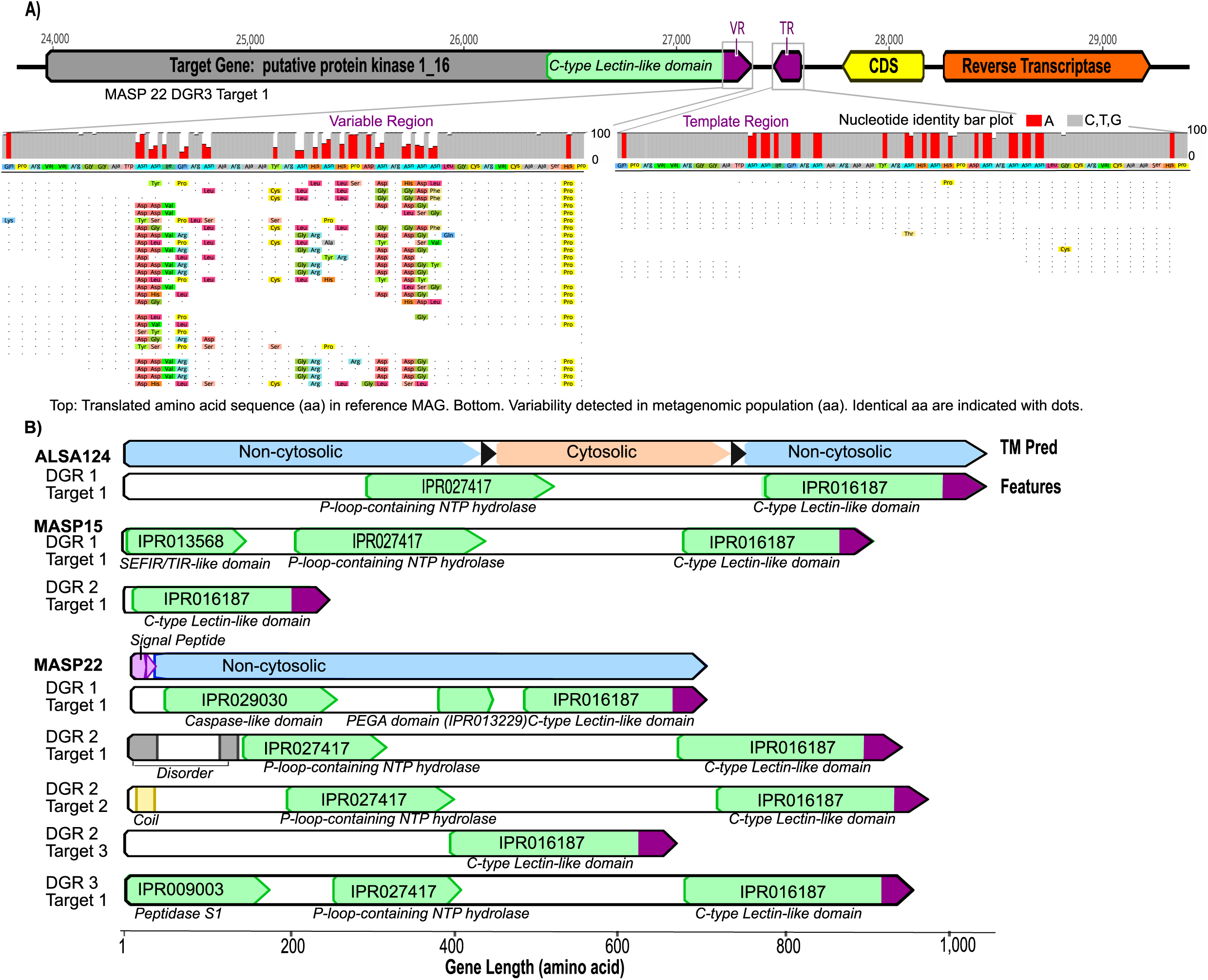
**A)** Structure of an example DGR identified in MASP22. Diversification of this DGR in this metagenomic sample is indicated by the accumulation of in-frame mutations in the variable region (VR) among the aligned reads. The template region (TR) is conserved both at the nucleotide and amino acid levels. In the bar graphs, the bar size is proportional to the degree of conservation of the nucleotide position. Adenine positions in the VR and TR regions are indicated in red. In the recruited reads, identical residues are indicated with dots, and changes are color coded. **B)** Structure and domain composition of the target genes containing variable regions (in purple) of the DGRs.

In all target genes, the variable region of the DGR was encoded in a C-type lectin-like domain. This domain is found in a diverse group of proteins, some of which are involved in protein-protein, protein-lipid, or protein-nucleic acid interactions (23). Energy-related domains such as NTP hydrolases were found in 5 out of the 8 target genes. The most obvious instance of active DGR activity was detected in MASP22. Adenines in the template region were highly conserved (Fig. 8B). In the DGR target gene, 15 variable codons correspond to template (TR) adenines, which have the potential for DGR-directed non-synonymous substitutions, while not directly recoding to a stop codon (Fig. 8A). These correspond to nine asparagine (AAY) residues, one isoleucine (ATC), one glutamine (CAA), one tyrosine (TAT), two histidines (CAY), and one aspartic acid (GAT) encoded in TR. Taken together, the DGR target region can accommodate up to ~1.97×10^14 amino acid variants. In the mapped reads to this MAG, we observed recoding to between 2-8 amino acids at each DGR-variable position. This observation suggests that a diversity of between ~378 million and ~2×10^14^ variants could be randomly rewritten through adenine mutagenesis of this single gene.

## Discussion

Sulfur-cycling organisms are taxonomically diverse and very abundant in salt marsh sediments. Deep metagenomic sampling in AL and MA marsh sediments and comparative genomic analysis enabled the characterization of microbial genetic and genomic patterns from single nucleotide to pangenome scales.

### Genomic patterns reveal the potential contribution of taxa across environments

We found SRB MAGs hosting genes for dissimilatory sulfate reduction (*sat*, *aprAB*, *dsrAB*) within multiple phyla, with *Desulfobacterota* dominant (53% of MAGs) as previously observed in Massachusetts (16), Mississippi (19), and Georgia (24) salt marshes. Other SRB MAGs belonged to groups (*Acidobacteriota*, *Bacteroidota*, and *Gemmatimonadota*) well recognized to contain sulfate reducers (10, 19, 25–28). All but one SRB primary MAG were co-assembled from AL samples, perhaps because SRBs are so diverse and the sequencing depth at AL was double that in MA. All sulfate-reducing MAGs belonged to genera without cultivated representatives, underlining the capacity of metagenomic analyses to further expand the recognized diversity and genomic potential of sulfate reducers (17, 29).

The taxonomic distribution of SOX MAGs containing the truncated thiosulfate oxidation system (*soxAXYZB*) was simpler, falling only within the phylum *Proteobacteria*, with *Gammaproteobacteria* representing 86% of the genomes. In contrast to the SRBs, 38% of the identified sulfur oxidizers belonged to genera with cultivated representatives. Consistent with previous work (5, 30), *Chromatiales* were well represented overall, and read-mapping to ALSA27 (a *Thiocapsa* sp.) indicated significantly greater abundance under *S. alterniflorus* in AL compared to any other sample set, including AL *J. roemerianus* (30). Variations in gene content associated with distinct metabolic steps of sulfur oxidation are well documented in the literature (1, 31, 32). For example, sulfur oxidizers with the truncated *sox* system (lacking *sox*CD) often have *soe*ABC (e.g., (31)), and this pattern was observed for 12 of 21 SOX MAGs.

Clear patterns emerged in read recruitment from the 24 samples to primary SOX and SRB MAGs. These patterns indicate far more sparse mapping to SRB MAGs no matter the group from which they were co-assembled, possibly reflecting, again, the greater diversity of SRBs in marsh sediment. NMDS analysis of the read-mapping metric CPM shows strong overall separation between MA and AL samples, no effect of sampling month (variable in MA only), and a distinctive surface (0-2 cm) community in AL. Using phospholipid fatty acid analysis, (33) similarly found that salt marsh microbial communities sampled at the Wadden Sea were influenced by sediment depth but not the month (April, July, October) of sampling. In a 16S rRNA-based comparison of sediment microbial communities from MA to South Carolina, (34) also detected strong north-south differentiation.

NMDS analysis also clearly shows separation within MA of samples taken under *S. pumilus* and *S. alterniflorus*, which may reflect the influence of the plants themselves, the tidal regime (inundation frequency and depth being low for *S. pumilus* and high for *S. alterniflorus*), or both. Data from (35) show that in the greenhouse, rhizosphere microbial communities vary even with different MA genotypes of *S. alterniflorus*, but once transplanted back into the marsh, plant-specific differences disappeared and microbial communities correlated instead with local sediment properties. (33) also demonstrated that salt marsh zones defined by flooding regimes had distinct microbial communities. The strong influence of the local sediment tidal regime is further suggested by the lack of difference between AL samples taken under *S. alterniflorus* and *J. roemerianus*, which were both rooted in the same area with the same tidal regime.

### Biogeography and potential microdiversification

Local variants were detected among sample-specific MAGs reassembled with guidance from the primary MAGs. Already we had seen that reads mapping to some groups of contigs appeared only in samples from particular locations (e.g., MA samples in the MASP16 group) or even only in samples under one vegetation/tidal regime in MA (e.g., MASA10). Not only did AL and MA S-cycling MAGs separate cleanly on NMDS plots, but also the phylogenetic trees built from sample-specific MAGs showed consistent separation of MA and AL samples (except when orthologous genes in MA sample-specific MAGs were rare, as in ALJR15, ALSA124, and ALSA 174. These geographic patterns are consistent with the possibility of distance and climate differences working as a geographical barrier and contributing to allopatric differentiation (when a species separates into two or more species, due to geographic isolation and differences in environmental conditions) of microbial communities and/or the emergence of subspecies/strains (36, 37).

UBA6429 MAGs, ALJR36, and MASA10, had particularly high similarity to one another (ANI >99%), high read recruitment across all sample-specific reassembled MAGs, strong representation of SCG clusters, and yet notable differences in the presence/absence of orthologous genes. The contrasts in genomic content in these two sets of MAGs could point to metabolic differentiation, or even hint at adaptation to different environments or niches. Functional enrichment analysis identified that the ALJR36 group of sample-specific MAGs were enriched in key enzymes of the reductive citrate cycle (Arnon-Buchanan cycle (38). This CO_2_ fixation cycle is found in anaerobic and microaerobic autotrophic bacteria (39) and anoxygenic phototrophs (40). When water-saturated, salt marsh sediments are anaerobic. Particularly conducive conditions for the reductive citrate cycle’s operation would be provided by the low permeability, high clay content sediment in AL marsh near Dauphin Island airport (41). In contrast, functional enrichment analysis identified that the MASA10 group of sample-specific MAGs were enriched in a diverse genetic repertoire of genes involved in assimilatory and dissimilatory sulfur cycling including the full set of *soeABC*, *sat*, and *aprAB* genes. ALJR36 MAGs lack *aprB*, *soeB*, and *sat* genes, potentially significantly altering transformations among sulfate, sulfite, and APS. Because these are not isolated organisms, it is not possible to test unique physiologies. There are no cultivated members of the UBA6429 family of sulfur oxidizers. But, if these MAGs do accurately reflect the genetic repertoire of microbes abundant enough to have supported MAG assembly, such differences in basic gene content could reflect genomic microdiversification contributing to the microbes’ coexistence despite their sharing large portions of their genomes (42, 43).

Finally, maintaining genetic variation is important for species stability and survival (44, 45). We found DGRs encoded in the MAGs and evidence of their dynamism in the form of extraordinarily elevated non-synonymous substitutions, corresponding to adenine-specific variation of VR encoded in the target genes. Though the identity of the target genes remains unknown in these MAGs, in all target genes the variable region of the DGR was encoded in a C-type lectin-like domain, known to be involved in protein-protein, protein-lipid or protein-nucleic acid interactions (23).

These results, from single nucleotide to pangenome scales, demonstrate that a tremendous range of genetic information available in metagenomic datasets, even those from microbially diverse systems such as salt marsh sediments, can be harnessed for analysis of biogeographic and biotic patterns in targeted taxa.

## Methods

### Data set collection

We compared the sulfur-cycling bacteria in sediments of two salt marshes, representing Northern (Massachusetts, MA) and Southern sites (Alabama, AL). These sites also differed in the type of sediment with MA area being carbon rich and AL formed by fine, sandy sediments. Metagenomic datasets were retrieved in July 2020 from the Genomes Online Database (https://gold.jgi.doe.gov/) and had been sequenced by the Joint Genome Institute using Illumina NovaSeq as sequencing technology and all samples consisted of paired-end, non-overlapping, 150 nt read length. The 24 metagenomes analyzed included samples taken from rooted sediment under common dominant plant species in their native range, and all of them have a sequencing depth of > 30 million reads per sample (Table S1). The Alabama dataset (Gs0135940 P.I. Olivia Mason), included rhizosphere sediment samples collected under *J. roemerianus* (n=8, total reads ~ 578 M, abbreviated as ALJR) and *S. alterniflorus* (n=6, total reads ~ 514 M, abbreviated as ALSA) on Dauphin Island (15). Two to three samples were taken at various depths (all within 0-10 cm) in May during 2015, 2016, and 2017. The elevation of *J. roemerianus* and *S. alterniflorus* patches do not differ and the marsh is flooded on every high tide. The Massachusetts dataset (Gs0142363, P.I. Jennifer Bowen) included samples taken during May, August, and October from low marsh sediments (0-5 cm depth) under *S. alterniflorus* (n=5, total reads ~ 231 M, abbreviated as MASA) and from high marsh sediments under *S. pumilus* (n=5, total reads ~ 208 M, abbreviated as MASP) in the Plum Island Long Term Ecological Research (PIE-LTER) site.

### Metagenomic assembly, binning and analyses

Samples were grouped by site and vegetation for increased sequencing depth during the initial reconstruction of “primary MAGs”. Reads were assembled using MEGAHIT (version 1.2.9) (46) and the quality of the four resultant assemblies was evaluated using MetaQUAST (47) (Table S2). Taxonomic classification of the contigs forming each co-assemblies was performed using Kraken2 (48). Contigs were assigned into bins using MetaWRAP (49) processing modules with initial extraction using MaxBin2 (50), metaBAT2 (51), and CONCOCT (52) and finalized using the bin refinement module. Completeness and contamination of each MAG were evaluated with CheckM (53). Quality was reported based on Bowers et al. (54). MAGs were annotated using Distilled and Refined Annotation of Metabolism (DRAM) (55). Sulfur-cycling pathways were manually curated by searching key sulfur genes in the raw (Supplemental Data 1) and metabolism (Supplemental Data 2) output files provided by DRAM. MAGs corresponding to sulfur-cycling bacteria were identified by the presence of dissimilatory reduction of sulfate to sulfite pathway (*sat*, *apr*AB, and *dsr*AB) or the truncated thiosulfate oxidation system (*sox*AX, *sox*YZ, *sox*B) (56). The presence of sulfide and sulfite oxidation genes including flavocytochrome c complex (*fcc*AB), sulfide:quinone oxidoreductase (*sqr*), and the sulfite:quinone oxidoreductase complex (*soe*ABC) was also investigated. Taxonomy and closest phylogenetic neighbors were assigned using GTDB-tk (v1.5.0) (57) and average nucleotide identity (ANI) computed by EZBioCloud OrthoANIu Calculator (58). A maximum likelihood phylogenetic tree of genomes was generated using the Bacterial and Viral Bioinformatic Resource Center (59, 60).

### Distribution of 38 primary MAGs’ sequences across samples

The metaWRAP Quant_bins module (49) with Salmon (61) was used to recruit reads from each sample to the contigs forming each of the 38 primary MAGs. Coverage values are standardized by library size and contig length, then expressed as counts per million (CPM). CPM metrics for entire MAGs were calculated as the average CPM across individual contigs. Results below 2 CPM were filtered out. For each MAG, average CPM values were compared to contig-specific CPM. Additionally, to determine MAGs occurrence in less deeply sequenced data, metagenomic reads from unvegetated salt marsh sediments from where the MA samples originated (62) (PRJNA812947) were mapped to the 38 MAG contigs.

Finally, we quantitatively compared the distribution of sulfur cycling primary MAG sequences across sites and vegetation (in AL and MA), sample depths (AL only), and sampling month (MA only) using permutational multivariate analysis of variance (PerMANOVA) (63) and post-hoc comparisons (64), based on the average CPM values calculated for each MAG. Pairwise statistical analysis of metagenomic profiles (STAMP) was used to identify primary MAGs for which read-mapping was statistically more abundant in particular sites and/or vegetation zones (65). Non-metric multidimensional scaling (NMDS) analyses were generated using the function metaMDS (63).

### Analysis of genomic diversity

We selected eight primary MAGs for further analyses, two from each site and vegetation, that were detected (by Quant_bin) at all sampling locations, and that varied notably in average CPM across samples. Six were sulfur-oxidizers (ALJR36, ALSA124, MASA10, MASA4, MASP15, MASP16) and two were sulfate-reducers (ALJR15, and ALSA174).

Genomic diversity was analyzed using Anvi’o v7.1 “hope” following the workflow outlined in (66). Open reading frames (ORFs) for each MAG were identified with Prodigal v2.6.3 (67) and HMMER (68) and annotated with COG20 (69) and GhostKoala (70). Anvi-init-bam was used to sort bam files generated with Bowtie2 (71) and calculate coverage and genetic variability metrics. Site-specific variability was analyzed in a subset of representative genes (Supplemental Data 7) including ribosomal proteins, DNA-directed RNA polymerase subunits, translation initiation factor, representative genes of the broad metabolism (72), and sulfur cycling genes. For each gene, we calculated the average gene entropy using custom R scripts (Supplemental Material).

### Pangenomic analysis of reference-guided, sample-specific, assembled MAGs

A reference-guided reassembly strategy allowed us to reconstruct sample-specific MAGs and capture the genomic variability of a given taxon across all the study sites. We generated new, sample-specific MAGs guided by the selected eight primary MAGs using BWA (73) and SPAdes (74) using the reassemble_bins module from metaWRAP (49). Primary MAG and the derived reassembled genomes were analyzed with the Anvi’o pangenomics pipeline [https://merenlab.org/2016/11/08/pangenomics-v2/]. Gene functions of the derived MAGs were assigned using NCBI COG20 (69), HMM hits from Kofam (75), and EBI’s Pfam database (76). The predicted amino acid sequence for a set of core genes was extracted with anvi-get-sequences-for-gene-clusters (Supplemental Data 9) and used to build maximum-likelihood phylogenetic trees with FastTree (77). The module anvi-compute-functional-enrichment was used to identify functions and pathways differentially represented in pan-groups consisting of site-specific MAGs at the various sites and vegetation zones.

### Identification of diversity-generating retroelements in sulfur cycling bacteria

Finally, to investigate possible sources of genetic dynamism, we explored the distribution and activity of diversity-generating retroelements (DGRs) (78). DGRs were identified in binned assemblies with the python package DGRpy (https://pypi.org/project/DGR-package/), which annotates the essential DGR features (retrotranscriptase RT, variable region VR, and template region TR) within 20kbp windows (78, 79). Putative DGR hypermutation target genes were identified by manual curation of the contigs. The dynamism of the DGRs was detected by visually inspecting the variability of recruited reads against the VR.

## Acknowledgements

This study was supported by a Gordon and Betty Moore Foundation grant #9437 to ZGC, ELP, BP, SER, and AEG, and by grants from the Simons Foundation (824763, to SER) and the National Science Foundation (2224608, to AEG). We thank Richard Fox and the MBL Bay Paul Center for supporting necessary computing resources.

## Data availability

All sequence data is available at NCBI under BioProject IDs PRJNA620304, PRJNA539072, and PRJNA814317. Accession numbers for individual metagenomes can be found in Table S1.

## Author contributions

S.P.C. and E.LP. designed the study, processed and analyzed the metagenomic data, and with Z.G.C. wrote the manuscript with input from all co-authors. O.M. and B.M. collected and sequenced Alabama samples. J.V. and J.B. collected and sequenced Massachusetts samples, A.G. and B.P. processed and analyzed the DGR surveys. S.P.C., E.LP., S.E.R., B.P., A.E.G., and Z.G.C. conceived and designed the study.

