## Supplemental Material for "Diversity at single nucleotide to pangenome scales among sulfur cycling bacteria in salt marshes"

### Diversity from single nucleotide variant to metagenome scales among sulfur cycling bacteria in Massachusetts and Alabama salt marsh

Sherlynette Pérez Castro, Elena L. Peredo, Olivia Mason, Joseph Vineis, Jennifer Bowen, Behzad Mortazavi, Anakha Ganesh, S. Emil Ruff, Blair Paul, Anne E. Giblin, Zoe G. Cardon

This pdf includes:

1. Supplemental Methods
2. Additional Results
3. References
4. Table S1-S8
5. Supplemental Data files description S1-S9
6. Figures S1-S17

#### 1. Supplemental Methods

**Note.** All scripts used for analysis can be found at https://github.com/elperedo/SaltMarshMBL

***Data set collection.*** All sequence data were downloaded from the National Center for Biotechnology Information (NCBI) Sequence Read Archive (SRA) using the SRA toolkit version 2.10. Metagenome samples were downloaded as .sra files using prefetch, checked with vdb-validate, and extracted to a readable format (.fastq.gz) using fastq-dump. Initial evaluation of read quality (raw reads) was performed using fastQC (1). Low-quality reads, unpaired reads, and sequencing adapters were removed using Trimmomatic (version 0.36) (2). Duplicated, low-quality reads (< 20 average PHRED score and/or Ns), and short reads (<60 bp) were removed using Preprocessing and Information of SEQuence data (PRINSEQ, version 0.20.4) (3).

***Metagenomic assembly, binning, and analyses.*** We divided the 24 metagenomic samples in the dataset into four groups, each of them encompassing samples from the same geographical site and vegetation type, that were used to generate co-assemblies. These four co-assemblies were ALJR (8 samples), ALSA (7), MASA (5), and MASP (5) (Table S1). Reads assembled using MEGAHIT (version 1.2.9) (4) using Kmer sizes 79, 99, 119, and 141 and selecting a minimum contig length of 3000 nt. Taxonomic classification of the contigs assembled for each of the co-assemblies was performed using Kraken2 (5). The number of contigs, length, N50, and other indicators of quality were evaluated for each assembly using MetaQUAST (6). Contigs longer than 3000 bp were assigned into bins using MetaWRAP processing modules with initial extraction using MaxBin2 (7), metaBAT2 (8), and CONCOT (9). MAGs were finalized using the bin refinement module (10). CheckM was used to assess the completeness and contamination of each resulting MAG (11). MAGs quality was reported based on (12) and only those with >90 % genome completeness and less than 5 % contamination were retained for further analysis.

MAGs were annotated using Distilled and Refined Annotation of Metabolism (DRAM), with default parameters (13). DRAM searches amino acid sequences against multiple databases such as KEGG, MEROPS, and dbCAN/CAZYmes, and consolidates annotations in a single database (DRAM.py annotate) and graphical summary (DRAM.py distill). For each MAG, pathways were manually curated by searching key sulfur genes in the raw (Supplemental Data 1) and metabolism summary (Supplemental Data 2) output files provided by DRAM. Taxonomy and closest phylogenetic neighbors of sulfur-cycling bacteria were assigned using GTDB-tk (v1.5.0) (14). Average nucleotide identity (ANI) was calculated using EZBioCloud OrthoANIu Calculator (15). Phylogenetic placement of MAGs was performed by generating a maximum likelihood phylogenetic tree of genomes using the Bacterial and Viral Bioinformatic Resource Center (16, 17).

***Distribution of primary MAGs genomic content across samples.*** We used the metaWRAP Quant_bins module (10) to identify the occurrence of each MAG across all samples included in this study. Quant_bins uses Salmon (18) to align reads from each sample to the contigs generated in the assembly, producing coverage values for each contig. To accurately quantify mapping efficiency, each subset of MAGs (ALSA, ALJR, MASP, MASA) was run against all samples included in the analysis, using the original respective co-assembly. These coverage values are standardized by library size (for every 1,000,000 metagenomic reads) and by contig length, similar to transcripts per million (TPM) in RNAseq analysis, and expressed as counts per million (CPM). Then, Quant_bins estimates the global abundance of each MAG in each sample by computing a length-weighted average of the MAG's contigs CPM values. Average CPM values for each MAG and sample were visualized using a bubble plot generated in R with a custom script. Samples with global abundance below 2 CPM were filtered out. For each MAG, we also analyzed the distribution of the mapped reads across the different contigs composing each MAG by plotting the read recruitment results, expressed as CPM (Fig. S3-S5). The individual contig coverage was obtained using a custom-made script to extract the contig-specific coverage values calculated by Salmon in Quant_bin for each sample and MAG (Supplemental Data 3).

We compared the microbial communities represented by the MAGs by vegetation type and site. We conducted a permutational multivariate analysis of variance in R (PerMANOVA, adonis function of the vegan package) (19) and post-hoc comparisons (pairwise.perm.manova function within the RVAideMemoire package) (20), based on the average CPM values (standardized estimated abundance) calculated for each MAG. Statistical analysis of metagenomic profiles (STAMP) based on CPM values was used to identify lineages that were statistically more abundant among locations and vegetation types (11). Non-metric multidimensional scaling (NMDS) analysis was used to visualize microbial communities represented by the MAGs by location and type of vegetation using the function metaMDS in the vegan package (19). Component scores were used to generate 95 % confidence ellipses around samples using the ordiellipse() function within Vegan (19).

We used the metaWRAP Quant_bins module (10) as indicated above to identify the occurrence of the MAGs identified in this paper in environmental metagenomic samples published and available from NCBI for bare salt marsh sediments in Plum Island (MA, USA) (n= 32, SRR24130617-48). For sample description consult Table S8. We selected the Plum Island dataset because it originated in the same salt marsh area as the Massachusetts dataset so the environmental conditions of the bare salt marsh sediments were broadly similar to those in the vegetated ones. Therefore, we expected that the bacterial communities in this site would share some similarities with those of vegetated areas. The sequencing depth of this dataset was very low, which hindered previous attempts at MAG recovery. As previously, MAG occurrence was calculated as the average CPM for each MAG and sample. Results were visualized in a bubble plot. The distribution of the mapped reads across the different contigs composing each MAG was investigated by plotting the contig-specific read recruitment results (Supplemental Data 4).

***Analysis of genomic diversity.*** We selected eight primary MAGs to further analyze the genetic and genomic diversity of sulfur-cycling bacterial populations inhabiting sediments characterized by different salt marsh vegetation and in contrasting geographical localities. We included in the analysis two MAGs per metagenomic co-assembly. Six of the MAGs corresponded to SOX bacteria (ALJR36, ALSA124, MASA10, MASA4, MASP15, MASP16) and two as SRB (ALJR15, and ALSA174). Additional selection criteria in this analysis included 1) varying levels of occurrence across samples determined by Quant_bin (10) and 2) taxonomic assignment, as when possible, we favored in this analysis those MAGs representing understudied taxa.

We used Anvi'o v7.1 "hope" (available from https://github.com/merenlab/anvio/releases/v7) to profile mapping results, finalize genomic bins, annotate, and visualize results following the workflow outlined in (21). In brief, we used anvi-gen-contigs-database to identify open reading frames (ORFs) with Prodigal v2.6.3 (22) and HMMER (23) to identify single-single copy core genes of bacterial and archaeal origin. We used anvi-run-ncbi-cogs to annotate the identified ORFs using the NCBI Clusters of Orthologous Genes (COG) database (COG20 release) (24) and KEGG Orthology using GhostKoala (25–27). Gene taxonomy was determined using the top hits against Centrifuge (28) and from the Kegg Taxonomy results produced by GhostKoala. The MAG taxonomical identity was resolved using anvi-run-scg-taxonomy. Anvi-init-bam was used to incorporate into Anvi'o the mapping results generated with Bowtie2 (29) using end-to-end alignment and sensitive presets ( -D 15 -R 2 -N 0 -L 22 -i S,1,1.15) to minimize spurious read recruitment. Anvi-profile with the flag –profile-SCVs was used to process each BAM file and to calculate coverage and genetic variability metrics. Anvi-merge was used to combine profiles from each metagenomic sample and create a merged Anvi'o profile of each MAG in the dataset. Finally, the results of the analysis were visualized with anvi-interactive.

For each profiled primary MAG, we selected a subset of representative genes to analyze site-specific variability using average entropy as a measurement of diversity. The gene subset included the genes encoding the ribosomal proteins, DNA-directed RNA polymerase subunits, and translation initiation factors used by CheckM (11) to calculate metagenome completeness. We also analyzed average entropy for genes representative of the broad metabolism (30), and genes involved in sulfur-cycling processes (see Supplemental Data 9 for the full list). For each of the genes, we identified single codon variants (SCVs) using the program anvi-gen-variability-profile as implemented in Anvi'o (21) run with the flag –kiefl-mode. In all cases, we computed gene-specific coverage statistics. We used custom-made R scripts to filter, summarize and visualize the data generated by anvi-gen-variability-profile. To summarize the variability level of each gene, we created the function average gene entropy. In it, and for each gene, we calculated the gene average entropy normalized to the gene length. [(entropy.sum/entropy.n)/gene_length]

***Reconstruction of reference-guided, sample-specific, assembled MAGs.*** We generated sample-specific reassembled MAGs using the reassemble_bins module implemented in metaWRAP (10). The MAGs selected for further analysis were the same as those previously selected for Anvi'o profiling. In this portion of the analysis, the MAGs generated through the coassembly strategy were used as a general reference to guide sample-specific assemblies using each of the 24 metagenomic samples listed in Table S1. In brief, the reassemble_bins module metaWRAP uses BWA (31) to align the reads in a metagenomic sample to a reference MAG followed by a genome reconstruction using SPAdes (32). This pipeline uses a parallel approach to reassembly derived from permissive and strict mapping conditions in the BWA step (2 and 5 mismatches respectively). Here, we only considered the results of the reassembly under strict conditions. The number of reads mapped per metagenomic sample to each MAG is presented in Supplemental Table 4. Initial completeness of the reference-guided, sample-specific, assembled MAGs was estimated using CheckM (11).

***Pangenomic analysis of the reference-guided, sample-specific, assembled MAGs***
We used the Anvi'o pangenomics pipeline [https://merenlab.org/2016/11/08/pangenomics-v2/] for the analysis of the suite of reference-guide genomes reassembled from each of the eight MAGs selected for further analysis. For each bacterial taxon, we stored the primary MAG generated through a coassembly (see Assembly and taxonomic assignment of contigs section) and each of the reference-guided reassembled metagenome in an Anvi'o database using the command anvi-gen-genomes-storage. We identified single-copy genes and rRNAs using anvil-run-hmms. Gene functions were assigned using NCBI COG20 (24), HMM hits from Kofam, and EBI's Pfam database (33). Additionally, tRNAs were scanned using anvi-scan-trnas. Anvi-run-scg-taxonomy was used to confirm taxonomy and anvi-estimate-metabolism to estimate metabolic capabilities.

To create the pangenomes from each metagenome, we used the command anvi-pan-genome which used NCBI's BLAST (34) to quantify gene similarity within and between genomes in each database and Markov Cluster algorithm (MCL) (35) to cluster groups of similar genes. We set the flag --min-ocurrence to 2 to remove singletons. The resulting pangenomes were visualized using anvi-display-pan. In the case of the closely related MAGs ALJR36 and MASA10, we completed an additional pangenomic analysis comparing the groups of metagenomes reassembled across samples for each of these sulfur-oxidizer bacteria affiliated with the UBA6429 family. We completed a functional enrichment analysis using the module anvi-compute-functional-enrichment to identify those genes, functions, and pathways over-represented in the pan-groups (reference MAG, and site and vegetation).

***Analysis of the phylogenetic relationships among the reference-guided reconstructed MAGs.*** We used the command anvi-get-sequences-for-gene-clusters to extract, from each metagenome, the predicted amino acid sequence of a subset of core genes (ribosomic proteins, polymerases, and transcription factors), housekeeping genes, and sulfur-cycling genes (Supplemental Data 9). Amino acid gene sequences were concatenated and samples with a high percentage of missing data in the gene alignments (>90% of missing sites) were removed from the analysis. Maximum-likelihood phylogenetic trees were calculated using FastTree (36) as implemented in anvi-gen-phylogenomic-tree.

***Identification of Diversity-Generating Retroelements in sulfur-cycling bacteria.*** DGRs were identified with the python package DGRpy (available at GitHub: BP_INSERT_LINK), which provides an integrated pipeline to annotate the essential features, RT, VR, and TR within 20kbp windows, as previously described (37, 38). Genomic sequences were used as the input for DGRpy and open reading frame (ORF) predictions determined using Prodigal with -p meta and otherwise default parameters (22). Next, ORFs were analyzed for RT homology via HMMER (23) against a custom RT-HMM profile (38). The genomes were processed into local 20 Kbp regions, extracted with +/- 10 Kbp flanking the RT-like sequence. The RT-proximal sequences were next used to generate short sliding windows as 200 nt fragments that overlap by 50 nt. All 200 nt sequences were compared via BLASTn (34) and the resulting XML was parsed to search for VR/TR pairs in alignments having an HSP score < 100 and alignment length ≥ 60 nt. A VR/TR pair was identified in alignments having mismatches that correspond to adenines (or thymidine if on the reverse strand) for one of the two pairwise sequences. Alignments with i) a relative proportion of 80% A-mismatches (or reverse-strand T-mismatches) along one sequence, relative to other mismatches (G, C, or T) and ii) a minimum of five A-/T- specific mismatches, were considered VR/TR pairs. Since the BLAST xml output contains both full and partial TRs/VRs, sequences were clustered at 100% identity using CD-HIT (39), to obtain the full-length representatives. Finally, RTs, VRs, and TRs were manually inspected in the original MAGs to identify the putative DGR hypermutation target genes.

#### 2. Additional results

***MAGs occurrence across all vegetated samples.*** In this work, we used a bioinformatic approach to estimate the occurrence of the newly assembled MAGs based on read recruitment. We quantified recruitment as average counts per million (CPM) per bin, to estimate the presence of a given MAG across all sites and samples (Fig. 4, main text). Because the metric CPM is normalized by the number of reads present in a given library, a comparison of read recruitment across samples is possible. We completed an additional analysis of the read recruitment at the contig level for each MAG (Supplemental Data 3). The graphic summary of the results of these analyses can be found in Figures S3-S5. The analysis at the contig level allowed us to confirm that the average occurrence calculated for each bin was a fair representation of the occurrence of a given MAG in a sample and not just driven by high read recruitment in a few contigs within that bin. In general, we observed that the samples that were part of the subset of samples originally used in the co-assembly of a given bin had a relatively homogeneous distribution of reads recruitment among the contigs confirming that bin (Fig. S3-S5). The CMP values per contig were more variable for those samples collected across larger geographical distances, as could be anticipated by the expected presence of potential genomic variability across sites. In samples where the global read recruitment value was very low (<2), read recruitment was often restricted to a subset of contigs consistent with high conservation of sections of the bacterial genomes across broader phylogenetic distances.

***MAGs occurrence estimates across other metagenomes***. We used the same bioinformatic approach based on read recruitment to estimate the MAGs' occurrence in metagenomic samples collected in other environments. We selected a dataset composed of sediment samples collected across a subtidal gradient in Plum Island LTER. These sediments were devoid of vegetation but still shared some broad environmental factors (e.g. salinity, tidal system) with the marsh sediment samples collected in Massachusetts. The sequencing depth of these samples was low (ranging between 5 and 10M) and was not enough to produce complete metagenomes. We bypassed the sequencing depth limitation by using the estimation of abundance based on read recruitment to identify the presence of the MAGs newly described in this study in these subtidal sites (Fig. S6).

We also completed an additional analysis of CPM values at the individual contig level. The coverage patterns, as relatively constant values across contigs included in a bin, were similar to those expected in samples containing S-cycling bacteria that were highly similar to the MAGs used as references for mapping reads. This was especially apparent with some of the MAGs reconstructed using samples collected in sediments inhabited by *S. alterniflorus* in Massachusetts (see Supplemental Data 4; MASP12, MASA6, MASA5, MASA4, MASA3, MASA1, ALSA234, ALSA191, ALSA113, ALJR36, ALJR32, ALJR30, ALJR18, ALJR15, ALJR4). As was the case on the vegetated sediments, higher read recruitment in sulfur-oxidizing lineages when compared to sulfate-reducers were also observed in these subtidal metagenome samples. In Particular, we detected the presence of sulfur-oxidizing taxa affiliated with *Alphaproteobacteria* and *Burkholderiales* MAGs across all sites and environments (Fig. S6).

The estimation of abundance using reference-guided mapping unveiled several trends that correlated with environmental factors. The effect of salinity could explain the changes in abundance for the *Gammaproteobacteria* MASP12. This sulfur-oxidizer bacterium was detected in all subtidal metagenomes with a notable increase in abundance towards higher salinities (G1 – G6 and R1 – R6, Fig. S6). In contrast, *Alphaproteobacteria* and *Burkholderiales* MAGs were found in a decreasing trend toward salinity (Fig. S6). Sulfate-reducing *Acidobaceriota* Fen-336 family, which was spread across marsh samples, was also detected in subtidal metagenomes.

***Gene heterogeneity as average entropy.*** We calculated the average entropy for a selected set of genes (listed in Supplemental Data 9) to get a more accurate picture of the distribution of the genetic heterogeneity in the coding regions of the genomes of these bacterial populations. While the SNV density is useful to identify mismatches to a 'reference' sequence, entropy provides a quantification of the disagreement from the consensus, measuring the diversity in a bacterial population by removing the possible effect of the fixed SNV in a given bacterial population. By focusing on the entropy of single codon variants (SCV) we specifically explored the changes affecting the triplets of nucleotides encoding amino acids, independently if they were synonymous changes or not.

Only four MAGs (ALJR36, ALJR15, ALSA124, and MASA10), specifically in the metagenomic samples collected in Alabama, independently of vegetation type, had enough coverage to fulfill the minimal coverage depth set as in the analysis. Average gene entropy values varied among genes annotated for a given MAG. In most cases, average entropy values were higher for the genes encoding ribosomal proteins than for housekeeping or sulfur-related genes (Fig. S16 and Supplemental Data 7). Entropy values were not dependent on the average gene coverage as indicated for the similar values of average entropy calculated for a given gene in diverse samples (Fig. S16A, heatmap in purple) despite variations in coverage (Fig. S16A heatmap in orange).

The main result we observed is that for a given gene of a given MAG, the entropy values calculated remained relatively constant across different bacterial populations in different samples analyzed here, independently of the year of collection or the plant dominating the sediments. More interestingly, we identified that the average entropy values of a given gene were MAG-specific. To better showcase these results, we compared the average entropy values of housekeeping genes and sulfur-cycling genes in the SOXs ALJR36 and MASA10 (Fig. S16b) and specifically, using samples collected in Alabama as they were consistently abundant and therefore produced more robust results. The average entropy values, and therefore diversity in the bacterial population, calculated e.g. for the gene Succinate dehydrogenase *Sdh*A−1 (COG1053) were consistently higher for ALJR36 than for MASA10. This was also observed for the genes encoding subunits alpha (K17993) and delta (K17994) of the Sulfhydrogenase (*Hyd*) while in MASA10 the entropy values of the two subunits were similar. In ALJR36 the values were not only much higher but the average entropy value for the alpha subunit almost doubled that for the delta subunit. In the case of the dissimilatory sulfite reductase (*dsr*), the average entropy calculated for the genes encoding subunits alpha (K11180) and beta (K11181) were similar for the bacterial populations matching MASA10 (~0.001). Values of dsrA in ALJR36 were also in that range while dsrB displayed a considerably lower genetic diversity.

**Table S1.** Summary of sampling locations, dominant vegetation, depth, dates, and the number of reads of the selected metagenomic datasets. (AL, Alabama; MA, Massachusetts).

| Samples (N) | Site | Vegetation type | Collection date | Depth (cm) | JGI Gp | Raw Reads | QC Reads |
|---|---|---|---|---|---|---|---|
| 1 | AL | *J. roemerianus* | 2015-05 | 5-7 | Gp0344160 | 91,291,894 | 83,691,004 |
| 2 | AL | *J. roemerianus* | 2015-05 | 0-2 | Gp0344159 | 77,016,234 | 66,168,546 |
| 3 | AL | *J. roemerianus* | 2015-05 | 0-2 | Gp0344159 | 45,424,663 | 40,593,738 |
| 4 | AL | *J. roemerianus* | 2016-05 | 5-7 | Gp0344164 | 96,984,667 | 89,533,791 |
| 5 | AL | *J. roemerianus* | 2016-05 | 0-2 | Gp0344163 | 117,191,856 | 107,792,129 |
| 6 | AL | *J. roemerianus* | 2017-05 | 0-10 | Gp0344168 | 66,913,351 | 59,159,686 |
| 7 | AL | *J. roemerianus* | 2017-05 | 0-10 | Gp0344167 | 47,964,230 | 41,031,419 |
| 8 | AL | *J. roemerianus* | 2017-05 | 0-10 | Gp0344167 | 104,213,562 | 90,031,614 |
| **Total** | | | | | | | **578,001,927** |
| 1 | AL | *S. alterniflorus* | 2015-05 | 5-7 | Gp0344162 | 102,973,665 | 94,441,225 |
| 2 | AL | *S. alterniflorus* | 2015-05 | 0-2 | Gp0344161 | 157,379,608 | 142,491,993 |
| 3 | AL | *S. alterniflorus* | 2016-05 | 5-7 | Gp0344166 | 54,423,170 | 46,496,251 |
| 4 | AL | *S. alterniflorus* | 2016-05 | 0-2 | Gp0344165 | 63,219,940 | 55,527,119 |
| 5 | AL | *S. alterniflorus* | 2017-05 | 0-10 | Gp0344170 | 66,689,192 | 58,817,185 |
| 6 | AL | *S. alterniflorus* | 2017-05 | 0-10 | Gp0344169 | 81,630,750 | 72,191,424 |
| 7 | AL | *S. alterniflorus* | 2017-05 | 0-10 | Gp0344169 | 49,833,666 | 44,316,881 |
| **Total** | | | | | | | **514,282,078** |
| 1 | MA | *S. alterniflorus* | 2015-05 | 0-5 | Gp0432379 | 48,929,864 | 43,938,191 |
| 2 | MA | *S. alterniflorus* | 2015-05 | 0-5 | Gp0432385 | 47,493,408 | 42,331,083 |
| 3 | MA | *S. alterniflorus* | 2015-08 | 0-5 | Gp0432391 | 52,916,253 | 46,675,383 |
| 4 | MA | *S. alterniflorus* | 2015-08 | 0-5 | Gp0432397 | 57,416,483 | 50,747,510 |
| 5 | MA | *S. alterniflorus* | 2015-10 | 0-5 | Gp0432373 | 53,164,672 | 47,120,515 |
| **Total** | | | | | | | **230,812,682** |
| 1 | MA | *S. pumilus* | 2015-05 | 0-5 | Gp0432378 | 44,550,785 | 40,039,222 |
| 2 | MA | *S. pumilus* | 2015-08 | 0-5 | Gp0432390 | 41,065,476 | 36,950,534 |
| 3 | MA | *S. pumilus* | 2015-08 | 0-5 | Gp0432396 | 44,913,560 | 40,622,394 |
| 4 | MA | *S. pumilus* | 2015-10 | 0-5 | Gp0432366 | 67,768,852 | 58,336,509 |
| 5 | MA | *S. pumilus* | 2015-10 | 0-5 | Gp0432372 | 35,457,645 | 31,923,829 |
| **Total** | | | | | | | **207,872,488** |
| **Sum** | | | | | | | **1,530,969,175** |

**Table S2.** Summary statistics for each of the metagenomic co-assemblies.

| Assembly | ALJR | ALSA | MASA | MASP |
|---|---|---|---|---|
| # contigs (>= 3000 bp) | 251,034 | 249,343 | 70,380 | 100,526 |
| # contigs (>= 5000 bp) | 103,473 | 106,678 | 27,050 | 41,948 |
| # contigs (>= 10000 bp) | 30,574 | 34,103 | 7,566 | 13,088 |
| # contigs (>= 25000 bp) | 4,984 | 6,331 | 1,301 | 2,412 |
| # contigs (>= 50000 bp) | 937 | 1,286 | 281 | 487 |
| Total length (>= 3000 bp) | 1,621,879,533 | 1,700,038,878 | 440,187,433 | 672,932,647 |
| Total length (>= 5000 bp) | 1,065,541,187 | 1,161,729,582 | 277,515,921 | 452,063,659 |
| Total length (>= 10000 bp) | 573,987,950 | 669,974,094 | 146,797,394 | 256,972,975 |
| Total length (>= 25000 bp) | 202,235,813 | 263,003,851 | 56,123,242 | 101,158,766 |
| Total length (>= 50000 bp) | 67,990,329 | 95,150,165 | 22,126,090 | 36,675,331 |
| # contigs | 251,034 | 249,343 | 70,380 | 100,526 |
| Largest contig | 294,481 | 311,488 | 311,637 | 276,711 |
| Total length | 1,621,879,533 | 1,700,038,878 | 440,187,433 | 672,932,647 |
| GC (%) | 59 | 59 | 58 | 61 |
| N50 | 6,860 | 7,514 | 6,470 | 7,248 |
| N75 | 4,265 | 4,417 | 4,117 | 43,432 |
| L50 | 59,575 | 55,022 | 16,873 | 22,506 |
| L75 | 136,219 | 130,825 | 38,696 | 53,266 |
| # N's per 100 kbp | 0 | 0 | 0 | 0 |

**Table S3.** Genome size, GC content, and taxonomic placement of the MAGs identified in this study.

| MAG | Size (Mb) | GC (%) | GTDB Taxonomy |
|---|---|---|---|
| ALJR11 | 3.3 | 65.5 | Proteobacteria; Gammaproteobacteria; Chromatiales; Chromatiaceae; Thiocapsa; |
| ALJR12 | 4.1 | 49.5 | Desulfobacterota; Syntrophobacteria; BM002; BM002; BM002; |
| ALJR14 | 4.4 | 69.4 | Gemmatimonadota; Gemmatimonadetes; Longimicrobiales; UBA6960; SZUA-318; |
| ALJR15 | 3.6 | 47.3 | Desulfobacterota; Desulfobacteria; Desulfobacterales; UBA11574; UBA11574; |
| ALJR17 | 2.2 | 54.4 | Proteobacteria; Gammaproteobacteria; Acidiferrobacterales; |
| ALJR18 | 3.5 | 64.2 | Proteobacteria; Gammaproteobacteria; Chromatiales; Sedimenticolaceae; 41T-STBD-0c-01a; |
| ALJR2 | 3.8 | 50.0 | Proteobacteria; Gammaproteobacteria; Thiohalomonadales_A; SZUA-152 |
| ALJR24 | 3.8 | 66.6 | Acidobacteriota; Thermoanaerobaculia; Thermoanaerobaculales; FEB-10; JAADFK01; |
| ALJR25 | 4.6 | 67.7 | Gemmatimonadota; Gemmatimonadetes; Longimicrobiales; UBA6960; SZUA-318; |
| ALJR28 | 3.6 | 67.5 | Proteobacteria; Gammaproteobacteria; Burkholderiales; SG8-39; SG8-39; |
| ALJR30 | 3.8 | 35.2 | Bacteroidota; Ignavibacteria; Ignavibacteriales; Ignavibacteriaceae; BMS3ABIN03; |
| ALJR32 | 3.9 | 50.4 | Desulfobacterota; Syntrophobacteria; BM002; BM002; BM002; |
| ALJR35 | 3.9 | 61.7 | Acidobacteriota; Thermoanaerobaculia; Thermoanaerobaculales; FEB-10 |
| ALJR36 | 2.2 | 54.8 | Proteobacteria; Gammaproteobacteria; Thiohalomonadales; UBA6429 |
| ALJR4 | 2.3 | 57.4 | Proteobacteria; Gammaproteobacteria; Burkholderiales; Gallionellaceae; Gallionella; |
| ALSA113 | 3.4 | 47.1 | Proteobacteria; Gammaproteobacteria; SZUA-229; SZUA-229; GCA-2746365; |
| ALSA124 | 4.8 | 68.2 | Proteobacteria; Gammaproteobacteria; Chromatiales; Chromatiaceae; Thiohalocapsa; |
| ALSA142 | 3.3 | 57.3 | Desulfobacterota; Desulfobulbia; Desulfobulbales; Desulfocapsaceae; JABDQA01; |
| ALSA143 | 6.3 | 70.2 | Acidobacteriota; Vicinamibacteria; Fen-336; Fen-336 |
| ALSA159 | 3.8 | 49.1 | Desulfobacterota; Syntrophobacteria; BM002; BM002; BM002; |
| ALSA174 | 4.2 | 35.2 | Bacteroidota; Ignavibacteria; Ignavibacteriales; Ignavibacteriaceae; BMS3ABIN03; |
| ALSA176 | 5.0 | 69.5 | Acidobacteriota; Vicinamibacteria; Fen-336; Fen-336 |
| ALSA183 | 4.4 | 55.2 | Desulfobacterota; Desulfobacteria; Desulfobacterales; B30-G6 |
| ALSA191 | 2.8 | 56.0 | Proteobacteria; Gammaproteobacteria; Thiohalobacterales; UBA9214; UBA9214; |
| ALSA234 | 4.0 | 63.3 | Desulfobacterota; Desulfobacteria; Desulfobacterales; JAABRJ01; JAABRJ01; |
| ALSA27 | 3.2 | 65.6 | Proteobacteria; Gammaproteobacteria; Chromatiales; Chromatiaceae; Thiocapsa; |
| ALSA42 | 4.0 | 59.8 | Desulfobacterota; Desulfobacteria; Desulfobacterales; UBA2174 |
| MASA1 | 3.9 | 63.5 | Proteobacteria; Alphaproteobacteria; Rhodobacterales; Rhodobacteraceae; Rhodobacter_B; |
| MASA10 | 2.3 | 55.0 | Proteobacteria; Gammaproteobacteria; Thiohalomonadales; UBA6429 |
| MASA3 | 3.3 | 63.3 | Proteobacteria; Alphaproteobacteria; Rhodobacterales; Rhodobacteraceae; Silicimonas; |
| MASA4 | 2.8 | 64.3 | Proteobacteria; Gammaproteobacteria; Arenicellales; HyVt-429 |
| MASA5 | 4.3 | 56.1 | Proteobacteria; Gammaproteobacteria; GCA-001735895; GCA-001735895; GCA-001735895; |
| MASA6 | 4.4 | 65.9 | Proteobacteria; Gammaproteobacteria; Burkholderiales; SG8-39; JAABRB01; |
| MASP1 | 5.0 | 50.6 | Desulfobacterota; Desulfobacteria; Desulfatiglandales; Desulfatiglandaceae; NaphS2; |
| MASP12 | 3.4 | 61.4 | Proteobacteria; Gammaproteobacteria; Thiohalobacterales; DSM-19610; Thiogranum; |
| MASP15 | 3.9 | 61.5 | Proteobacteria; Gammaproteobacteria; UBA4575; JACCYU01 |
| MASP16 | 3.4 | 61.8 | Proteobacteria; Alphaproteobacteria; Rhizobiales; Rhizobiaceae; Salaquimonas; |
| MASP6 | 4.4 | 61.2 | Proteobacteria; Gammaproteobacteria; Chromatiales; Chromatiaceae |

**Table S4.** Number of reads recruited per MAG after Bowtie2 metagenomic sample alignment.

| Site | Sample | QC Reads | ALSA 124 | ALSA 174 | ALJR 36 | ALJR 15 | MASA 10 | MAS A4 | MASP 15 | MASP 16 |
|---|---|---|---|---|---|---|---|---|---|---|
| ALJR | SRR11061153 | **66,168,546** | 91,688 | 25,241 | 56,844 | 205,401 | 55,677 | 2,340 | 14,527 | 6,510 |
| ALJR | SRR11061154 | **40,593,738** | 53,120 | 17,621 | 37,973 | 137,232 | 37,214 | 1,549 | 9,459 | 4,516 |
| ALJR | SRR11567260 | **90,031,614** | 195,351 | 107,487 | 173,642 | 104,455 | 169,858 | 1,959 | 12,945 | 8,576 |
| ALJR | SRR11567261 | **41,031,419** | 88,205 | 49,593 | 80,280 | 47,615 | 79,093 | 906 | 5,938 | 4,136 |
| ALJR | SRR9045291 | **83,691,004** | 85,564 | 69,877 | 106,631 | 384,306 | 104,804 | 2,516 | 13,924 | 20,028 |
| ALJR | SRR9045294 | **107,792,129** | 560,620 | 103,103 | 134,973 | 151,345 | 132,545 | 2,644 | 27,358 | 6,436 |
| ALJR | SRR9045295 | **59,159,686** | 194,273 | 78,062 | 123,209 | 320,041 | 120,770 | 2,031 | 12,806 | 9,257 |
| ALJR | SRR9045297 | **89,533,791** | 235,235 | 125,168 | 127,352 | 95,328 | 124,740 | 2,099 | 16,693 | 10,299 |
| | *Average* | *72,250,240* | *188,007.0* | *72,019.0* | *105,113.0* | *180,715.4* | *103,087.6* | *2,005.5* | *142,06.3* | *8,719.8* |
| ALSA | SRR10854653 | **46,496,251** | 77,199 | 117,876 | 70,538 | 28,116 | 69,321 | 937 | 3,702 | 3,071 |
| ALSA | SRR11567157 | **116,508,305** | 628,802 | 101,396 | 299,068 | 58,002 | 294,029 | 2,854 | 17,103 | 8,821 |
| ALSA | SRR9045292 | **142,491,993** | 948,105 | 51,523 | 69,618 | 92,285 | 68,649 | 4,537 | 22,458 | 8,663 |
| ALSA | SRR9045293 | **94,441,225** | 415,145 | 96,176 | 174,055 | 43,758 | 171,089 | 2,054 | 9,960 | 4,508 |
| ALSA | SRR9045296 | **58,817,185** | 212,625 | 56,378 | 209,538 | 44,498 | 205,944 | 1,368 | 8,981 | 3,630 |
| ALSA | SRR9045298 | **55,527,119** | 276,908 | 46,031 | 107,904 | 76,680 | 106,493 | 1,665 | 8,923 | 2,891 |
| | *Average* | *85,713,679* | *426,464.0* | *78,230.0* | *155,120.2* | *57,223.2* | *152,587.5* | *2,235.8* | *11,854.5* | *5,264.0* |
| MASA | SRR11828800 | **47,120,515** | 4,184 | 2,614 | 47,096 | 3,769 | 50,612 | 42,444 | 4,487 | 5,837 |
| MASA | SRR11828899 | **43,938,191** | 15,191 | 1,767 | 21,895 | 4,840 | 23,771 | 16,869 | 7,222 | 3,897 |
| MASA | SRR11829000 | **42,331,083** | 4,036 | 2,276 | 35,056 | 4,237 | 38,172 | 24,673 | 3,785 | 3,821 |
| MASA | SRR11829104 | **46,675,383** | 3,841 | 1,896 | 30,439 | 10,508 | 33,171 | 94,327 | 4,918 | 3,713 |
| MASA | SRR11829269 | **50,747,510** | 2,166 | 1,364 | 26,735 | 2,512 | 29,769 | 51,265 | 3,962 | 4,869 |
| | *Average* | *46,162,536* | *5,883.6* | *1,983.4* | *32,244.2* | *5,173.2* | *35,099.0* | *45,915.6* | *4,874.8* | *4,427.4* |
| MASP | SRR11828593 | **31,923,829** | 819 | 641 | 2,503 | 6,426 | 2,653 | 4,752 | 26,749 | 10,550 |
| MASP | SRR11828889 | **40,039,222** | 1,813 | 221 | 4,811 | 5,610 | 5,043 | 2,788 | 39,275 | 22,421 |
| MASP | SRR11829102 | **36,950,534** | 1,794 | 443 | 12,424 | 15,484 | 12,910 | 2,481 | 30,723 | 20,022 |
| MASP | SRR11829268 | **40,622,394** | 2,283 | 356 | 1,507 | 268,190 | 1,625 | 3,028 | 9,608 | 6,983 |
| MASP | SRR12659820 | **58,336,509** | 9,438 | 3,301 | 31,743 | 26,305 | 33,274 | 8,328 | 363,112 | 127,955 |
| | *Average* | *41,574,497* | *3,229.4* | *992.4* | *10,597.6* | *64,403.0* | *11,101.0* | *4,275.4* | *93,893.4* | *37,586.2* |

**Table S5**. Estimation of completeness (CheckM) of the primary MAGs and the reference-guided reassembled MAGs.[1]

| Site | Sample | ALSA 124 | ALSA 174 | ALJR 36 | ALJR 15 | MASA 10 | MASA 4 | MASP 15 | MASP 16 |
|---|---|---|---|---|---|---|---|---|---|
| | **Primary MAG** | **95.59** | **92.18** | **90.37** | **92.74** | **91.77** | **96.21** | **96.41** | **94.58** |
| ALJR | SRR11061153 | 87.28 | 79.61 | 91.59 | 93.55 | 88.82 | 64.21 | 57.41 | 57.98 |
| ALJR | SRR11061154 | 82.39 | 48.32 | 86.98 | 92.88 | 84.65 | 58.82 | 55.1 | 55.54 |
| ALJR | SRR11567260 | 86.44 | 93.3 | 93.53 | 90.3 | 92.79 | 64.05 | 66.16 | 78.37 |
| ALJR | SRR11567261 | 79.08 | 88.69 | 91.79 | 71.91 | 90.46 | 54.13 | 46.02 | 66.03 |
| ALJR | SRR9045291 | 90.46 | 15.01 | 85.29 | - | 92.16 | 73.74 | 68.67 | 83.54 |
| ALJR | SRR9045294 | 96.53 | 93.58 | 87.88 | 90.61 | 91.08 | 73.69 | 77.24 | 82.29 |
| ALJR | SRR9045295 | 93.31 | 94.97 | 92.57 | 57.92 | 91.3 | 67.14 | 76.72 | 81.71 |
| ALJR | SRR9045297 | 90.2 | 89.66 | 93.01 | - | 92.75 | 81.27 | 65.05 | 57.98 |
| ALSA | SRR10854653 | 89.83 | 92.72 | - | 48.89 | 90.57 | 59.3 | 50.44 | 57.87 |
| ALSA | SRR11567157 | 93.21 | 94.69 | 88.04 | 81.23 | 93.47 | 78.63 | 75.86 | 57.98 |
| ALSA | SRR9045292 | 98.12 | 91.76 | 87.88 | 89.61 | 88.36 | 77.25 | 91.38 | 79.7 |
| ALSA | SRR9045293 | 94.25 | 91.2 | 92.74 | 0 | 92.82 | 71.26 | 66.67 | 57.98 |
| ALSA | SRR9045296 | 90.75 | 91.9 | 93.28 | - | 92.71 | 54.37 | 61.57 | 60.55 |
| ALSA | SRR9045298 | 88.58 | 94.67 | 92.32 | - | 91.28 | 70.36 | 73.98 | 57.98 |
| MASA | SRR11828800 | 72.68 | 4.23 | 86.44 | 0 | 90.53 | 90.84 | 42.44 | 65.44 |
| MASA | SRR11828899 | 66.38 | 0 | 74.25 | 5.49 | 83.82 | 82.29 | 44.11 | 65.27 |
| MASA | SRR11829000 | 41.61 | 0 | 83.18 | 5.26 | 88.2 | 85.91 | 27.87 | 51.75 |
| MASA | SRR11829104 | 58.09 | 2.08 | 81.7 | 10.69 | 87.97 | 95.29 | 38.71 | 61.99 |
| MASA | SRR11829269 | 60.78 | 2.82 | 80.8 | 10.85 | 86.82 | 86.28 | 41.43 | 70.95 |
| MASP | SRR11828593 | 30.17 | - | - | 3.51 | 15.67 | 31.33 | 86.23 | 67.26 |
| MASP | SRR11828889 | 24.66 | - | 19.12 | 0 | 27.87 | 26.78 | 92.39 | 83.92 |
| MASP | SRR11829102 | 35.59 | - | 55.95 | 0 | 75.24 | 32.47 | 84.92 | 84.19 |
| MASP | SRR11829268 | 44.73 | 3.61 | 8.03 | 70.74 | 11.19 | 33.58 | 48.5 | 48.6 |
| MASP | SRR12659820 | 82.39 | 2.59 | 81.81 | 6.4 | 89.36 | 69.73 | 96.93 | 89.52 |

[1] Empty cells (-) indicate reference-guided reassembly efforts that failed to produce any contigs/bins.

**Table S6**. Results of the functional enrichment analysis comparing the metabolic functionalities of ALJR36 and MASA10 pangenomes at COG20 Category level. Each pangenome group is represented by the primary MAG and the site-specific reassembled MAGs.

| COG20 CATEGORY | Adjusted q value | COG ID |
|---|---|---|
| **Categories significantly enriched in the ALJR36 pangenome group.** | | |
| Carbohydrate transport and metabolism-Signal transduction mechanisms | 3.69E-08 | G-T |
| Cell motility-Carbohydrate transport and metabolism | 3.69E-08 | N-G |
| Coenzyme transport and metabolism-Transcription | 3.69E-08 | H-K |
| Energy production and conversion-General function prediction only | 1.45E-07 | C-R |
| Intracellular trafficking, secretion, and vesicular transport-Intracellular trafficking, secretion, and vesicular transport | 1.45E-07 | U-U |
| Amino acid transport and metabolism-General function prediction only-Replication, recombination and repair | 1.45E-07 | E-R-L |
| Amino acid transport and metabolism-Posttranslational modification, protein turnover, chaperones | 3.21E-06 | E-O |
| Coenzyme transport and metabolism-Inorganic ion transport and metabolism | 1.13E-05 | H-P |
| Transcription-General function prediction only | 1.13E-05 | K-R |
| Extracellular structures | 3.89E-05 | W |
| Energy production and conversion-Amino acid transport and metabolism | 0 | C-E |
| **Categories significantly enriched in the MASA10 pangenome group.** | | |
| Cell wall/membrane/envelope biogenesis-Intracellular trafficking, secretion, and vesicular transport | 3.69E-08 | M-U |
| Signal transduction mechanisms-Intracellular trafficking, secretion, and vesicular transport-Extracellular structures | 3.69E-08 | T-U-W |
| General function prediction only-Inorganic ion transport and metabolism | 1.45E-07 | R-P |
| Cell motility-Intracellular trafficking, secretion, and vesicular transport | 7.50E-07 | N-U |
| Defense mechanisms-Lipid transport and metabolism | 3.21E-06 | V-I |
| Cell wall/membrane/envelope biogenesis-Coenzyme transport and metabolism-General function prediction only | 1.13E-05 | M-H-R |
| Coenzyme transport and metabolism-Nucleotide transport and metabolism | 1.62E-05 | H-F |
| Inorganic ion transport and metabolism-Coenzyme transport and metabolism | 0.05 | P-H |

**Table S7**. Results of the functional enrichment analysis comparing the metabolic functionalities of ALJR36 and MASA10 pangenomes at the KEGG Module level. Each pangenome group includes the primary MAG and the site-specific reassembled MAGs.

| KEGG Module | Adjusted q value | KEGG ID |
|---|---|---|
| **Modules significantly enriched in the ALJR36 pangenome group.** | | |
| Glycolysis (Embden-Meyerhof pathway), glucose => pyruvate-Glycolysis, core module involving three-carbon compounds-Gluconeogenesis, oxaloacetate => fructose-6P-Semi-phosphorylative Entner-Doudoroff pathway, gluconate => glycerate-3P-D-galactonate degradation, De Ley-Doudoroff pathway, D-galactonate => glycerate-3P-Reductive pentose phosphate cycle (Calvin cycle)-Reductive pentose phosphate cycle, ribulose-5P => glyceraldehyde-3P-Oxygenic photosynthesis in plants and cyanobacteria-Anoxygenic photosynthesis in purple bacteria | 3.92E-08 | M00001-M00002-M00003-M00308-M00552-M00165-M00166-M00611-M00612 |
| Pentose phosphate pathway (Pentose phosphate cycle)-Pentose phosphate pathway, non-oxidative phase, fructose 6P => ribose 5P-Reductive pentose phosphate cycle (Calvin cycle)-Reductive pentose phosphate cycle, glyceraldehyde-3P => ribulose-5P-Oxygenic photosynthesis in plants and cyanobacteria-Anoxygenic photosynthesis in purple bacteria | 3.92E-08 | M00004-M00007-M00165-M00167-M00611-M00612 |
| Inosine monophosphate biosynthesis, PRPP + glutamine => IMP-Adenine ribonucleotide biosynthesis, IMP => ADP,ATP | 1.73E-07 | M00048-M00049 |
| Phosphate acetyltransferase-acetate kinase pathway, acetyl-CoA => acetate-Methanogenesis, acetate => methane-Methanogen-Acetogen | 8.13E-07 | M00579-M00357-M00617-M00618 |
| GABA (gamma-Aminobutyrate) shunt | 6.40E-05 | M00027 |
| Citrate cycle (TCA cycle, Krebs cycle)-Citrate cycle, second carbon oxidation, 2-oxoglutarate => oxaloacetate-Reductive citrate cycle (Arnon-Buchanan cycle)-Succinate dehydrogenase, prokaryotes-Anoxygenic photosynthesis in green nonsulfur bacteria-Anoxygenic photosynthesis in green sulfur bacteria | 9.23E-05 | M00009-M00011-M00173-M00149-M00613-M00614 |
| Isoleucine biosynthesis, pyruvate => 2-oxobutanoate-Leucine biosynthesis, 2-oxoisovalerate => 2-oxoisocaproate | 0.0015 | M00535-M00432 |
| **Modules significantly enriched in the MASA10 pangenome group.** | | |
| Cobalamin biosynthesis, anaerobic, uroporphyrinogen III => sirohydrochlorin => cobyrinate a,c-diamide-Cobalamin biosynthesis, aerobic, uroporphyrinogen III => precorrin 2 => cobyrinate a,c-diamide | 3.92E-08 | M00924-M00925 |
| Pentose phosphate pathway (Pentose phosphate cycle)-Pentose phosphate pathway, non-oxidative phase, fructose 6P => ribose 5P-Pentose phosphate pathway, archaea, fructose 6P => ribose 5P-Reductive pentose phosphate cycle (Calvin cycle)-Reductive pentose phosphate cycle, glyceraldehyde-3P => ribulose-5P-Oxygenic photosynthesis in plants and cyanobacteria-Anoxygenic photosynthesis in purple bacteria | 3.92E-08 | M00004-M00007-M00580-M00165-M00167-M00611-M00612 |
| Phosphatidylcholine (PC) biosynthesis, PE => PC | 3.92E-08 | M00091 |
| Assimilatory sulfate reduction, sulfate => H2S-Dissimilatory sulfate reduction, sulfate => H2S-Sulfate-sulfur assimilation | 1.73E-07 | M00176-M00596-M00616 |
| Catechol meta-cleavage, catechol => acetyl-CoA / 4-methylcatechol => propanoyl-CoA | 1.73E-07 | M00569 |
| Isoleucine biosynthesis, threonine => 2-oxobutanoate => isoleucine | 8.13E-07 | M00570 |
| Acylglycerol degradation | 0.030088581742318 | M00098 |

**Table S8.** NCBI accession numbers of unvegetated creekbed sediment samples (PRJNA814317).

| sampleID | SRAid | metadata | Reads |
|---|---|---|---|
| G1_1_3 | SRR24130648 | Greenwood Creek, Ipswich, MA. Enriched creek (receiving input from treated sewage effluent). Increasing salinity gradient from freshwater terrestrial stream to Plum Island Sound (1-6) | 5,598,089 |
| G1_2_4 | SRR24130647 | | 7,853,810 |
| G1_3_4_1 | SRR24130636 | | 8,772,437 |
| G1_3_4_2 | SRR24130625 | | 6,054,309 |
| G2_1_2 | SRR24130622 | | 7,261,066 |
| G2_2_1 | SRR24130621 | | 8,208,631 |
| G2_3_3 | SRR24130620 | | 9,320,183 |
| G3_1_2 | SRR24130619 | | 6,326,147 |
| G3_2_1 | SRR24130618 | | 5,978,731 |
| G3_3_2 | SRR24130617 | | 6,445,823 |
| G4_1_2 | SRR24130646 | | 9,757,468 |
| G4_2_3 | SRR24130645 | | 9,156,045 |
| G4_3_1 | SRR24130644 | | 11,095,313 |
| G5_1_2 | SRR24130643 | | 5,624,988 |
| G5_2_4 | SRR24130642 | | 7,813,299 |
| G5_3_4 | SRR24130641 | | 5,898,050 |
| G6_2_4 | SRR24130640 | | 8,759,716 |
| R1_1_4 | SRR24130639 | Egypt Creek, Ipswich, MA. Reference creek (receiving input from drinking water reservoir). Increasing salinity gradient from freshwater terrestrial stream to Plum Island Sound (1-6) | 9,179,906 |
| R1_2_4 | SRR24130638 | | 3,536,787 |
| R1_3_4 | SRR24130637 | | 5,327,512 |
| R2_1_1 | SRR24130635 | | 9,563,259 |
| R2_2_1 | SRR24130634 | | 9,559,918 |
| R2_3_4 | SRR24130633 | | 10,067,392 |
| R3_1_3 | SRR24130632 | | 10,325,098 |
| R3_2_1 | SRR24130631 | | 9,799,442 |
| R3_3_3 | SRR24130630 | | 10,554,621 |
| R4_1_3 | SRR24130629 | | 10,067,285 |
| R4_2_3 | SRR24130628 | | 7,681,587 |
| R4_3_3 | SRR24130627 | | 9,884,460 |
| R6_1_2 | SRR24130626 | | 6,638,196 |
| R6_2_1 | SRR24130624 | | 10,642,403 |
| R6_3_1 | SRR24130623 | | 6,145,961 |

**Supplemental Data files description**

**Supplemental Data 1.** DRAM raw annotations.

**Supplemental Data 2.** DRAM metabolism summary.

**Supplemental Data 3.** Contig-specific coverage values of the Alabama and Massachusetts salt marsh samples.

**Supplemental Data 4.** Contig-specific coverage values of the unvegetated salt marsh samples.

**Supplemental Data 5.** Anvio profiling (read recruitment results, SNVs, indels).

**Supplemental Data 6.** Genes in MASA10 and MASP16 MAGs identified as geographically restricted.

**Supplemental Data 7.** Reference-guided metagenome reconstruction.

**Supplemental Data 8.** Enrichment analyses (MASA10, ALJR36).

**Supplemental Data 9.** Genes selected for phylogenetic tree reconstruction and average gene entropy analysis. Results of average gene entropy analysis.

**Figure S1.** Comparison of taxonomic hits at phylum-level determined with Kraken for the contigs identified in each of the co-assemblies: ALSA (Alabama, *S. alterniflorus*), ALJR (Alabama, *J. roemerianus*), MASP (Massachusetts, *S. pumilus*), and MASA (Massachusetts, *S. alterniflorus*).

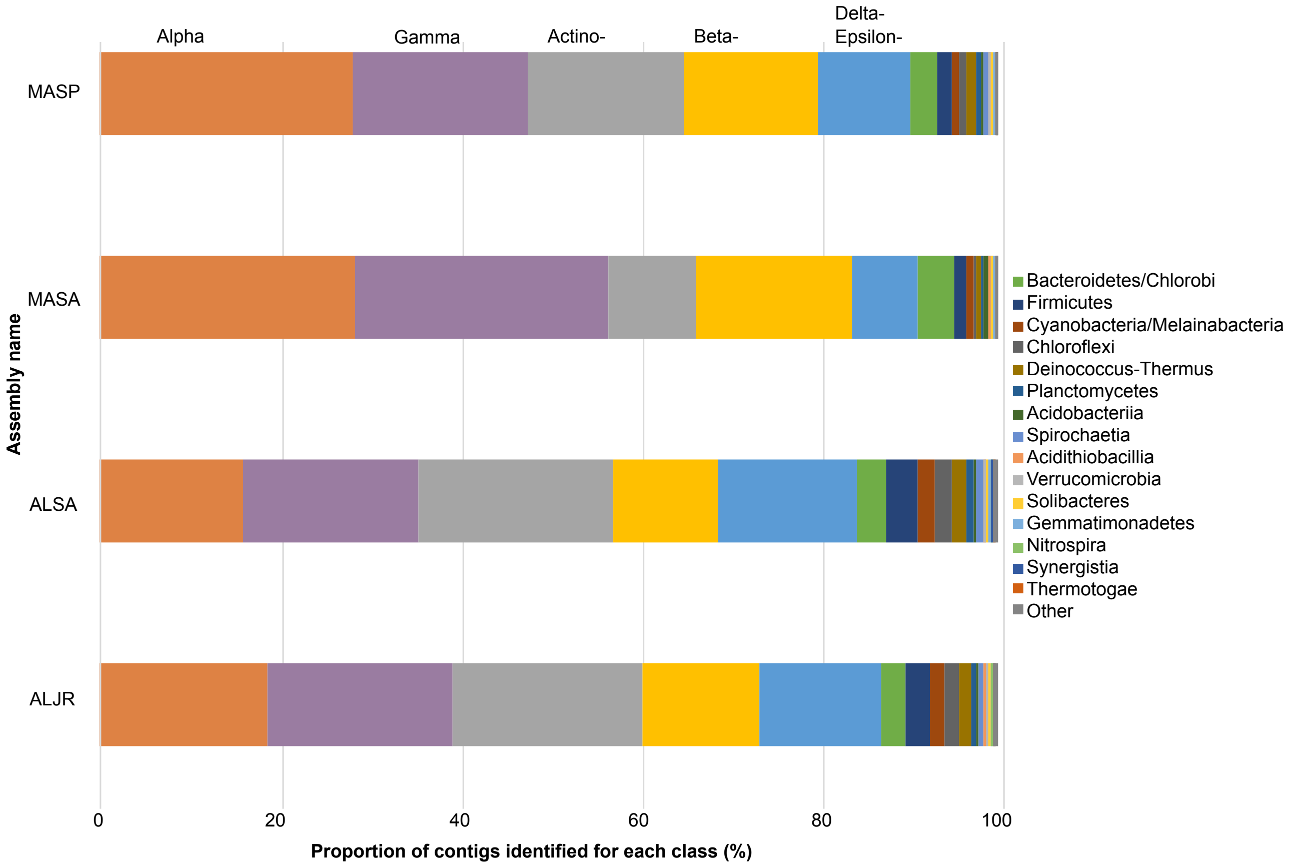

**Figure S2.** Primary MAGs for which read-mapping was statistically more abundant across locations and vegetation zones by pairwise statistical analysis (STAMP) using average CPM values calculated for each primary MAG. Mean proportion of reads (and pairwise differences) are shown with confidence intervals. A) Comparison by location and vegetation: Massachusetts- *S. pumilus* (MASP), Massachusetts- *S. alterniflorus* (MASA), Alabama- *S. alterniflorus* (ALSA), Alabama- *J. roemerianus* (ALJR). B) Comparison by location (Alabama vs. Massachusetts).

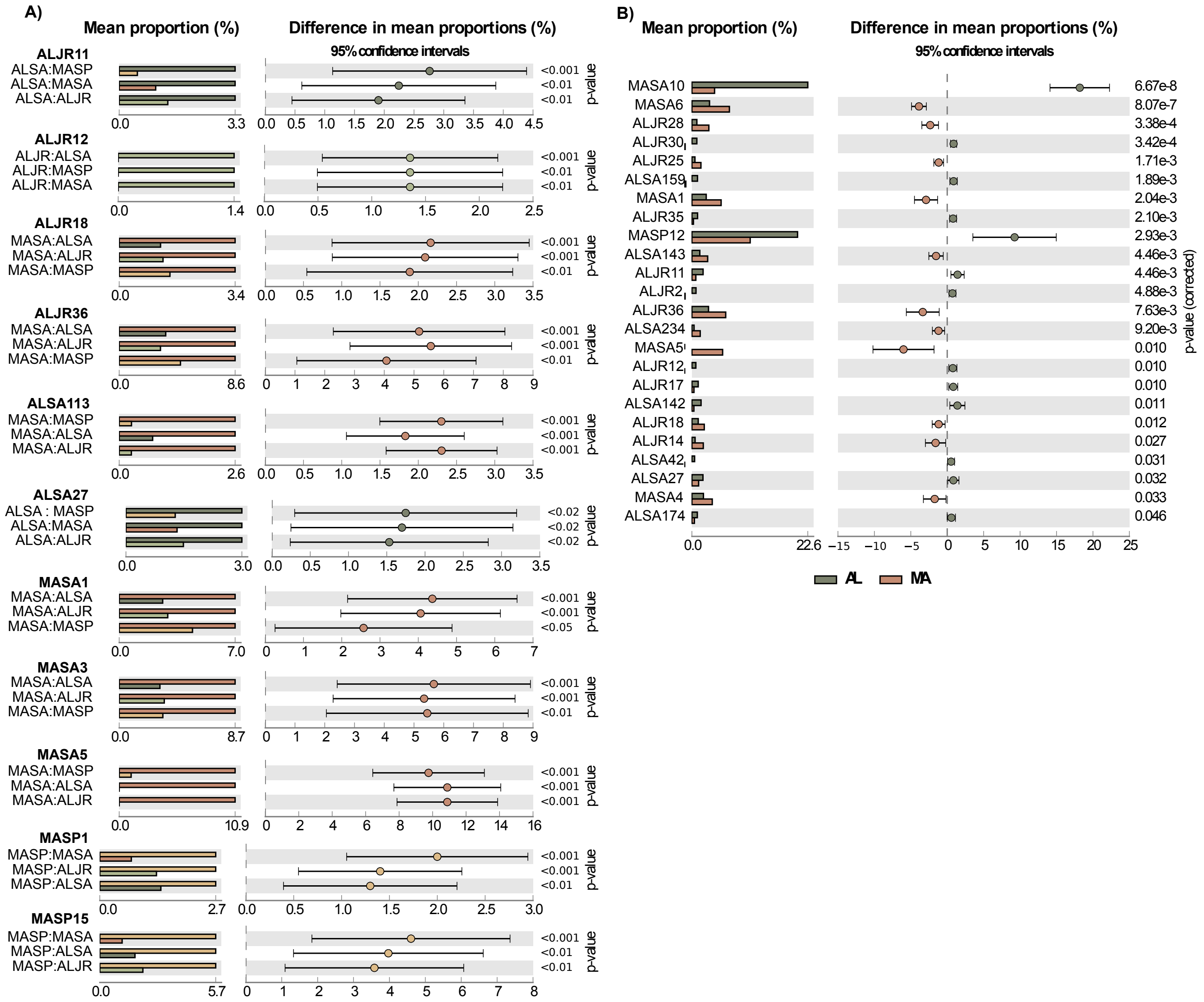

**Figure S3.** Contig-specific coverage values for each salt marsh sample included in this study of each sulfur cycling bacteria MAG identified in the co-assembly from Alabama in sediments inhabited by the plant *Juncus roemerianus* (ALJR). For each sample, contig coverage (expressed as CMP, y-axis) is sorted from low to high. For comparison purposes, samples collected in the same geographical area and in the proximity of the same plant are colored with the same color. Full data is presented in Supplemental Data 3.

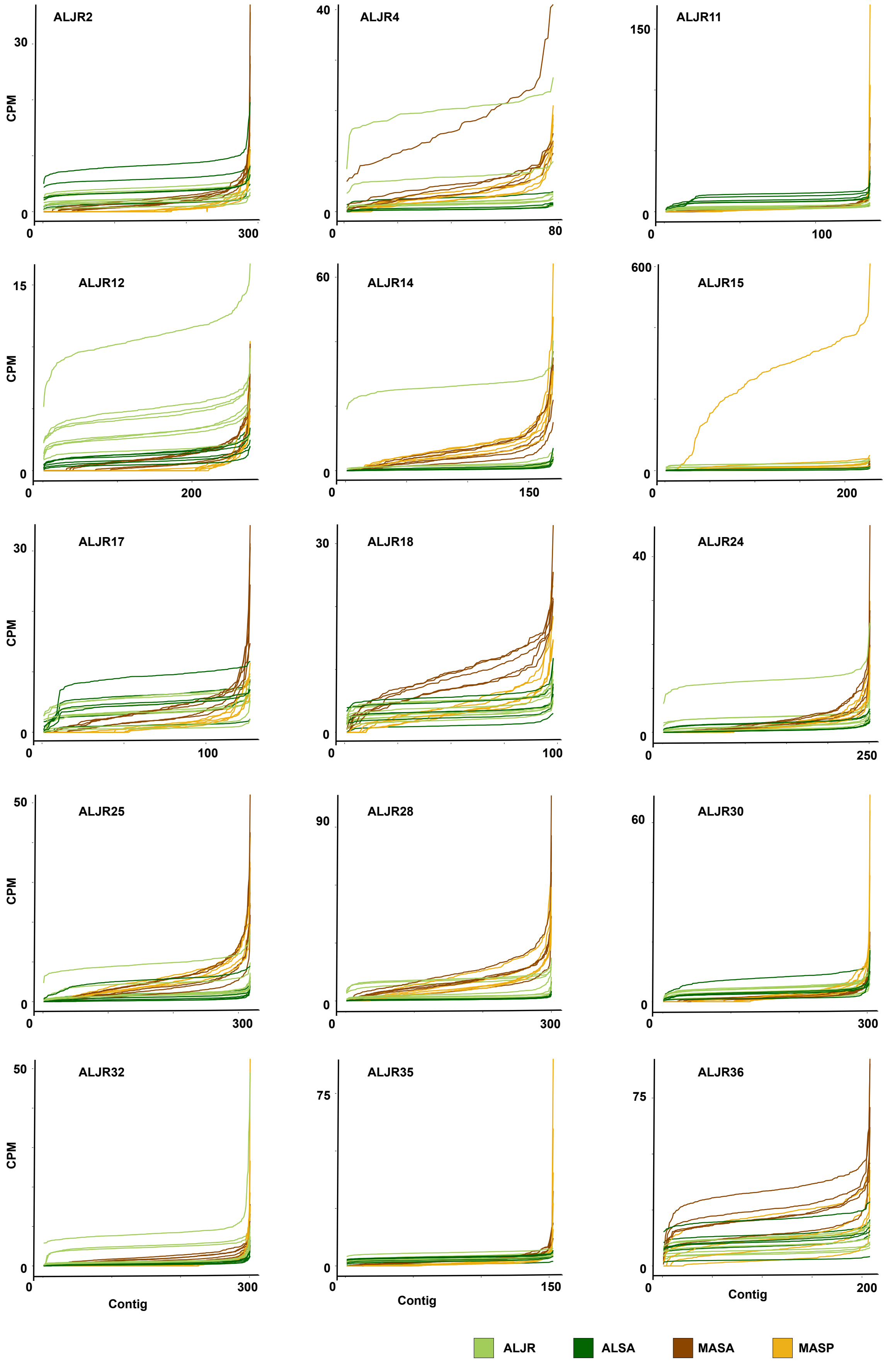

**Figure S4.** Contig-specific coverage values for each salt marsh sample included in this study of each sulfur cycling bacteria MAG identified in the co-assembly from Alabama in sediments inhabited by the plant *Sporobulus alterniflorus* (ALSA). For each sample, contig coverage (expressed as CMP, y-axis) is sorted from low to high. For comparison purposes, samples collected in the same geographical area and in the proximity of the same plant are colored with the same color. Full data is presented in Supplemental Data 3.

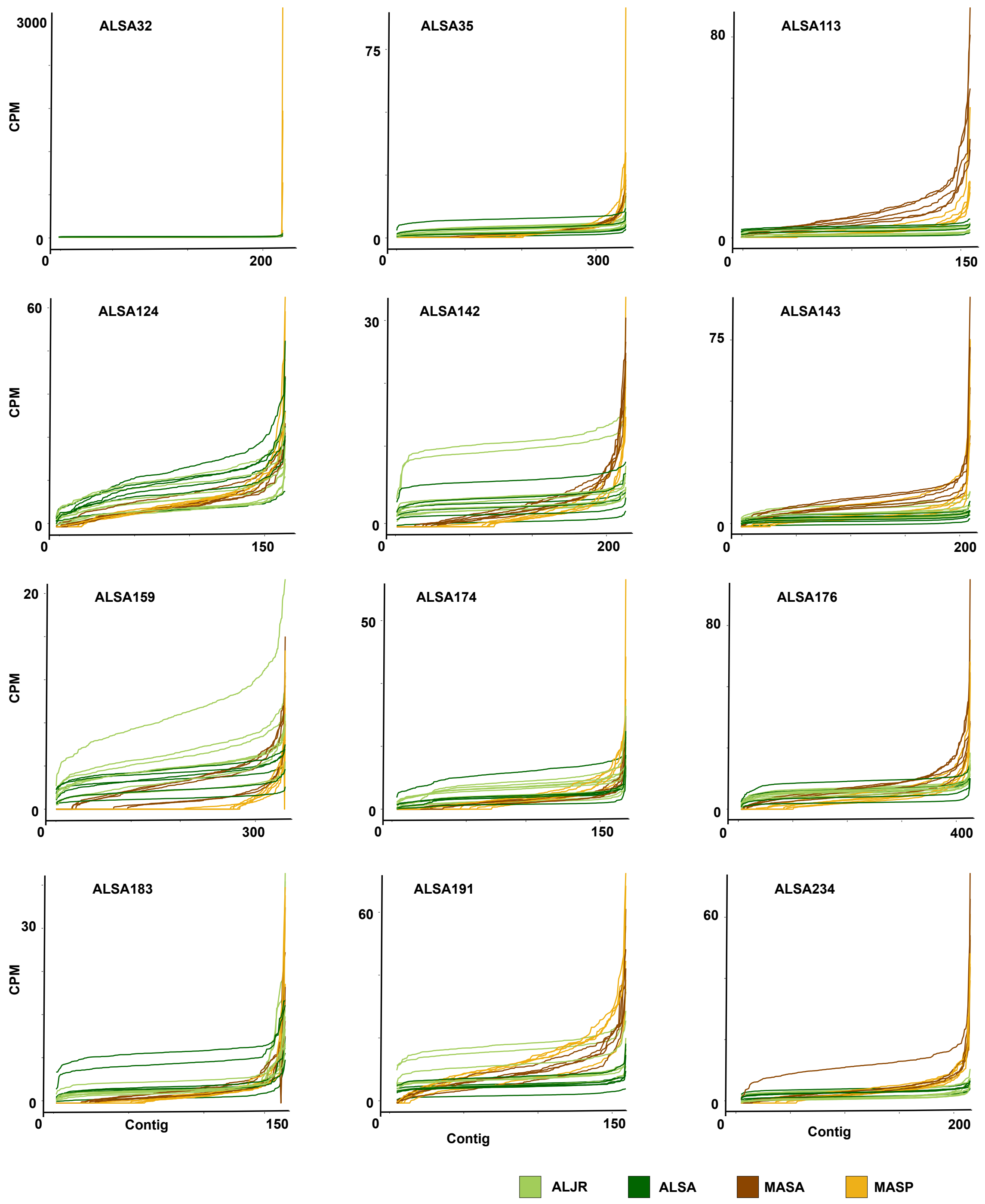

**Figure S5.** Contig-specific coverage values for each salt marsh sample included in this study of each sulfur cycling bacteria MAG identified in the co-assembly from Massachusetts in sediments inhabited by *Sporobolus alterniflorus* (MASA) or *Sporobolus pumilus* (MASP). For each sample, contig coverage (expressed as CMP, y-axis) is sorted from low to high. For comparison purposes, samples collected in the same geographical area and in the proximity of the same plant are colored with the same color. Full data is presented in Supplemental Data 3.

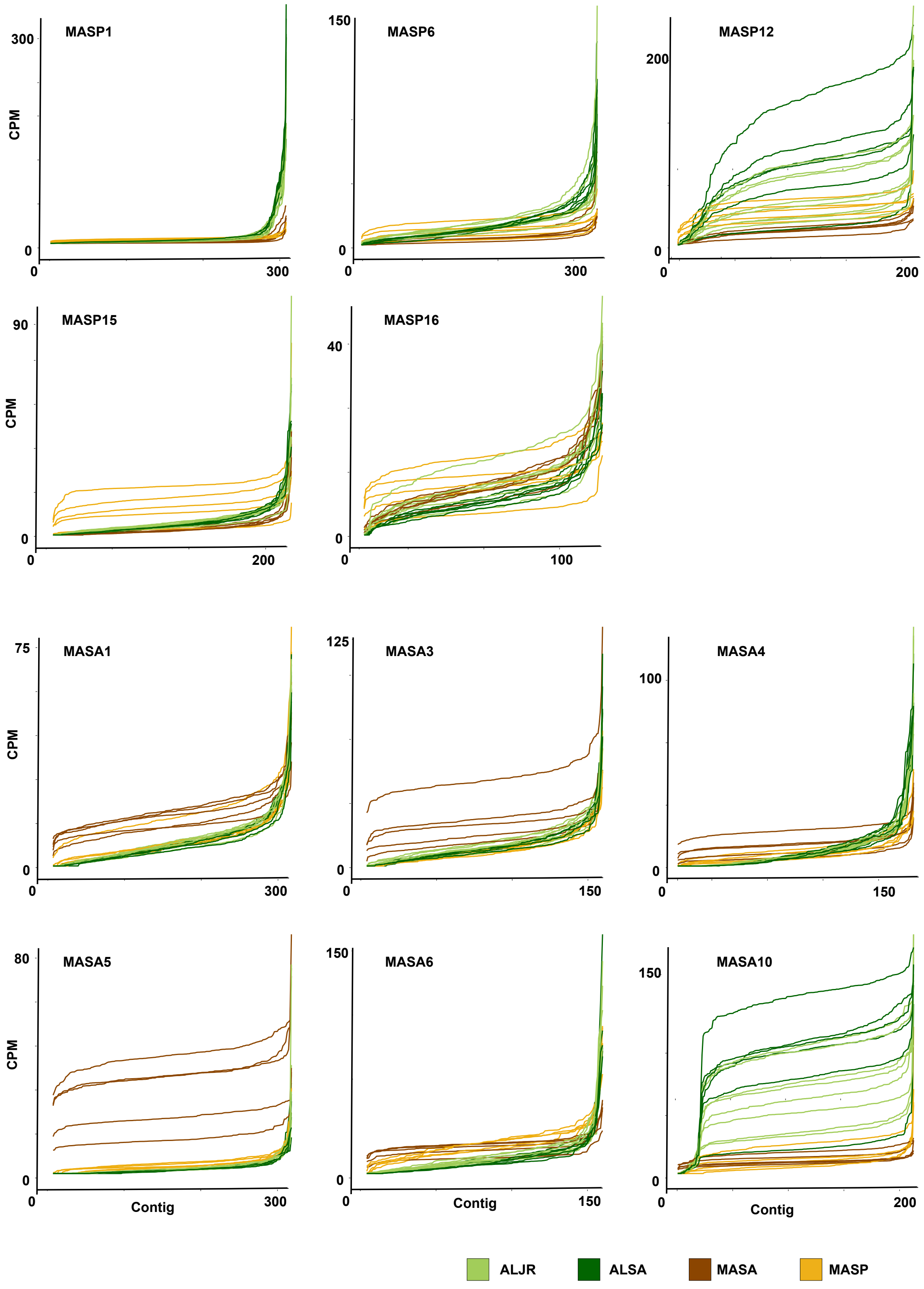

**Figure S6.** Distribution of sulfur cycling primary MAG sequences across environmental metagenomes. Samples included published metagenomes from subtidal sediments along a salinity gradient. Circle size reflects averaged CPM across all contigs for each MAG. Contig-specific coverage values can be found in Supplemental Data 4 and sample description in Table S8.

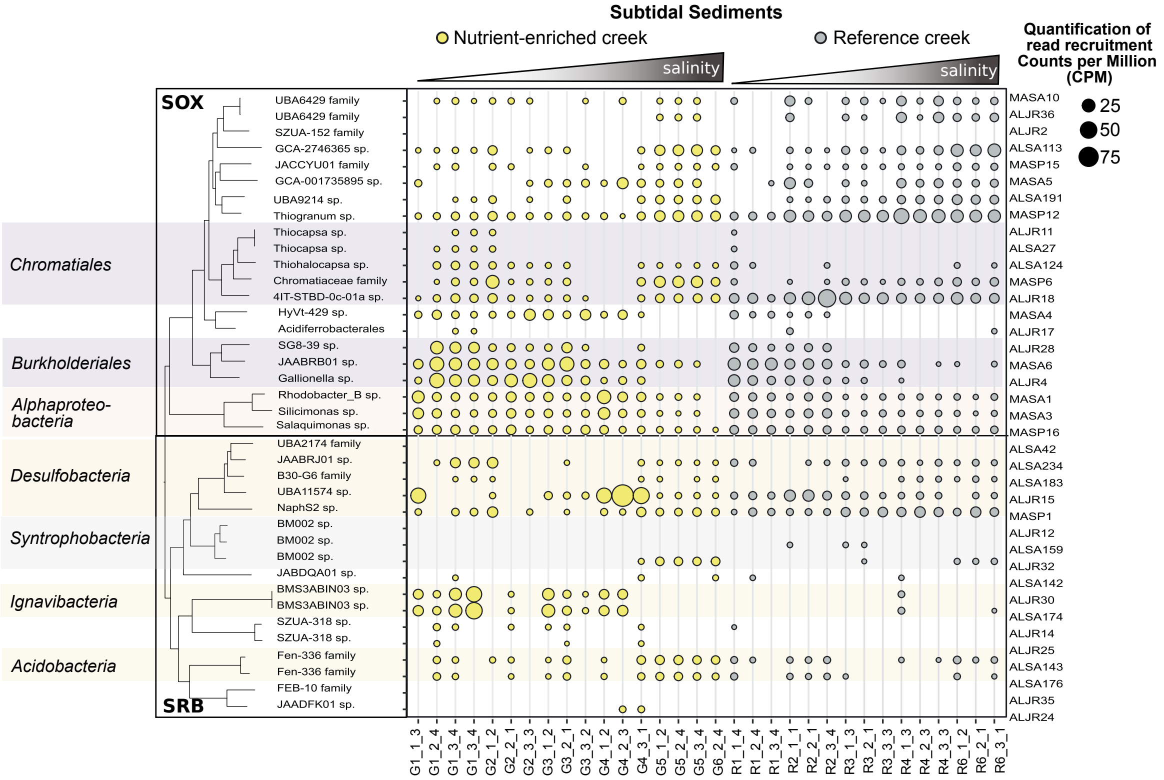

**Figure S7.** Distribution, across the 24 samples, of eight sulfur cycling primary MAG sequences selected for further analyses. A) Detail of Fig. 4a highlighting the MAGs used for the Anvi'o-based analysis. Quantification was by read-mapping from AL and MA samples to primary MAG contigs, with coverage standardized by library size and contig length and expressed as counts per million reads (CPM). B) Results of Bowtie read recruitment expressed as million reads mapped (left) and the percentage of reads of a given metagenomic sample recruited to each of the MAGs (right). C) Example relationships between sampling depth (as the number of reads recruited after mapping) and the number of detected SNVs for two of the eight MAGs. Legend as for panel A) (Additional graphs are presented in Supplemental Data 6).

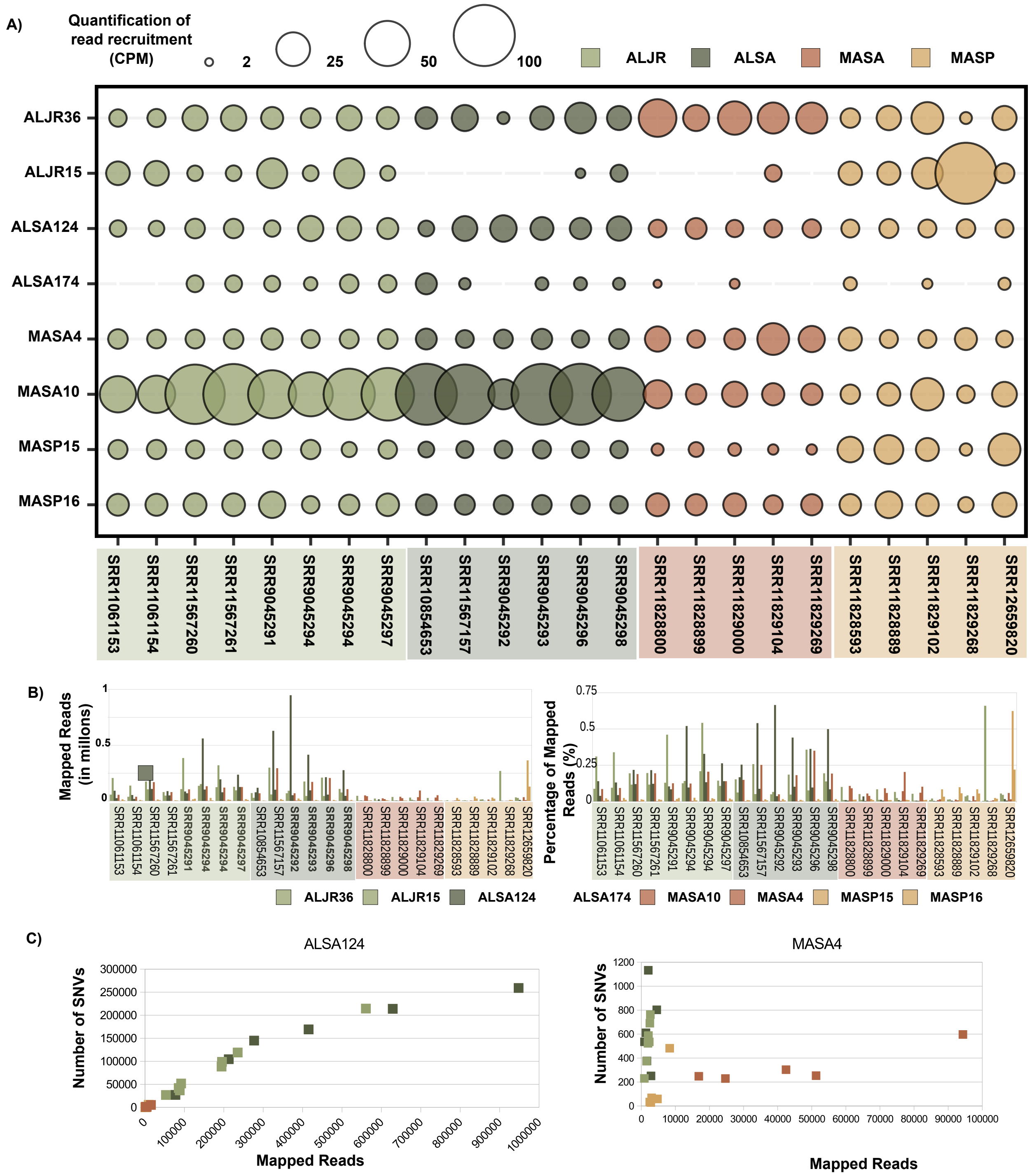

**Figure S8-S15. Top.** Anvi'o coverage profiles of eight selected sulfur-cycling bacteria MAGs. In each anviogram, outer cycles represent the contig coverage (as mean coverage) of each of the 24 salt-marsh metagenomic samples included in this analysis. Samples are color-coded according to geographical site and the dominant plant species in the area where sediment collection occurred. Inner rings represent GC content (dark gray) and contig length (gray). Contigs are clustered (inner tree) based on the sequence composition and differential coverage using Euclidean distance and Ward hierarchical clustering method. The samples' order (rings) was determined using a clustering method based on the mean coverage. Bar plots represent (top to bottom) the total number of reads in each library, the total number of reads mapped to each respective MAG, the percentage of mapped reads, and the total number of Single Nucleotide Variants (SNV), Single Codon Variants (SCV) and indels. **Bottom.** Anvi'o pangenomic analysis of eight sulfur-cycling bacteria MAGs. In each ring, each vertical line represents a gene. The genes' order was determined using Euclidean distance and Ward clustering method based on the presence or absence of a gene cluster across all MAGs. In the outer rings, we indicate the number of genes in each gene cluster, the number of paralogous, and the results of the combined homogeneity test. Dark gray bars in the next set of rings indicate a known gene function, annotated according to COG 2020. The single-copy core gene cluster, present in all reference-guided reassembled MAGs, is highlighted in purple. The primary MAG used as a reference is shown in dark teal. Each of the remaining circles represents a reference-guided reassembled MAG from each of the salt-mash metagenomic samples included in this analysis. Samples are color-coded according to geographical site and the dominant plant species in the area where sediment collection occurs.

### Figure S8

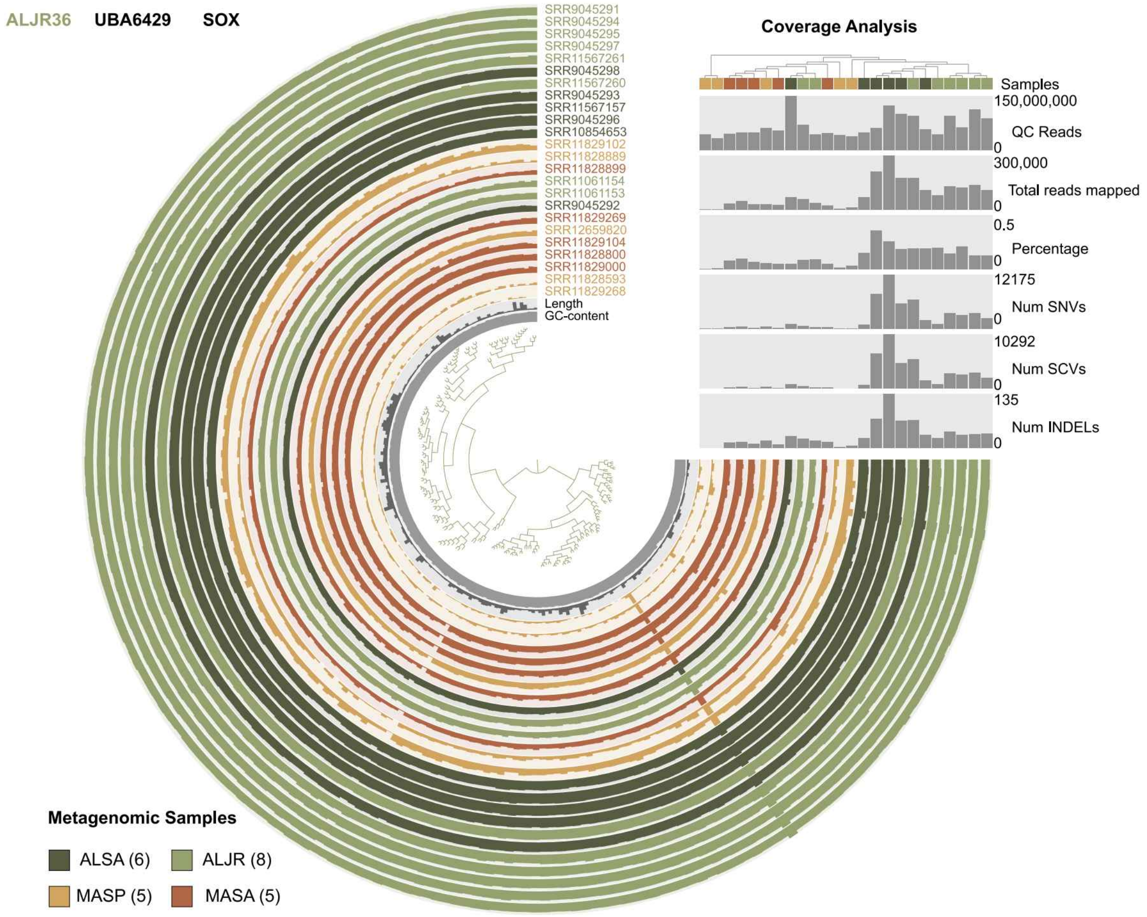

ALJR36 UBA6429 SOX
Coverage Analysis
SRR9045291
SRR9045294
SRR9045295
SRR9045297
SRR11567261
SRR9045298
SRR11567260
SRR9045293
SRR11567157
SRR9045296
SRR10854653
SRR11829102
SRR11828889
SRR11828899
SRR11061154
SRR11061153
SRR9045292
SRR11829269
SRR12659820
SRR11829104
SRR11828800
SRR11829000
SRR11828593
SRR11829268
Length
GC-content
Samples
150,000,000
QC Reads
0
300,000
Total reads mapped
0
0.5
Percentage
0
12175
Num SNVs
0
10292
Num SCVs
0
135
Num INDELs
0
Metagenomic Samples
ALSA (6)
ALJR (8)
MASP (5)
MASA (5)


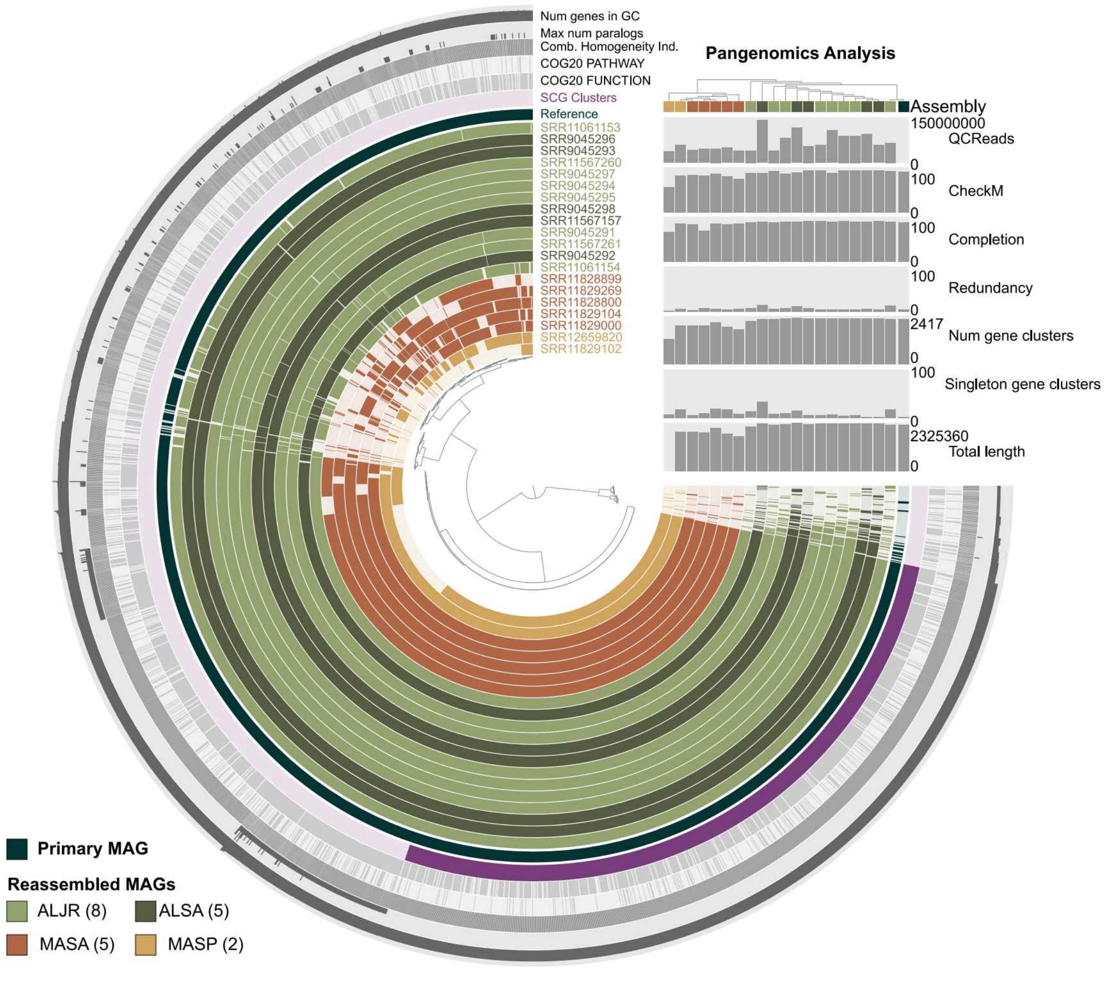

Num genes in GC
Max num paralogs
Comb. Homogeneity Ind.
COG20 PATHWAY
COG20 FUNCTION
SCG Clusters
Reference
Pangenomics Analysis
Assembly
SRR11061153
SRR9045296
SRR9045293
SRR11567260
SRR9045297
SRR9045294
SRR9045295
SRR9045298
SRR11567157
SRR9045291
SRR11567261
SRR9045292
SRR11061154
SRR11828899
SRR11829269
SRR11828800
SRR11829104
SRR11829000
SRR12659820
SRR11829102
150000000
QCReads
0
100
CheckM
0
100
Completion
0
100
Redundancy
0
2417
Num gene clusters
0
100
Singleton gene clusters
0
2325360
Total length
0
Primary MAG
Reassembled MAGs
ALJR (8)
ALSA (5)
MASA (5)
MASP (2)

#### Figure S9

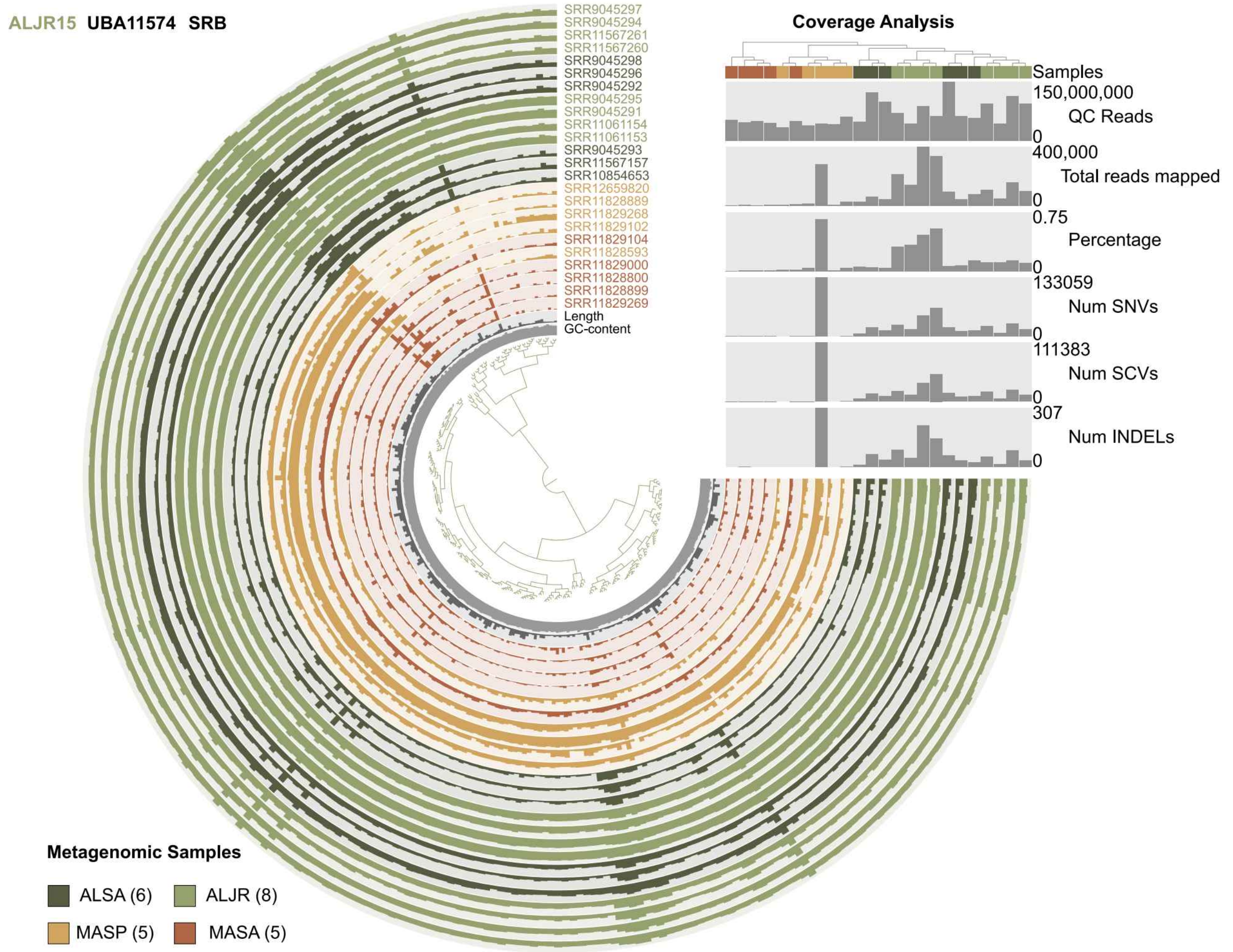
ALJR15 UBA11574 SRB
Coverage Analysis
SRR9045297
SRR9045294
SRR11567261
SRR11567260
SRR9045298
SRR9045296
SRR9045292
SRR9045295
SRR9045291
SRR11061154
SRR11061153
SRR9045293
SRR11567157
SRR10854653
SRR12659820
SRR11828889
SRR11829268
SRR11829102
SRR11829104
SRR11828593
SRR11829000
SRR11828800
SRR11828899
SRR11829269
Length
GC-content
Samples
150,000,000
QC Reads
0
400,000
Total reads mapped
0
0.75
Percentage
0
133059
Num SNVs
0
111383
Num SCVs
0
307
Num INDELs
0
Metagenomic Samples
ALSA (6)
ALJR (8)
MASP (5)
MASA (5)

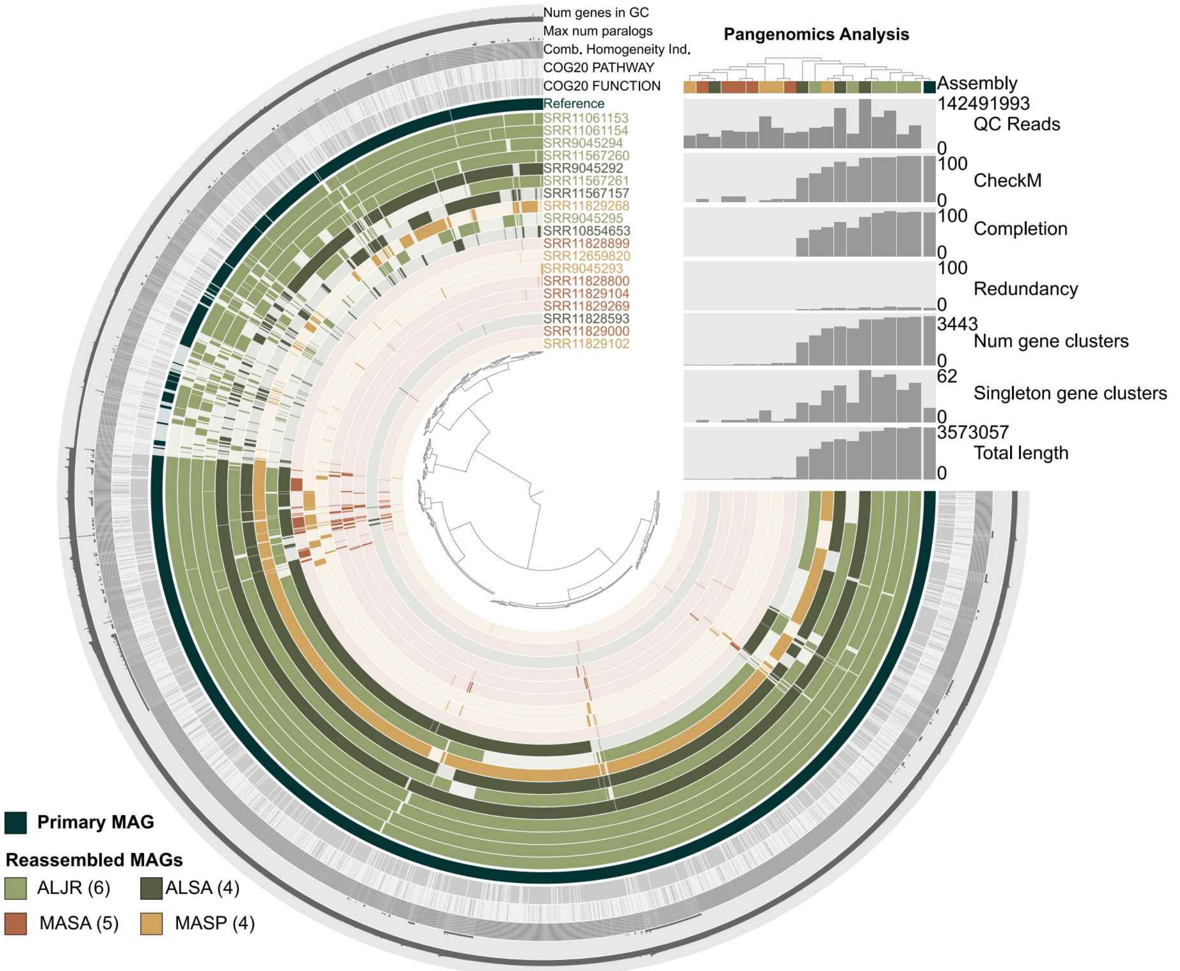
Num genes in GC
Max num paralogs
Comb. Homogeneity Ind.
COG20 PATHWAY
COG20 FUNCTION
Reference
SRR11061153
SRR11061154
SRR9045294
SRR11567260
SRR9045292
SRR11567261
SRR11567157
SRR11829268
SRR9045295
SRR10854653
SRR11828899
SRR12659820
SRR9045293
SRR11828800
SRR11829104
SRR11829269
SRR11828593
SRR11829000
SRR11829102
Pangenomics Analysis
Assembly
142491993
QC Reads
0
100
CheckM
0
100
Completion
0
100
Redundancy
0
3443
Num gene clusters
0
62
Singleton gene clusters
0
3573057
Total length
0
Primary MAG
Reassembled MAGs
ALJR (6)
ALSA (4)
MASA (5)
MASP (4)

### Figure S10

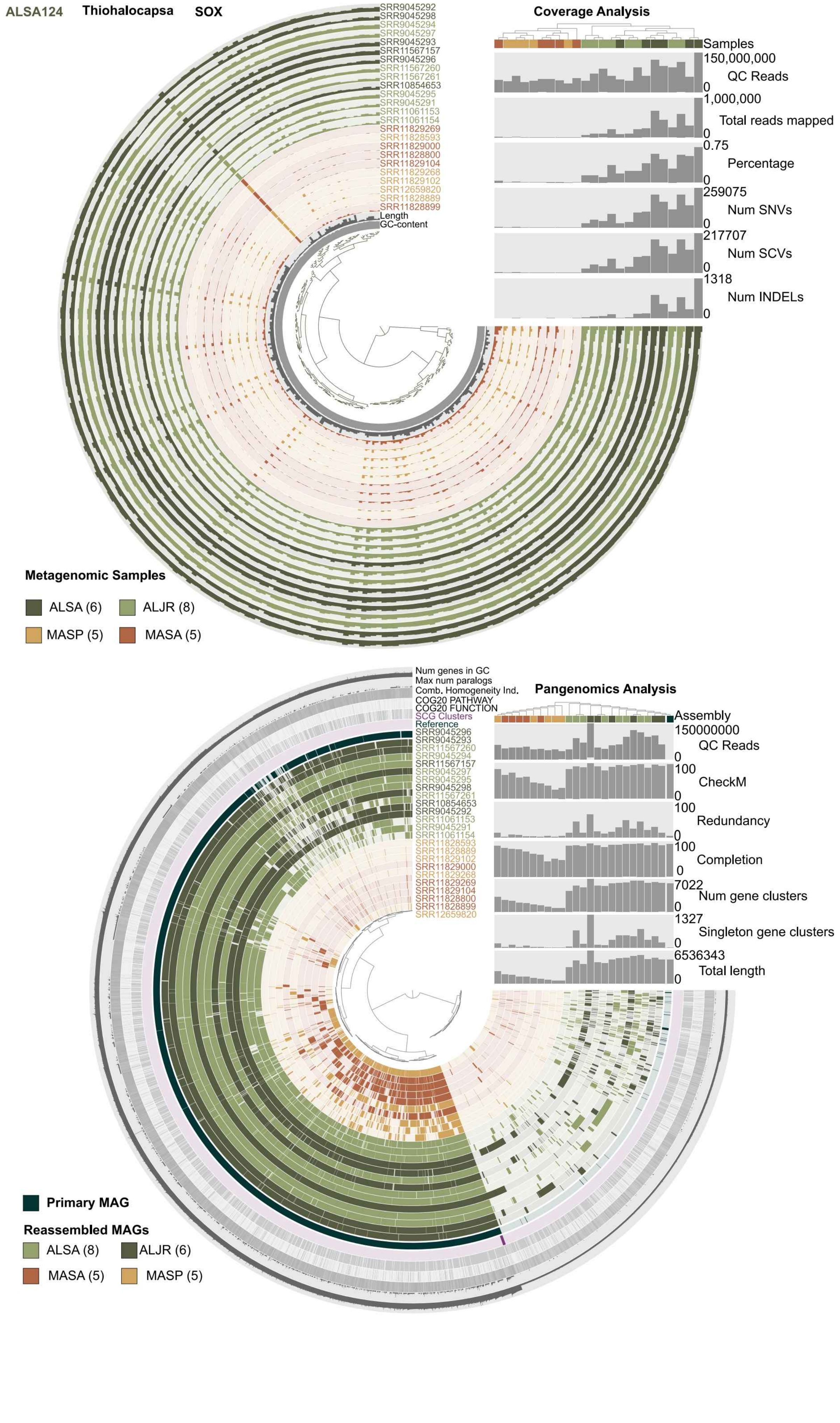
ALSA124
Thiohalocapsa
SOX
SRR9045292
SRR9045298
SRR9045294
SRR9045297
SRR9045293
SRR11567157
SRR9045296
SRR11567260
SRR11567261
SRR10854653
SRR9045295
SRR9045291
SRR11061153
SRR11061154
SRR11829269
SRR11828593
SRR11829000
SRR11828800
SRR11829104
SRR11829268
SRR11829102
SRR12659820
SRR11828889
SRR11828899
Length
GC-content
Coverage Analysis
Samples
150,000,000
QC Reads
0
1,000,000
Total reads mapped
0
0.75
Percentage
0
259075
Num SNVs
0
217707
Num SCVs
0
1318
Num INDELs
0
Metagenomic Samples
ALSA (6)
ALJR (8)
MASP (5)
MASA (5)
Num genes in GC
Max num paralogs
Comb. Homogeneity Ind.
COG20 PATHWAY
COG20 FUNCTION
SCG Clusters
Reference
SRR9045296
SRR9045293
SRR11567260
SRR9045294
SRR11567157
SRR9045297
SRR9045295
SRR9045298
SRR11567261
SRR10854653
SRR9045292
SRR11061153
SRR9045291
SRR11061154
SRR11828593
SRR11828889
SRR11829102
SRR11829000
SRR11829268
SRR11829269
SRR11829104
SRR11828800
SRR11828899
SRR12659820
Pangenomics Analysis
Assembly
150000000
QC Reads
0
100
CheckM
0
100
Redundancy
0
100
Completion
0
7022
Num gene clusters
0
1327
Singleton gene clusters
0
6536343
Total length
0
Primary MAG
Reassembled MAGs
ALSA (8)
ALJR (6)
MASA (5)
MASP (5)

### Figure S11

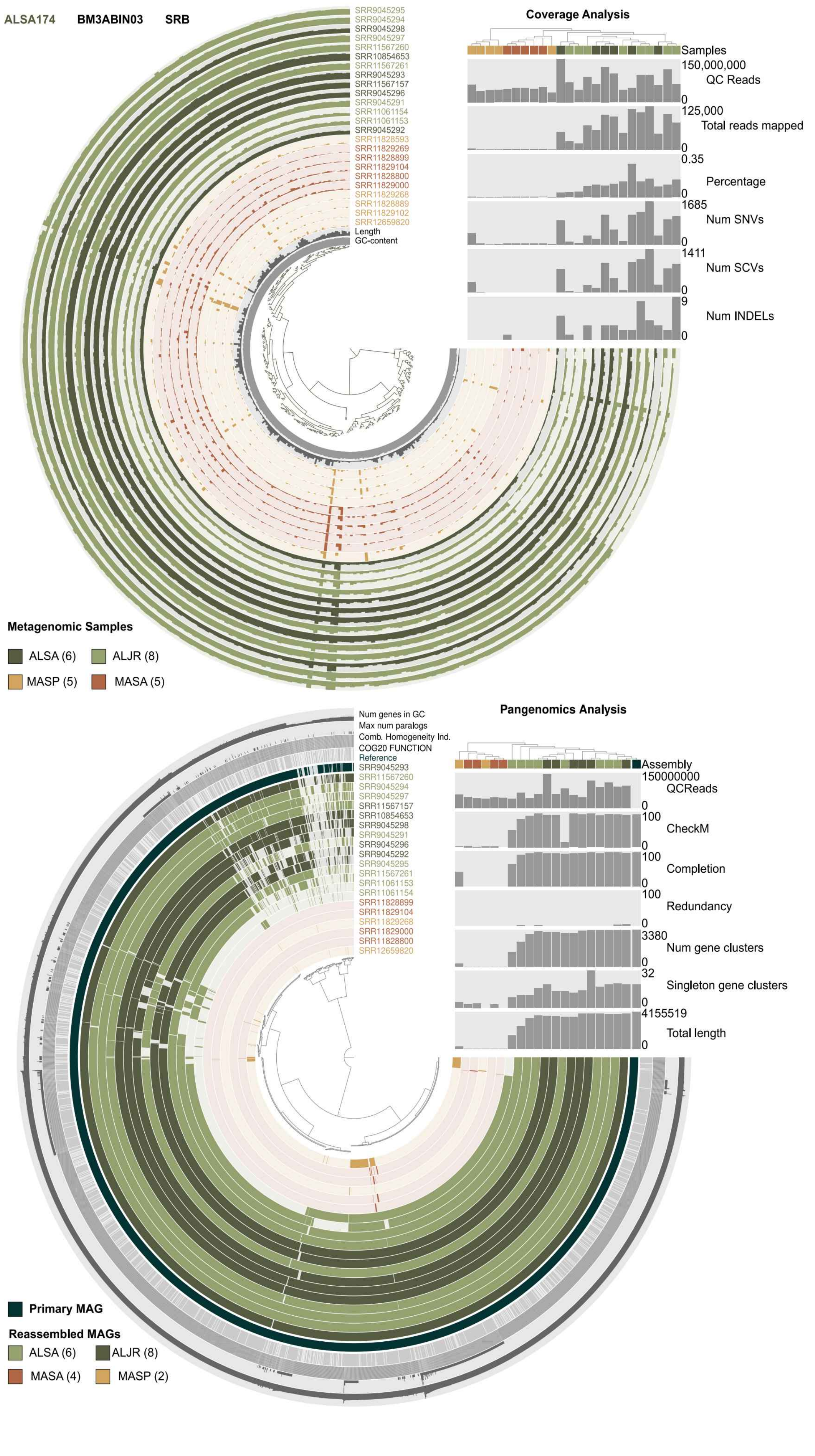

ALSA174
BM3ABIN03
SRB
SRR9045295
SRR9045294
SRR9045298
SRR9045297
SRR11567260
SRR10854653
SRR11567261
SRR9045293
SRR11567157
SRR9045296
SRR9045291
SRR11061154
SRR11061153
SRR9045292
SRR11828593
SRR11829269
SRR11828899
SRR11829104
SRR11828800
SRR11829000
SRR11829268
SRR11828889
SRR11829102
SRR12659820
Length
GC-content
Coverage Analysis
Samples
150,000,000
QC Reads
0
125,000
Total reads mapped
0
0.35
Percentage
0
1685
Num SNVs
0
1411
Num SCVs
0
9
Num INDELs
0
Metagenomic Samples
ALSA (6)
ALJR (8)
MASP (5)
MASA (5)
Num genes in GC
Max num paralogs
Comb. Homogeneity Ind.
COG20 FUNCTION
Reference
SRR9045293
SRR11567260
SRR9045294
SRR9045297
SRR11567157
SRR10854653
SRR9045298
SRR9045291
SRR9045296
SRR9045292
SRR9045295
SRR11567261
SRR11061153
SRR11061154
SRR11828899
SRR11829104
SRR11829268
SRR11829000
SRR11828800
SRR12659820
Pangenomics Analysis
Assembly
150000000
QCReads
0
100
CheckM
0
100
Completion
0
100
Redundancy
0
3380
Num gene clusters
0
32
Singleton gene clusters
0
4155519
Total length
0
Primary MAG
Reassembled MAGs
ALSA (6)
ALJR (8)
MASA (4)
MASP (2)

**Figure S12**

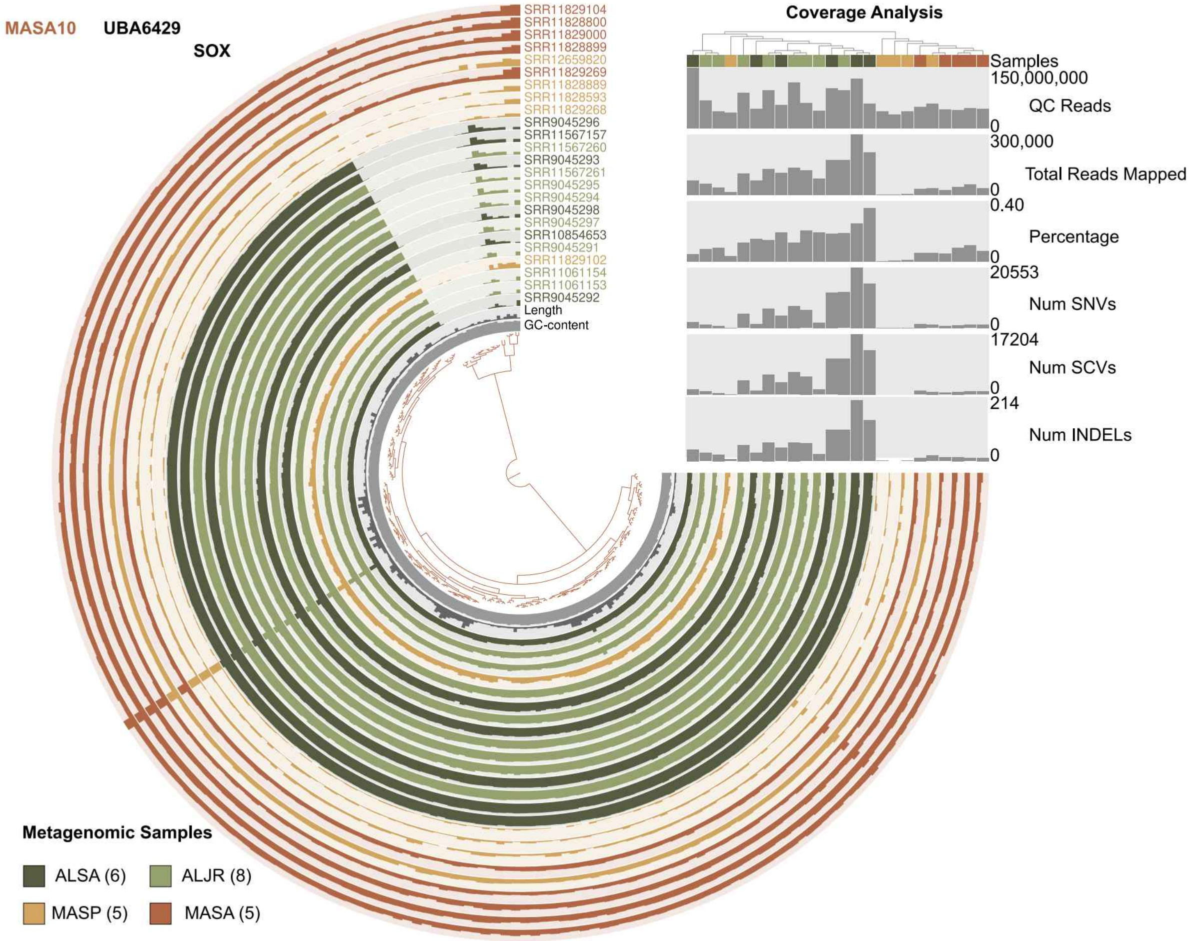

MASA10
UBA6429
SOX
SRR11829104
SRR11828800
SRR11829000
SRR11828899
SRR12659820
SRR11829269
SRR11828889
SRR11828593
SRR11829268
SRR9045296
SRR11567157
SRR11567260
SRR9045293
SRR11567261
SRR9045295
SRR9045294
SRR9045298
SRR9045297
SRR10854653
SRR9045291
SRR11829102
SRR11061154
SRR11061153
SRR9045292
Length
GC-content
Coverage Analysis
Samples
150,000,000
QC Reads
0
300,000
Total Reads Mapped
0
0.40
Percentage
0
20553
Num SNVs
0
17204
Num SCVs
0
214
Num INDELs
0
Metagenomic Samples
ALSA (6)
ALJR (8)
MASP (5)
MASA (5)


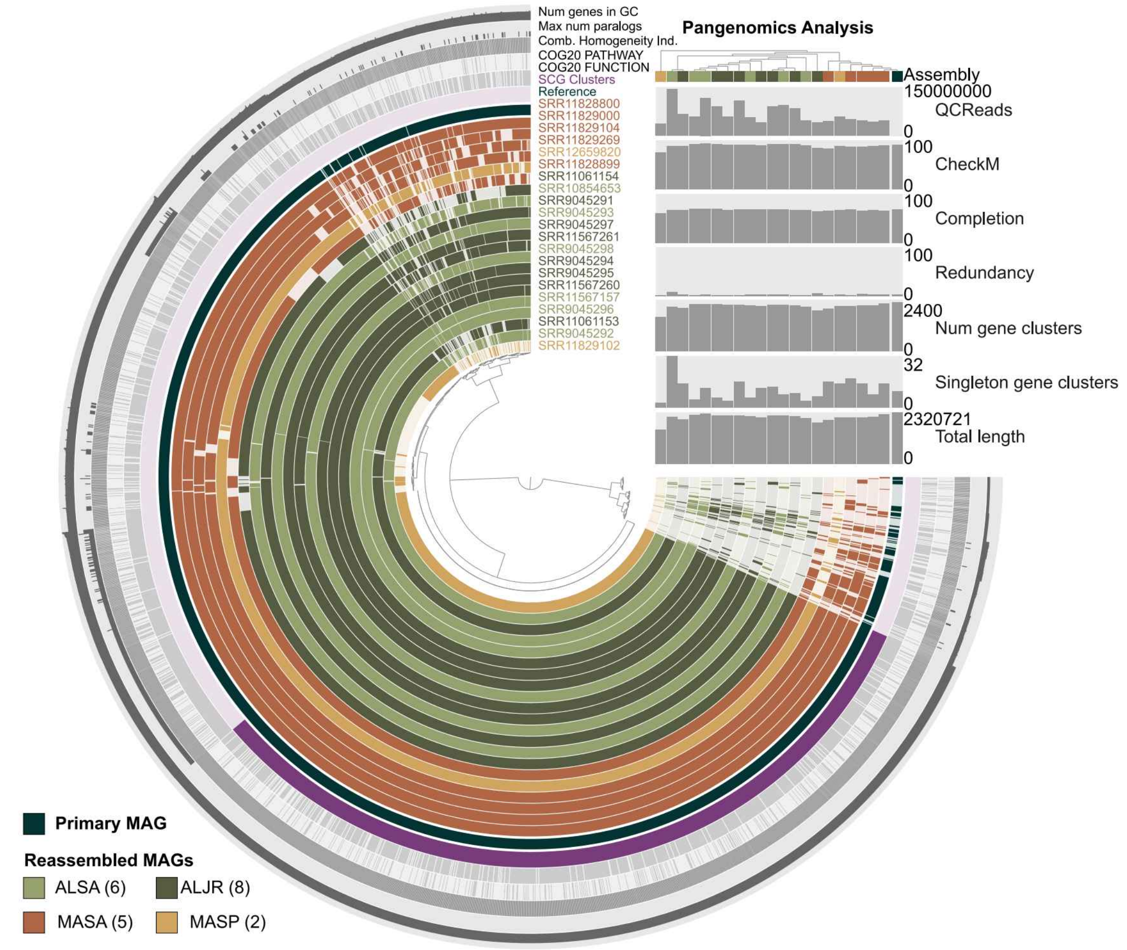

Num genes in GC
Max num paralogs
Comb. Homogeneity Ind.
COG20 PATHWAY
COG20 FUNCTION
SCG Clusters
Reference
SRR11828800
SRR11829000
SRR11829104
SRR11829269
SRR12659820
SRR11828899
SRR11061154
SRR10854653
SRR9045291
SRR9045293
SRR9045297
SRR11567261
SRR9045298
SRR9045294
SRR9045295
SRR11567260
SRR11567157
SRR9045296
SRR11061153
SRR9045292
SRR11829102
Pangenomics Analysis
Assembly
150000000
QCReads
0
100
CheckM
0
100
Completion
0
100
Redundancy
0
2400
Num gene clusters
0
32
Singleton gene clusters
0
2320721
Total length
0
Primary MAG
Reassembled MAGs
ALSA (6)
ALJR (8)
MASA (5)
MASP (2)

### Figure S13

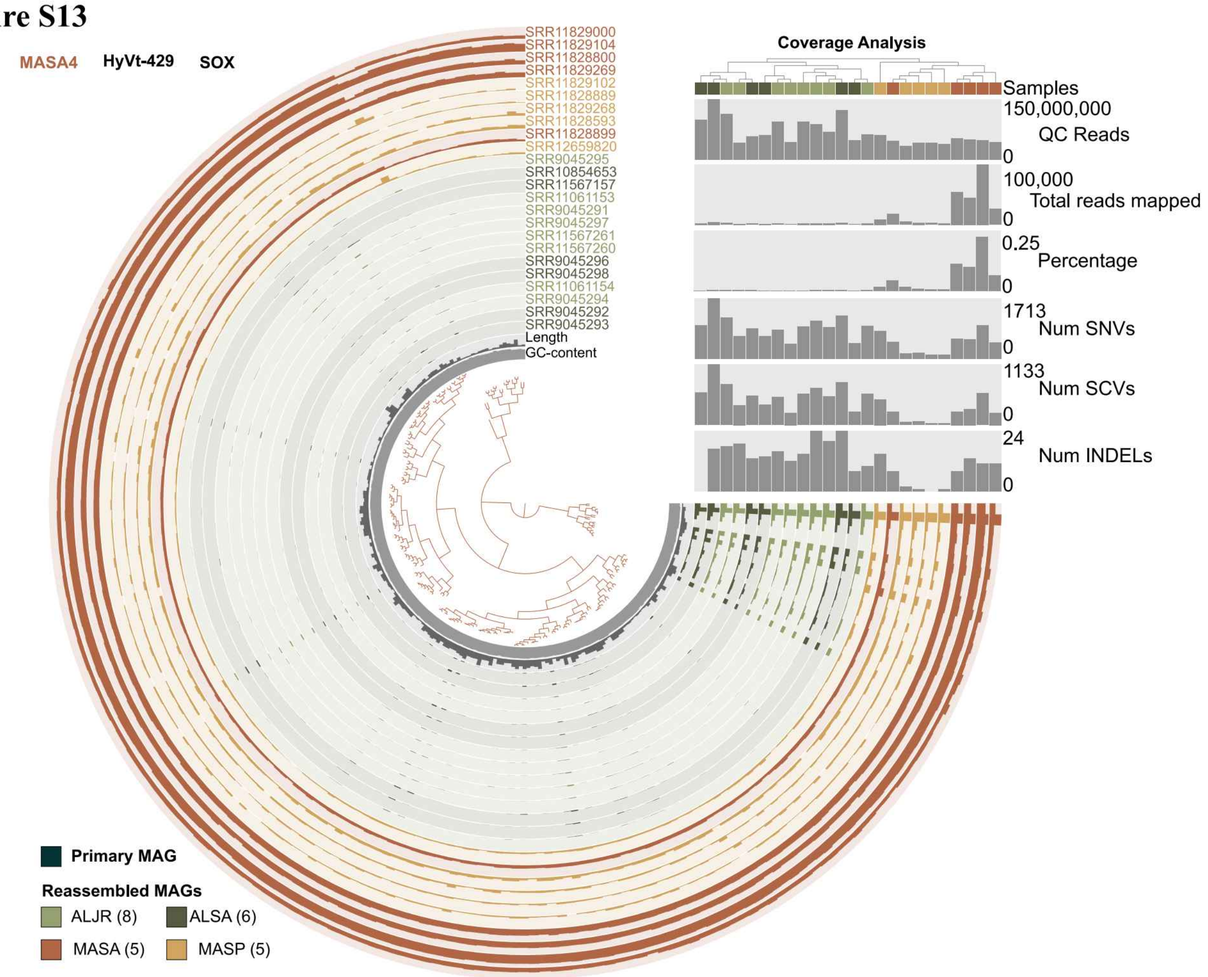

MASA4
HyVt-429
SOX
Coverage Analysis
SRR11829000
SRR11829104
SRR11828800
SRR11829269
SRR11829102
SRR11828889
SRR11829268
SRR11828593
SRR11828899
SRR12659820
SRR9045295
SRR10854653
SRR11567157
SRR11061153
SRR9045291
SRR9045297
SRR11567261
SRR11567260
SRR9045296
SRR9045298
SRR11061154
SRR9045294
SRR9045292
SRR9045293
Length
GC-content
Samples
150,000,000
QC Reads
0
100,000
Total reads mapped
0
0.25
Percentage
0
1713
Num SNVs
0
1133
Num SCVs
0
24
Num INDELs
0
Primary MAG
Reassembled MAGs
ALJR (8)
ALSA (6)
MASA (5)
MASP (5)


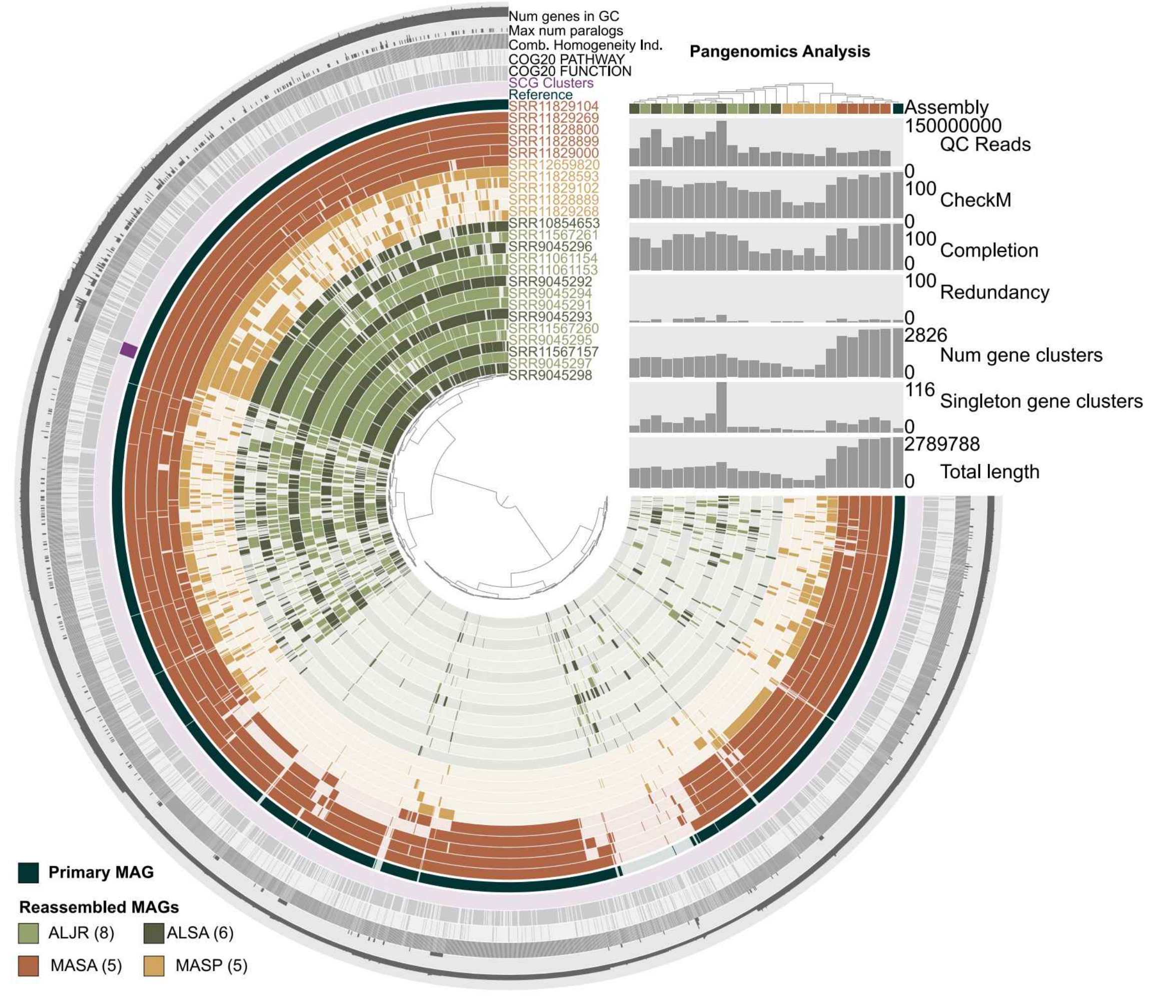

Num genes in GC
Max num paralogs
Comb. Homogeneity Ind.
COG20 PATHWAY
COG20 FUNCTION
SCG Clusters
Reference
Pangenomics Analysis
SRR11829104
SRR11829269
SRR11828800
SRR11828899
SRR11829000
SRR12659820
SRR11828593
SRR11829102
SRR11828889
SRR11829268
SRR10854653
SRR11567261
SRR9045296
SRR11061154
SRR11061153
SRR9045292
SRR9045294
SRR9045291
SRR9045293
SRR11567260
SRR9045295
SRR11567157
SRR9045297
SRR9045298
Assembly
150000000
QC Reads
0
100
CheckM
0
100
Completion
0
100
Redundancy
0
2826
Num gene clusters
0
116
Singleton gene clusters
0
2789788
Total length
0
Primary MAG
Reassembled MAGs
ALJR (8)
ALSA (6)
MASA (5)
MASP (5)

### Figure S14

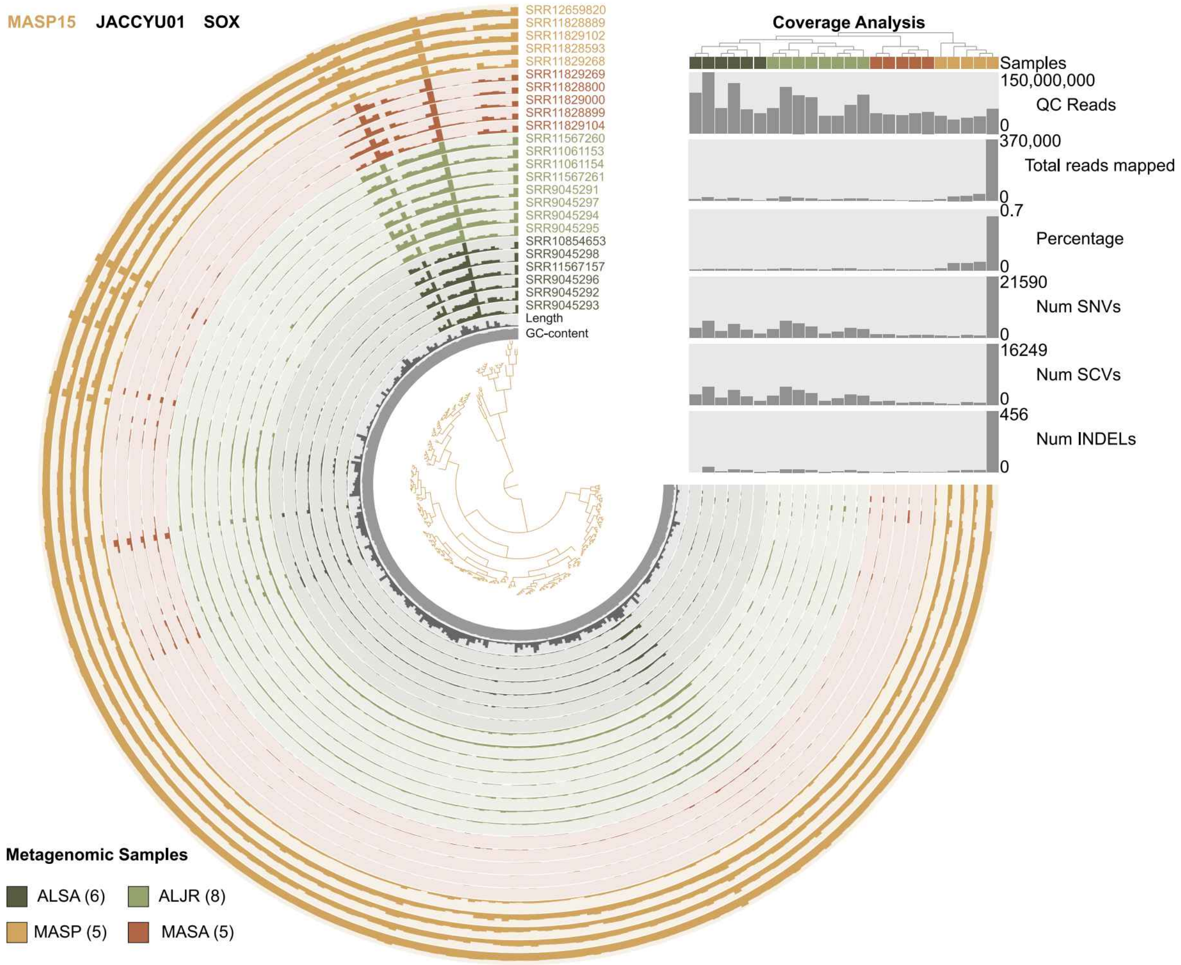
MASP15 JACCYU01 SOX
SRR12659820
SRR11828889
SRR11829102
SRR11828593
SRR11829268
SRR11829269
SRR11828800
SRR11829000
SRR11828899
SRR11829104
SRR11567260
SRR11061153
SRR11061154
SRR11567261
SRR9045291
SRR9045297
SRR9045294
SRR9045295
SRR10854653
SRR9045298
SRR11567157
SRR9045296
SRR9045292
SRR9045293
Length
GC-content
Coverage Analysis
Samples
150,000,000
QC Reads
0
370,000
Total reads mapped
0
0.7
Percentage
0
21590
Num SNVs
0
16249
Num SCVs
0
456
Num INDELs
0
Metagenomic Samples
ALSA (6)
ALJR (8)
MASP (5)
MASA (5)

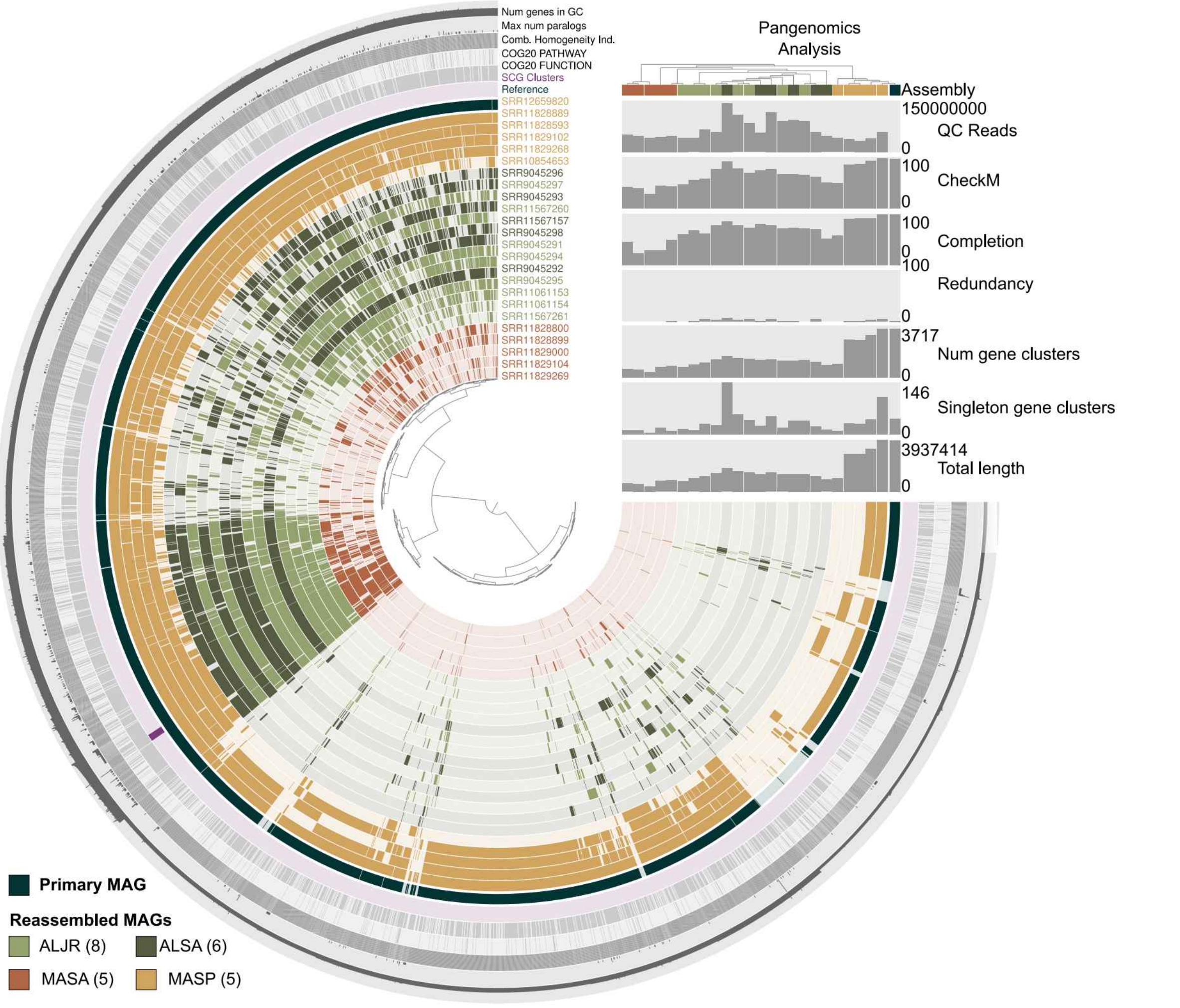
Num genes in GC
Max num paralogs
Comb. Homogeneity Ind.
COG20 PATHWAY
COG20 FUNCTION
SCG Clusters
Reference
SRR12659820
SRR11828889
SRR11828593
SRR11829102
SRR11829268
SRR10854653
SRR9045296
SRR9045297
SRR9045293
SRR11567260
SRR11567157
SRR9045298
SRR9045291
SRR9045294
SRR9045292
SRR9045295
SRR11061153
SRR11061154
SRR11567261
SRR11828800
SRR11828899
SRR11829000
SRR11829104
SRR11829269
Pangenomics Analysis
Assembly
150000000
QC Reads
0
100
CheckM
0
100
Completion
0
100
Redundancy
0
3717
Num gene clusters
0
146
Singleton gene clusters
0
3937414
Total length
0
Primary MAG
Reassembled MAGs
ALJR (8)
ALSA (6)
MASA (5)
MASP (5)

### Figure S15

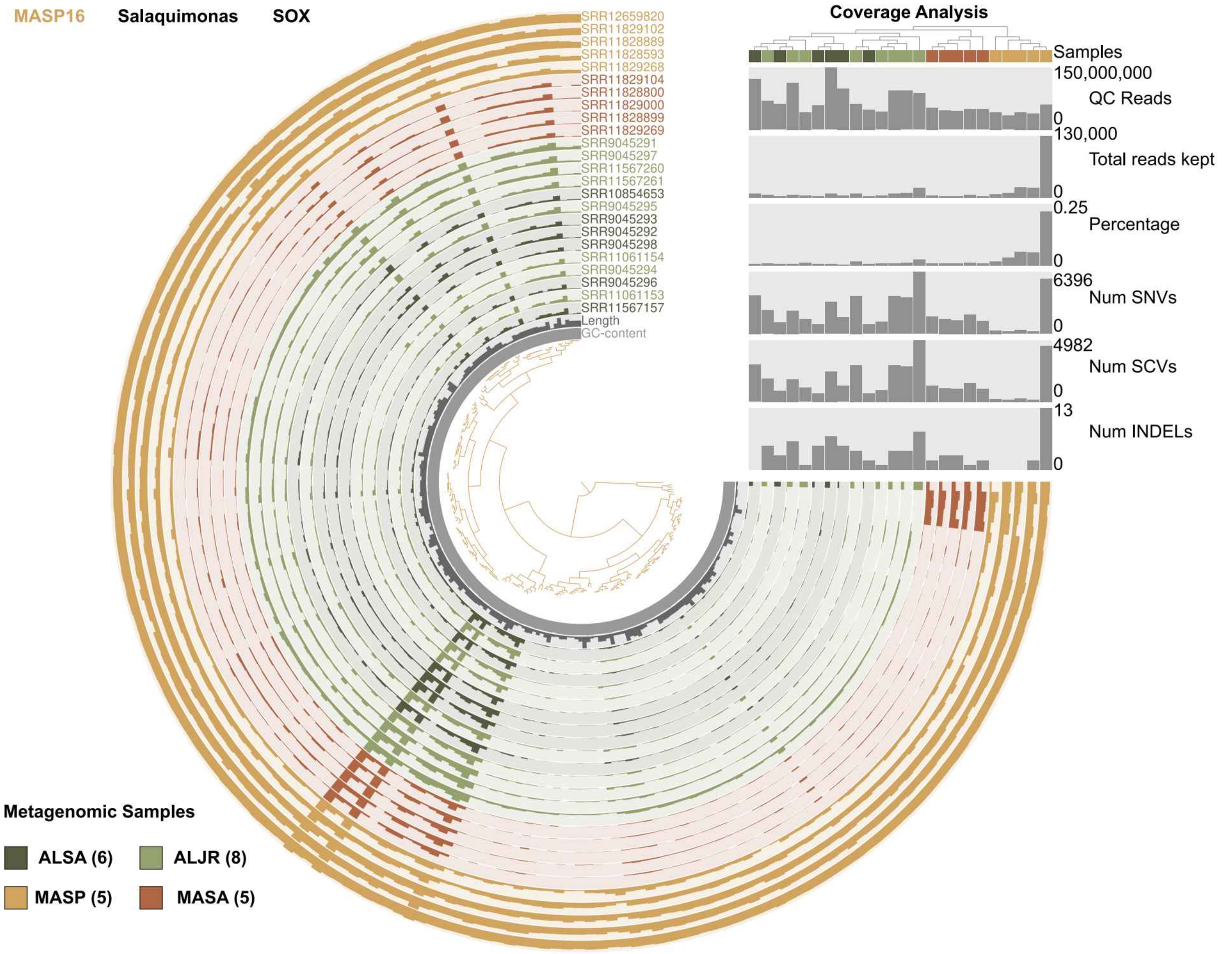
MASP16
Salaquimonas
SOX
Coverage Analysis
Samples
150,000,000
QC Reads
0
130,000
Total reads kept
0
0.25
Percentage
0
6396
Num SNVs
0
4982
Num SCVs
0
13
Num INDELs
0
SRR12659820
SRR11829102
SRR11828889
SRR11828593
SRR11829268
SRR11829104
SRR11828800
SRR11829000
SRR11828899
SRR11829269
SRR9045291
SRR9045297
SRR11567260
SRR11567261
SRR10854653
SRR9045295
SRR9045293
SRR9045292
SRR9045298
SRR11061154
SRR9045294
SRR9045296
SRR11061153
SRR11567157
Length
GC-content
Metagenomic Samples
ALSA (6)
ALJR (8)
MASP (5)
MASA (5)

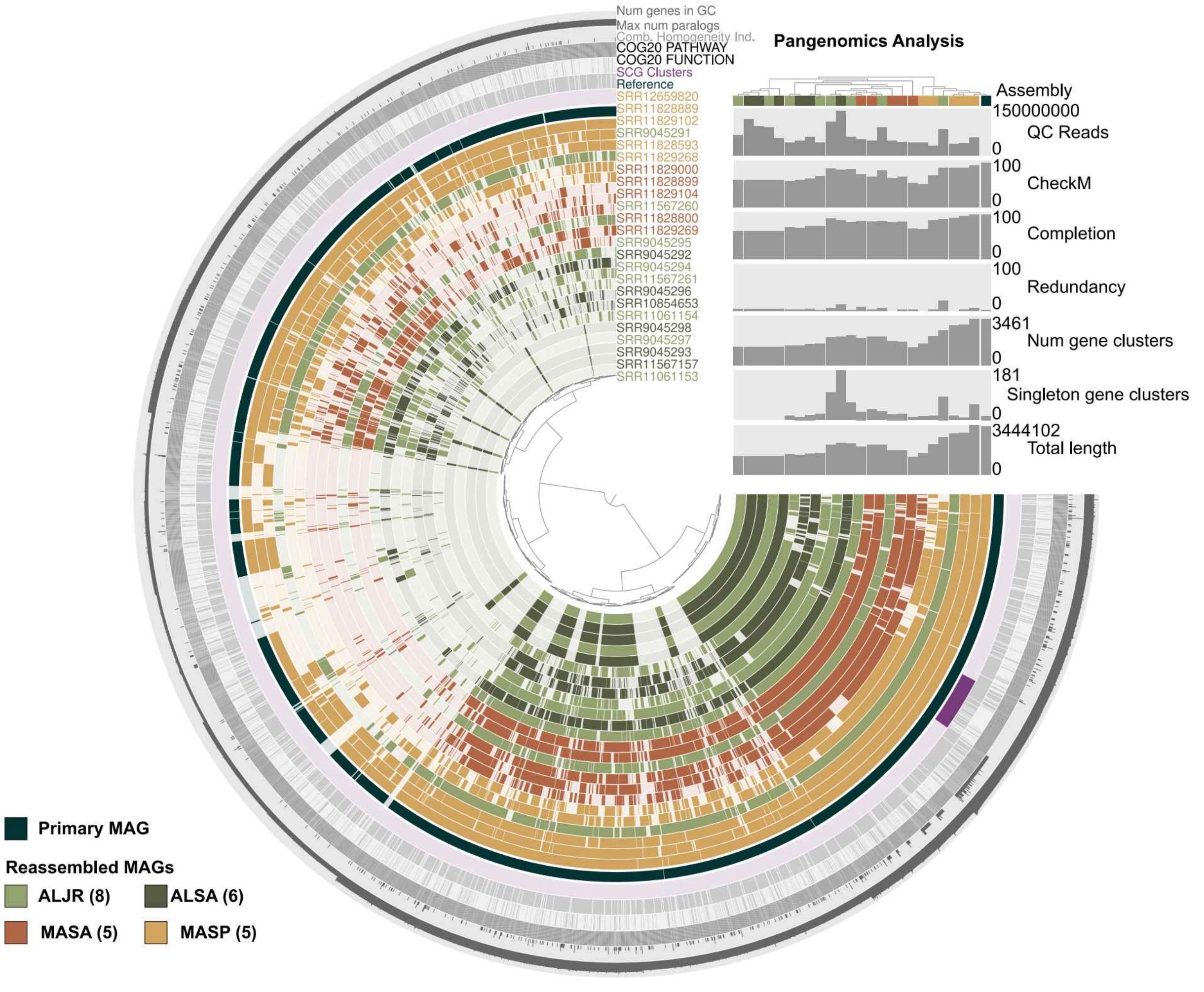
Num genes in GC
Max num paralogs
Comb. Homogeneity Ind.
COG20 PATHWAY
COG20 FUNCTION
SCG Clusters
Reference
SRR12659820
SRR11828889
SRR11829102
SRR9045291
SRR11828593
SRR11829268
SRR11829000
SRR11828899
SRR11829104
SRR11567260
SRR11828800
SRR11829269
SRR9045295
SRR9045292
SRR9045294
SRR11567261
SRR9045296
SRR10854653
SRR11061154
SRR9045298
SRR9045297
SRR9045293
SRR11567157
SRR11061153
Pangenomics Analysis
Assembly
150000000
QC Reads
0
100
CheckM
0
100
Completion
0
100
Redundancy
0
3461
Num gene clusters
0
181
Singleton gene clusters
0
3444102
Total length
0
Primary MAG
Reassembled MAGs
ALJR (8)
ALSA (6)
MASA (5)
MASP (5)

**Figure S16.** A) Coverage (left) and gene variability (right) of selected genes. In the heatmaps, each row represents a gen (see Supplementary Data 7 for the full list of genes), and each column is a metagenomic sample. Genes with average coverage under 5 were excluded from the analysis and are indicated in gray. Gene variability was calculated as average gene entropy from single codon variants (SCVs) and results are summarized in Supplemental Data 7. B) Comparison of coverage and entropy values for the two closely-related MAGs ALJR36 and MASA10.

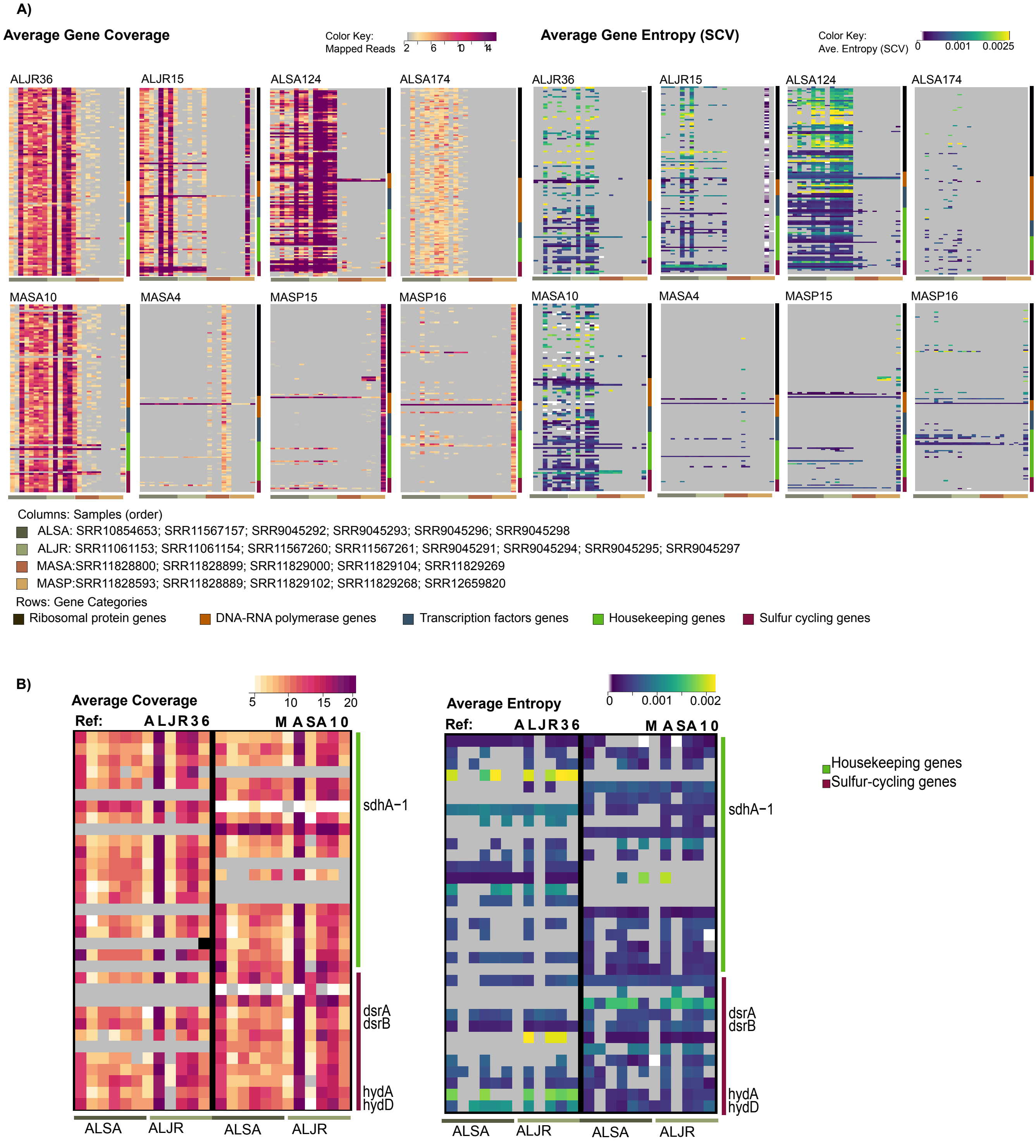

**Figure S17.** Squeme of the Reverse Citrate Cycle (Arnon-Buchanan) and Phosphate acetyltransferase-acetate kinase pathways. The enzymes detected in the reference-guided reassembled metagenomes corresponding to the group ALJR36 are indicated in dark grey and those in the MASA10 reassembled group are in light grey. Adapted from Kegg map (00720) representing the Carbon Fixation Pathways in Prokaryotes.

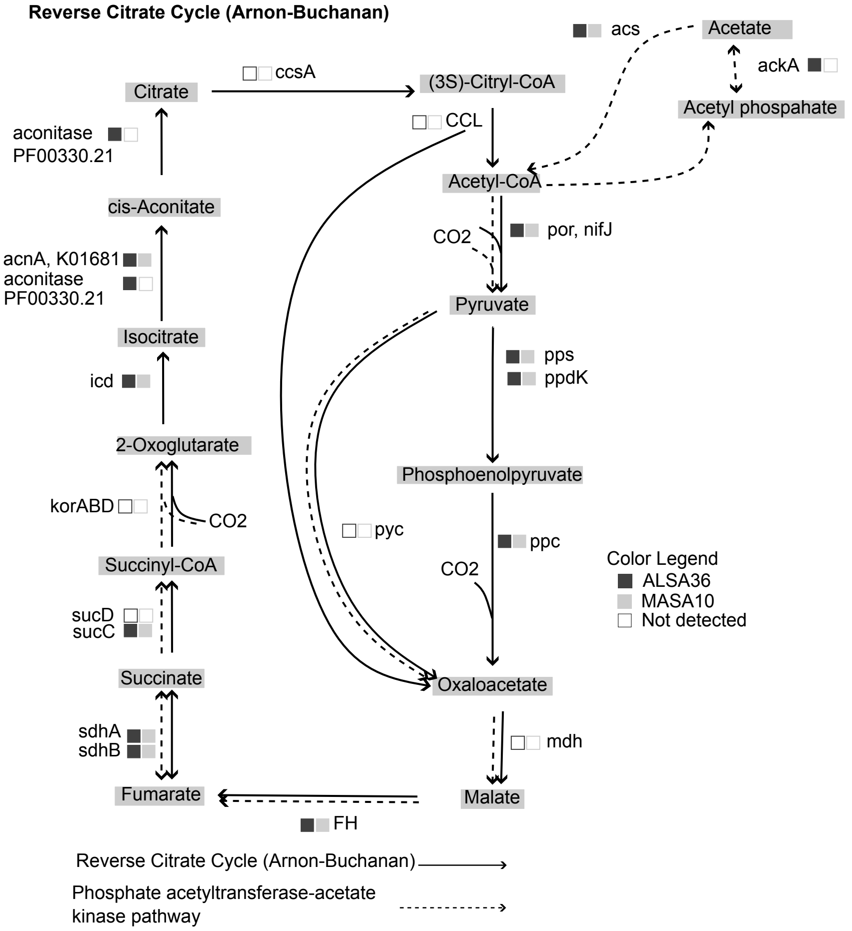
